# Identification of Unusual Oxysterols Biosynthesised in Human Pregnancy by Charge-Tagging and Liquid Chromatography - Mass Spectrometry

**DOI:** 10.1101/2022.02.07.478301

**Authors:** Alison L. Dickson, Eylan Yutuc, Catherine A. Thornton, Yuqin Wang, William J. Griffiths

## Abstract

The aim of this study was to identify sterols, oxysterols and any down-stream metabolites in placenta, umbilical cord blood plasma, maternal plasma and amniotic fluid to enhance our knowledge of the involvement of these molecules in pregnancy. We confirm the identification of 20S- hydroxycholesterol in human placenta, previously reported in a single publication, and propose a pathway from 22R-hydroxycholesterol to a C_27_ bile acid of probable structure 3β,20R,22R- trihydroxycholest-5-en-(25R)26-oic acid. The pathway is evident not only in placenta, but pathway intermediates are also found in umbilical cord plasma, maternal plasma and amniotic fluid but not non-pregnant women.

## 1. Introduction

Amongst other functions, the placenta plays a key role in the transport of cholesterol from the mother to the fetus (1). The placenta is rich in cholesterol metabolising enzymes, particularly those involved in progesterone and estrogen synthesis (2). Hence, it should also be a site for oxysterol synthesis and further metabolism. Cytochrome P450 (CYP) 11A1 (also known as P450_SCC_) is the enzyme that generates pregnenolone from cholesterol via consecutive 22R- and 20R-hydroxylations followed by side-chain cleavage (3). Progesterone is then formed from pregnenolone by oxidation at C-3 and Δ^5^ – Δ^4^ isomerisation by hydroxysteroid dehydrogenase (HSD) 3B1. Although 22R-hydroxycholesterol (22R- HC) and 20R,22R-dihydroxycholesterol (20R,22R-diHC) are known intermediates in CYP11A1- mediated side-chain cleavage of cholesterol to give pregnenolone (4), few studies have explored the oxysterol profile of placenta (5, 6). One important, but until now not replicated, finding made in the early 2000’s was the presence of 20S-hydroxycholesterol (20S-HC) in human placenta (7). 20S-HC is an enigmatic oxysterol with many biological properties, but seldom reported in mammalian systems (8). Similar to the situation with placenta, there are few reports of the oxysterol profiles of umbilical cord blood (5), i.e. blood of fetal origin that remains in the placenta and in the attached umbilical cord after childbirth, or of amniotic fluid, the fluid that acts as a cushion for the growing fetus and serves to facilitate the exchange of biochemicals between mother and fetus. Interestingly, however, oxysterols have been found as their sulphate esters in meconium, including 20,22-diHC, 22-HC, 23- hydroxycholesterol (23-HC) and 24-hydroxycholesterol (24-HC) (9, 10).

Here we report liquid chromatography (LC) – mass spectrometry (MS)-based methods for the identification of oxysterols in full term placenta, plasma derived from cord blood and pregnant female blood (maternal blood) and in mid-gestation amniotic fluid. The methods are based on high mass resolution MS with multistage fragmentation (MS^n^) exploiting charge-tagging to enhance analyte signal (Supplemental Figure S1A).

## 2. Materials and Methods

### 2.1. Human Material

Maternal blood was taken 24 - 48 hr prior to elective caesarean section at 37+ weeks of gestation for reasons that did not include maternal or fetal anomaly. Umbilical cord blood was collected at delivery of the baby. Control plasma was from non-pregnant females. Amniotic fluid was obtained at 16 – 18 weeks of pregnancy during diagnostic amniocentesis; only samples with no fetal chromosomal abnormality were used. All samples were collected with approval from an appropriate Health Research Authority Research Ethics Committee. All participants provided informed consent and the study adhered to the principles of the Declaration of Helsinki.

### 2.2. Sterol and Oxysterol Standards

Isotope-labelled standards [25,26,26,26,27,27,27-^2^H_7_]24R/S-hydroxycholesterol ([^2^H_7_]24R/S-HC), [25,26,26,26,27,27,27-^2^H_7_]22R-hydroxycholesterol ([^2^H_7_]22R-HC), [25,26,26,26,27,27,27-^2^H_7_]22S-hydroxycholesterol ([^2^H_7_]22S-HC), [25,26,26,26,27,27,27-^2^H_7_]7α-hydroxycholesterol ([^2^H_7_]7α-HC), [26,26,26,27,27,27-^2^H_6_]7α,25-dihydroxycholesterol ([^2^H_6_]7α,25-diHC) were from Avanti Polar Lipids, Alabaster, AL. [25,26,26,26,27,27,27-^2^H_7_]20S-Hydroxycholesterol ([^2^H_7_]20S-HC) was purchased from Toronto Research Chemicals (TCI, Toronto, Canada). [^2^H_7_]22R-Hydroxycholest-4-en-3-one ([^2^H_7_]22R- HCO) was prepared from [^2^H_7_]22R-HC by treatment with cholesterol oxidase enzyme (*Streptomyces sp*., Merck, Dorset, UK) (11).

### 2.3. Sterol and Oxysterol Extraction

#### 2.3.1. Placenta

Sterols and oxysterols were extracted from placental tissue using a modified protocol previously used to extract oxysterols from brain and liver tissue (12, 13). Approximately 400 mg of tissue was cut from the maternal side of fresh placenta, weighed and washed three times in PBS to remove blood. The tissue was then transferred to a gentleMACS™ C tube (Miltenyi Biotec, Woking, UK) followed by 4.2 mL of absolute ethanol containing 50 ng of [^2^H_7_]24R/S-HC and 50 ng of [^2^H_7_]22R-HCO. The tissue was homogenised for 2 min. The homogenate was transferred to a 15 mL corning tube and sonicated for 15 min. Whilst sonicating, 1.8 mL of HPLC grade water was added dropwise to give 6 mL of homogenate at 70% ethanol. The homogenate was then centrifuged for 1 hr at 4,000 x *g.* The supernatant was transferred to a fresh 15 mL corning tube and the remaining pellet re-suspended in a further 4.2 mL ethanol containing 50 ng of [^2^H_7_]24R/S-HC and 50 ng of [^2^H_7_]22R-HCO. The suspension was vortex mixed and transferred back into the original gentleMACS™ C tube where it was then homogenised for a further 2 min. The homogenate was removed and sonicated for 15 min, then 1.8 mL of water added to give 6 mL of 70% ethanol. The supernatants from the two extractions were combined to yield 12 mL in 70% ethanol. This was mixed by vortex and sonicated for a further 10 min followed by centrifugation for 1 hr at 4,000 x *g*. Ten % (1.2 mL) of the total supernatant was added to 300 µL of 70% ethanol under sonication. The 1.5 mL of sample was subjected to solid phase extraction (SPE) by a procedure modified from an earlier protocol (12, 13) to allow the collection of C_21_ steroids besides sterols and oxysterols including sterol acids.

The sample from above was loaded onto a Certified Sep-Pak C_18_, 200 mg (3 cm^3^, Waters Inc. Elstree, UK) reversed-phase SPE column previously conditioned with ethanol (4 mL) followed by 70% ethanol (6 mL). The sample flow-through (1.5 mL) was combined with a column wash of 70% ethanol (5.5 mL) resulting in SPE1-Fr1 (7 mL) which contained oxysterols, sterol acids and C_21_ steroids. A second fraction was obtained by further washing with 70% ethanol (4 mL) and collected as SPE1-Fr2. Cholesterol and other sterols of similar hydrophobicity were eluted from the SPE column with absolute ethanol (2 mL) to give SPE1-Fr3. A final fourth fraction was eluted with a further 2 mL of absolute ethanol (SPE1-Fr4). Each of the four fractions was divided equally into A and B sub-fractions and dried overnight under vacuum by centrifugal evaporation (ScanLaf ScanSpeed vacuum concentrator, Lynge, Denmark).

Each of the lyophilised samples were reconstituted in propan-2-ol (100 µL) and mixed thoroughly by vortex. To fractions (A), 50 mM K_2_HPO_4_ buffer, pH 7 (1 mL) containing cholesterol oxidase solution (3.0 µL, 2 mg/mL in water, 44 units/mg of protein) was added. The samples were mixed by vortex and incubated at 37°C for 1 hr in a water bath. The reaction was then quenched by the addition of methanol (2 mL). Fractions (B) were treated in parallel in an identical fashion to fractions (A) but in the absence of cholesterol oxidase. Glacial acetic acid (150 µL) was added to fractions (A) and (B) and mixed by vortex. [^2^H_5_]Girard P (GP) reagent (11) (190 mg, bromide salt) was added to fractions (A) and [^2^H_0_]GP reagent (150 mg, chloride salt, TCI Europe, Oxford UK) was added to fractions (B). The samples were mixed by vortex until the derivatising reagent had dissolved. The reaction was left to proceed overnight at room temperature protected from light.

An OASIS HLB 60 mg (3 cm^3^) SPE cartridge was washed with methanol (6 mL), 5% methanol (6 mL) and conditioned with 70% methanol (4 mL). Sample from above (3.25 mL, 69% organic) was loaded onto the column and the flow-through collected. The sample tube was washed with 70% methanol (1 mL) which was then loaded onto the SPE column and the eluent combined with the earlier flow-through. The column was re-conditioned with 35% methanol (1 mL) and the eluent combined with the earlier collection. The total eluent (∼5 mL) was diluted with 4 mL of water to give ∼9 mL of 35% methanol. The 9 mL sample solution was loaded onto the column and the flow-through collected. The sorbent was re-conditioned with 17.5% methanol (1 mL) and the eluent combined with earlier flow-through. To the combined 10 mL, water (9 mL) was added to give a 19 mL of 17.5% methanol. This solution was loaded onto the column and the flow-through collected once more. The sorbent was re-conditioned with 8.75% methanol (1 mL) and the flow-through combined with the earlier collection. The total combined eluent of 20 mL was diluted with 19 mL of water to give 39 mL 8.75% methanol. The solution was loaded onto the column and the flow-through discarded. A 5% methanol solution (6 mL) was used to wash the column before the analytes were eluted. The samples were eluted into four separate 1.5 mL microcentrifuge tubes using 3 x 1 mL methanol followed by 1 mL ethanol to give SPE2-FR1, - Fr2, -Fr3, -Fr4. Oxysterols originating from SPE1-Fr1 elute across SPE2-Fr1 and SPE2-Fr2, cholesterol originating from SPE1-Fr3 elutes across SPE2-Fr1,-Fr2, Fr-3. Here we report data only for oxysterols and more polar metabolites.

Immediately prior to LC-MS analysis of oxysterols, equal volumes of SPE2-Fr1A and SPE2-Fr2A were combined with equal aliquots of SPE2-Fr1B and SPE2-Fr2B and diluted with water to form a solvent composition of 60% methanol.

#### 2.3.2. Plasma

The extraction protocol for sterols and oxysterols was essentially that described previously (11, 13, 14), with minor modification to allow for extraction of C_21_ steroids. 100 µL of plasma was added dropwise to a solution of acetonitrile (1.05 mL) containing 20 ng of [^2^H_7_]24R/S-HC and 20 ng of [^2^H_7_]22R-HCO in a ultrasonic bath with sonication. After a further 5 min of sonication, 350 µL of water was added. The sample (1.5 mL), now in 70% acetonitrile, was sonicated for a further 5 minutes and centrifuged at 17,000 x *g* at 4°C for 30 min. The sample was subjected to SPE and prepared for LC-MS analysis exactly as for the placental extract with the following modification: SPE1, Certified Sep-Pak C_18_, 200 mg, was conditioned with 70% acetonitrile rather than 70% ethanol.

#### 2.3.3. Amniotic Fluid

The protocol for extraction of sterols and oxysterols from amniotic fluid was exactly as that described for plasma except the internal standards were 7 ng of [^2^H_7_]24R/S-HC and 7 ng of [^2^H_7_]22R-HCO.

### 2.4. LC-MS(MS^n^)

Analysis was performed on a Dionex Ultimate 3000 UHPLC system (Dionex, now Thermo Fisher Scientific, Hemel Hempstead, UK) interfaced via an electrospray ionisation (ESI) probe to an Orbitrap Elite MS (Thermo Fisher Scientific). Chromatographic separation was carried out on a Hypersil Gold reversed phase C_18_ column (1.9 µm particle size, 50 x 2.1 mm, Thermo Fisher Scientific, UK). Details of the mobile phase and gradients employed are given in Supplemental Materials and Methods. MS analysis on the Orbitrap Elite was performed in the positive-ion mode with five scan events, one high resolution (120,000 full width at half maximum height at *m/z* 400) scan over the *m/z* range 400 – 610 in the Orbitrap and four MS^3^ scans performed in parallel in the linear ion trap (LIT). Mass accuracy in the Orbitrap was typically < 5 ppm. More details of the scan events are provided in Supplemental Materials and Methods. Injection volumes were 35 µL for plasma extracts and at 90 µL for amniotic fluid and placental extracts.

## 3. Results

The aim of this study was to identify oxysterols and any down-stream metabolites in placenta, cord plasma, maternal plasma and amniotic fluid to enhance our knowledge of the involvement of these molecules in pregnancy. For side-chain oxysterols expected to be present quantitative measurements were possible by the inclusion of isotope-labelled standards in the sample preparation protocol, for unexpected metabolites only semi-quantitative data was obtained, however, this could be used for relative quantification between sample sets (Table 1).

**Table 1.**
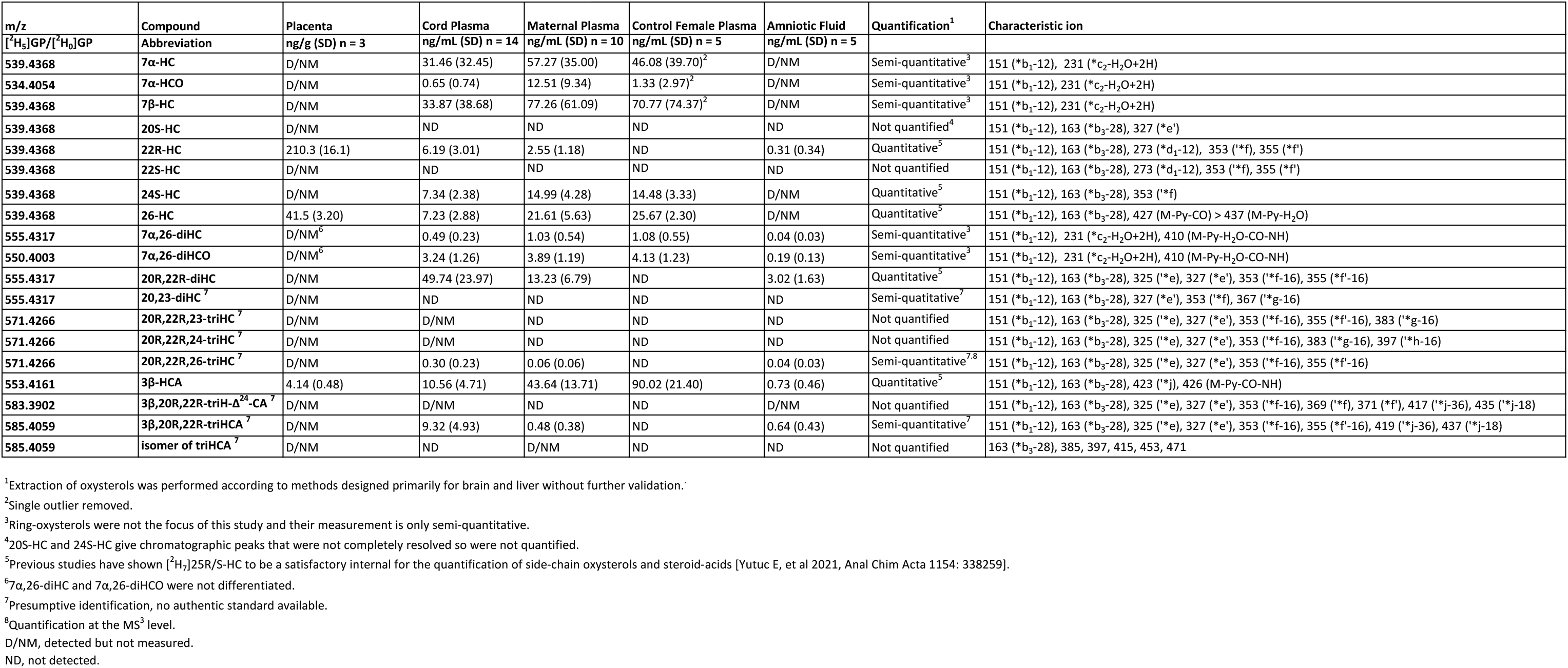
Oxysterols in placenta, cord plasma, maternal plasma and amniotic fluid.

### 3.1. Identification of Oxysterols in Placenta

The placenta is a blood-rich organ. The maternal side contains less vascular tissue than the fetal side and was selected for analysis. During sample preparation tissue was washed three times with PBS to remove blood.

#### 3.1.1. Monohydroxycholesterols:- 20S-HC, 22R-HC, 22S-HC, 24S-HC, (25R)26-HC, 7α-HC and 7β-HC

Shown in Figure 1A (upper panel) is the LC-MS reconstructed ion chromatogram (RIC) for monohydroxycholesterols (HC, *m/z* 539.4368 ± 5ppm) found in placenta following derivatisation with [^2^H_5_]GP reagent. At first glance, the chromatogram shows some similarity to that of adult plasma (Figure 1A central panel) except for the additional presence an intense pair of peaks corresponding to the *syn* and *anti* conformers of [^2^H_5_]GP-derivatised 22R-HC in the placental sample. Note *syn* and *anti* conformers are a consequence of GP-derivatisation (see Supplemental Figure S1B). 22R-HC is usually only a minor oxysterol in adult plasma/serum (14, 15) and is essentially absent in the NIST SRM 1950 plasma sample (representative of the adult population of the USA) illustrated here (16). The observation of 22R-HC in placenta is not surprising as CYP11A1, the enzyme which generates this oxysterol in the pathway from cholesterol to pregnenolone, is abundant in placenta (2, 17). The identity of the two early eluting peaks was confirmed by reference to [^2^H_7_]22R-HC authentic standard which co-elutes and gives an identical MS^3^ ([M]^+^→[M-Py]^+^→, where Py is pyridine) fragmentation pattern (Figures 1A & 1B). A major advantage of the GP-derivatisation method is that unlike the un- derivatised [M+H]^+^ or [M+NH_4_]^+^ ion, the [M]^+^ ion of the GP-derivative gives a structurally informative MS^3^ spectrum. The fragmentation of GP-derivatised oxysterols has been described in detail elsewhere (18, 19). In brief, a 3β-hydroxy-5-ene function in the parent structure, with no additional substitutions on the ring system, gives following cholesterol oxidase treatment, GP derivatisation and MS^3^, a pattern of low-*m/z* fragment ions at 151.1 (*b_1_-12), 163.1 (*b_3_-28) and 177.1 (*b_2_) and a mid-*m/z* fragment ion at 325.2 (‘*e, Supplemental Figure S1C). A characteristic, but not unique, fragment ion of 22R-HC is at *m/z* 355.3 (*f’, Figure 1B & Supplemental Figure S2A), this appears alongside a satellite peak at *m/z* 353.3 (‘*f). By generating a multiple reaction monitoring (MRM)-like chromatogram [M]^+^→[M-Py]^+^→355.3 the 22R-HC peaks in placenta are highlighted (Figure 1C upper panel). A minor unknown pair of peaks were also observed eluting much later in the MRM chromatogram. Their MS^3^ spectra suggests that these correspond to the 22S-epimer (Figure 1D). This was confirmed by analysis of [^2^H_7_]22S-HC which gave an identical MS^3^ fragmentation pattern and co-eluted with the endogenous molecule.

**Figure 1.**
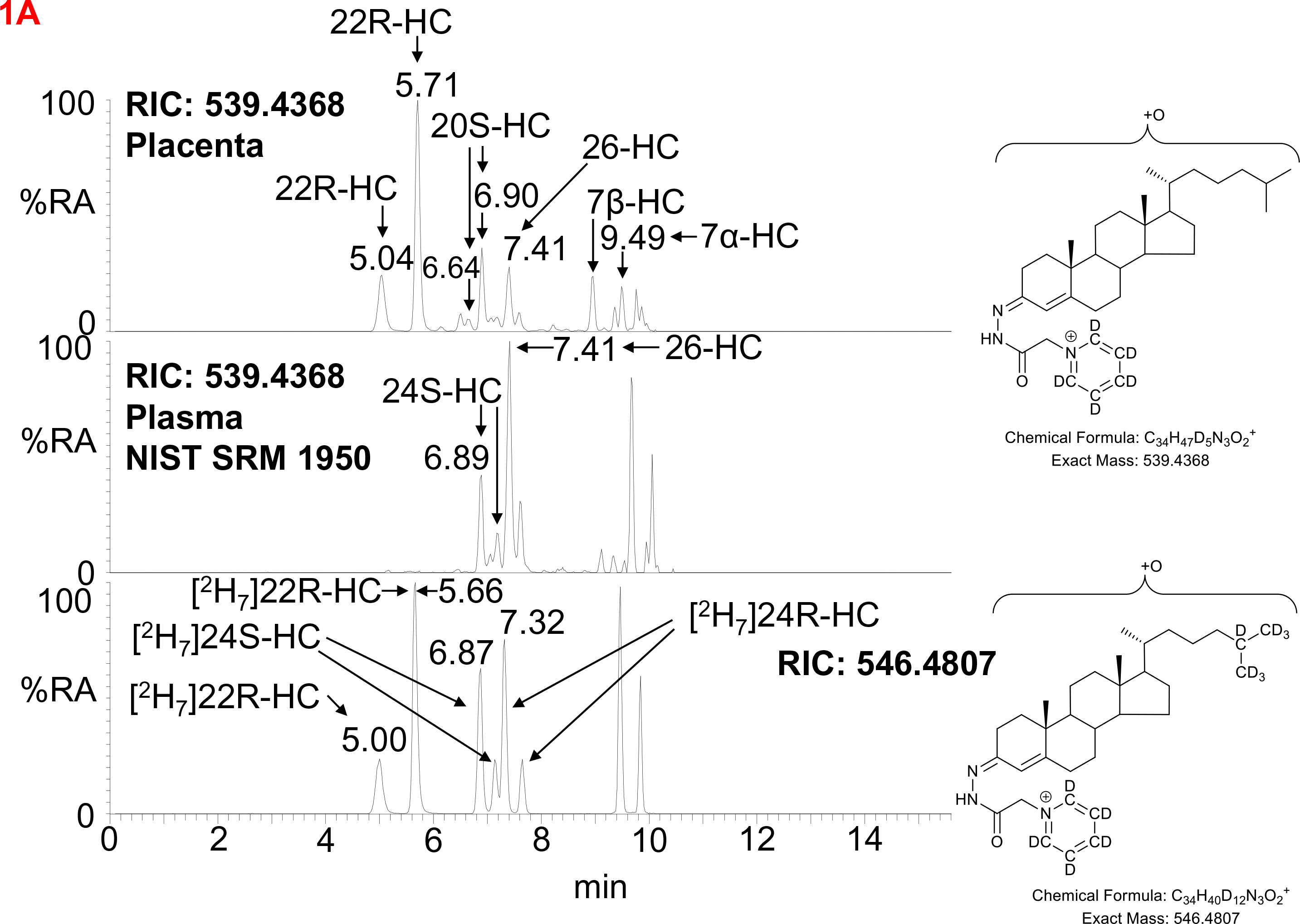
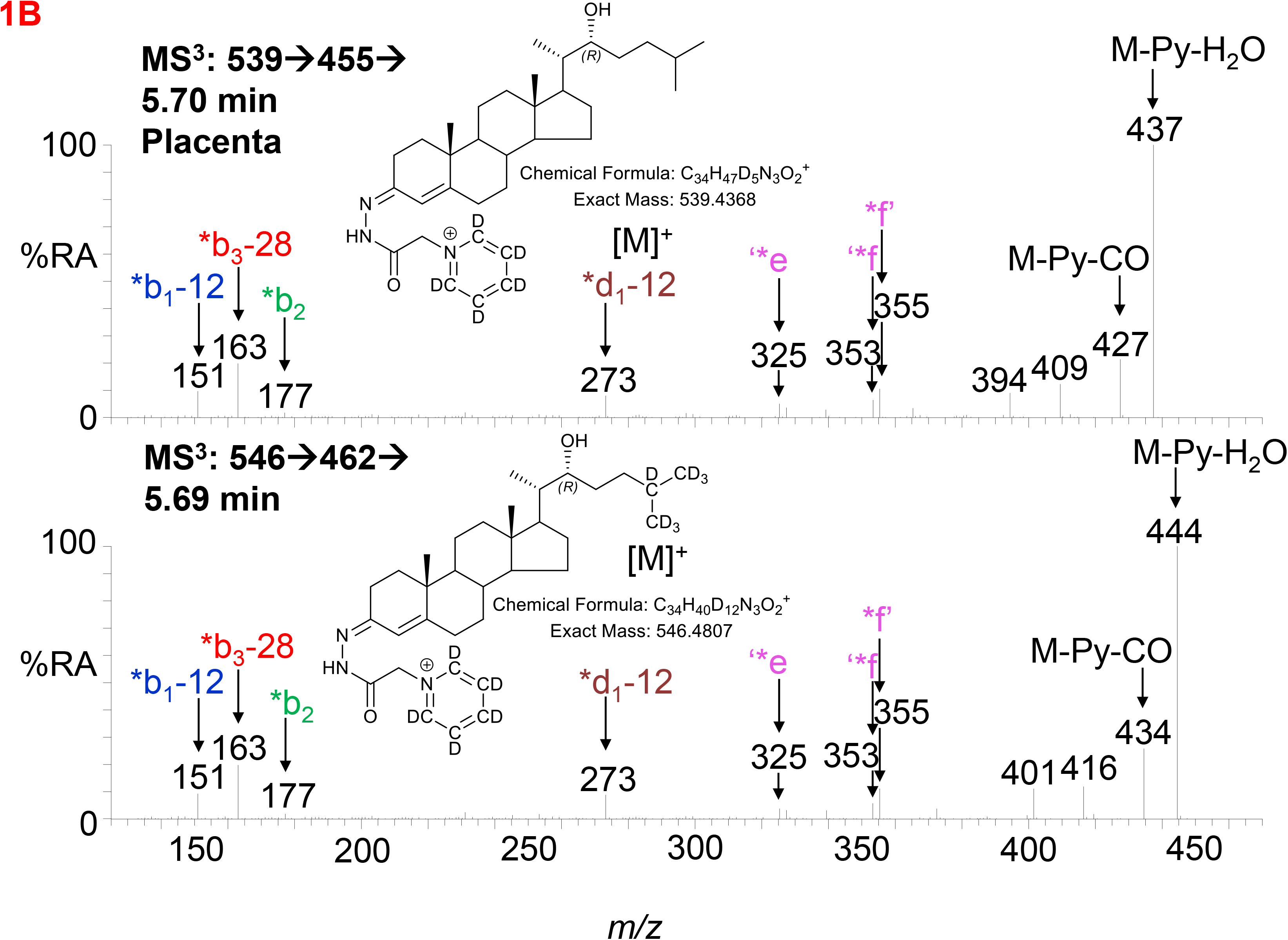
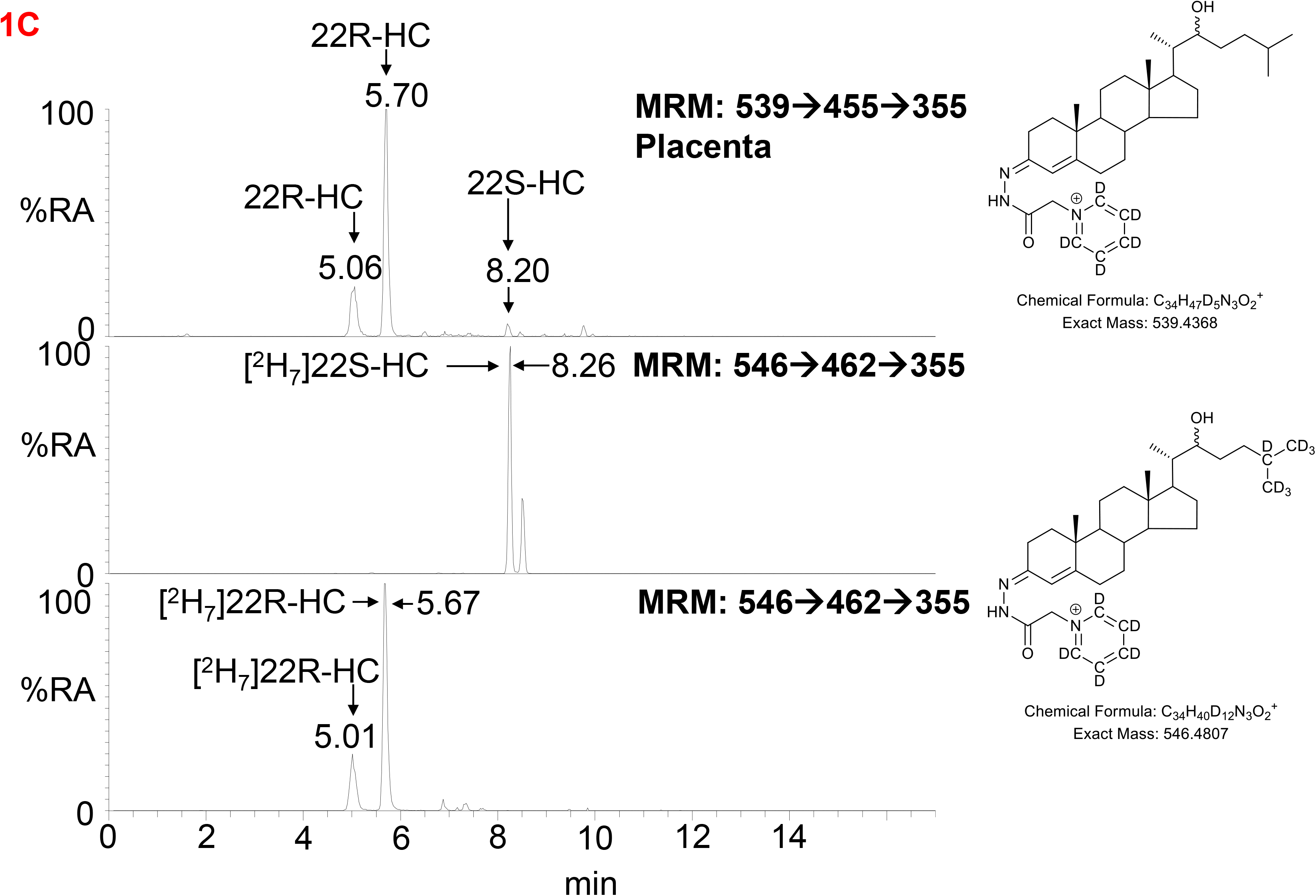
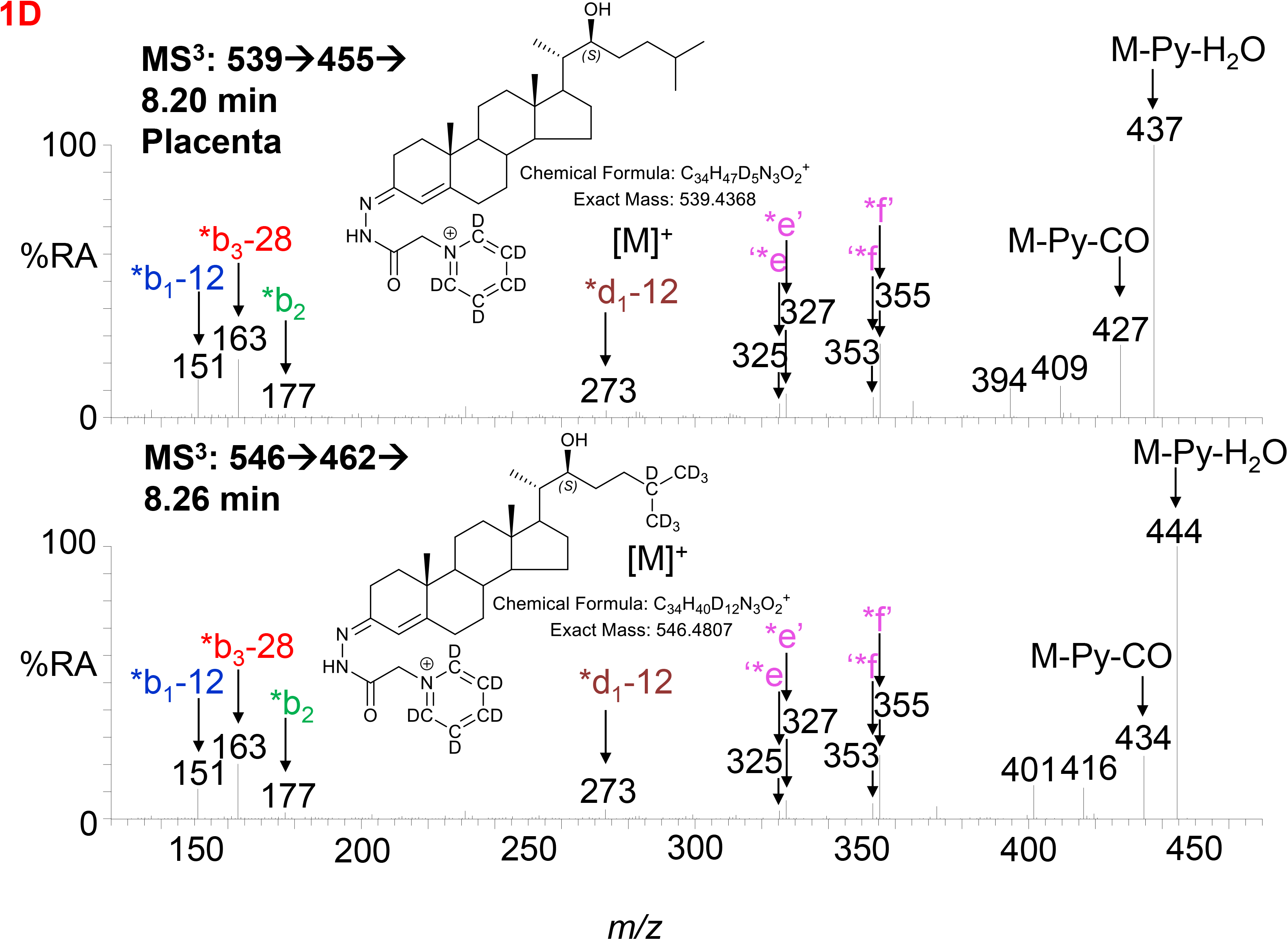
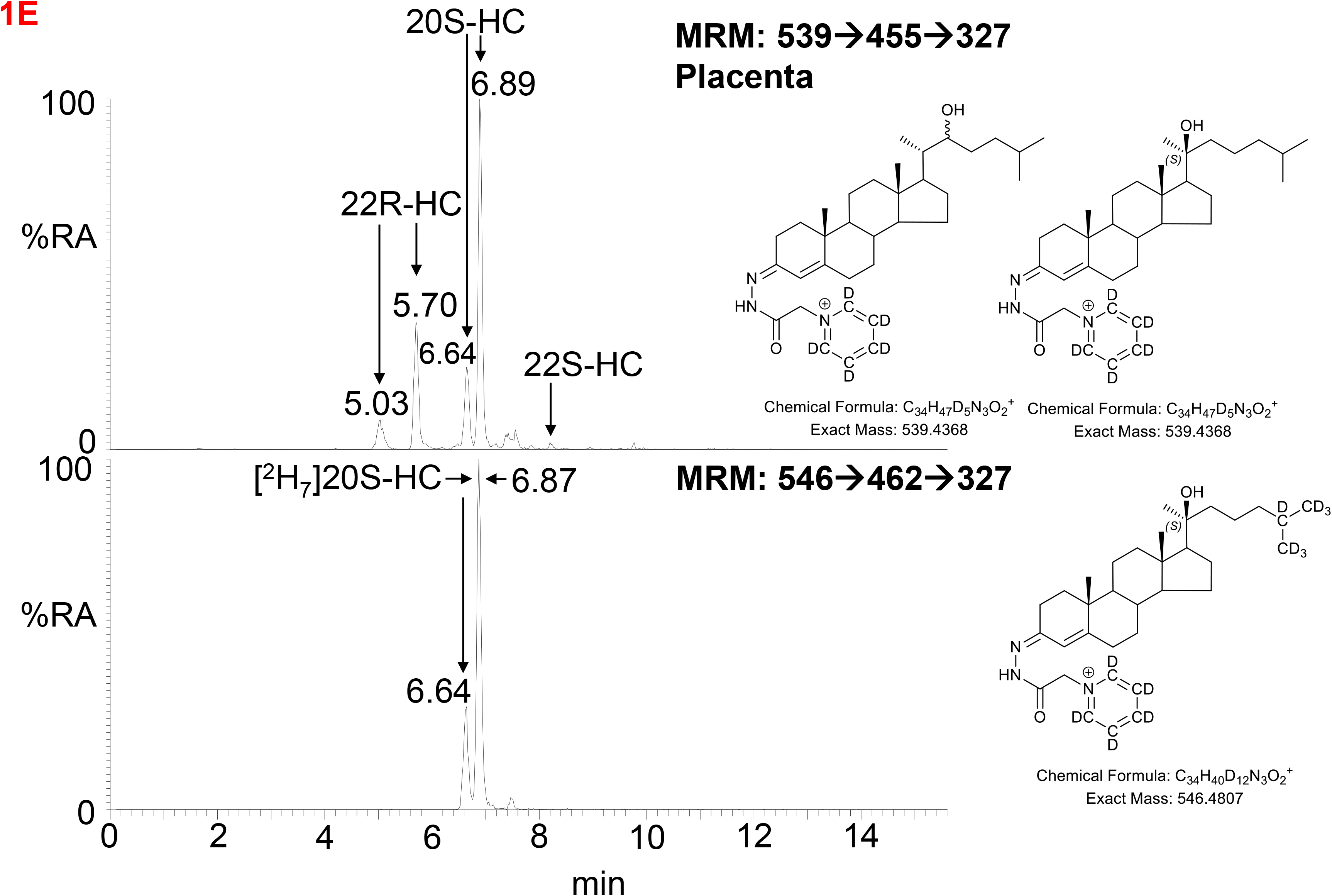
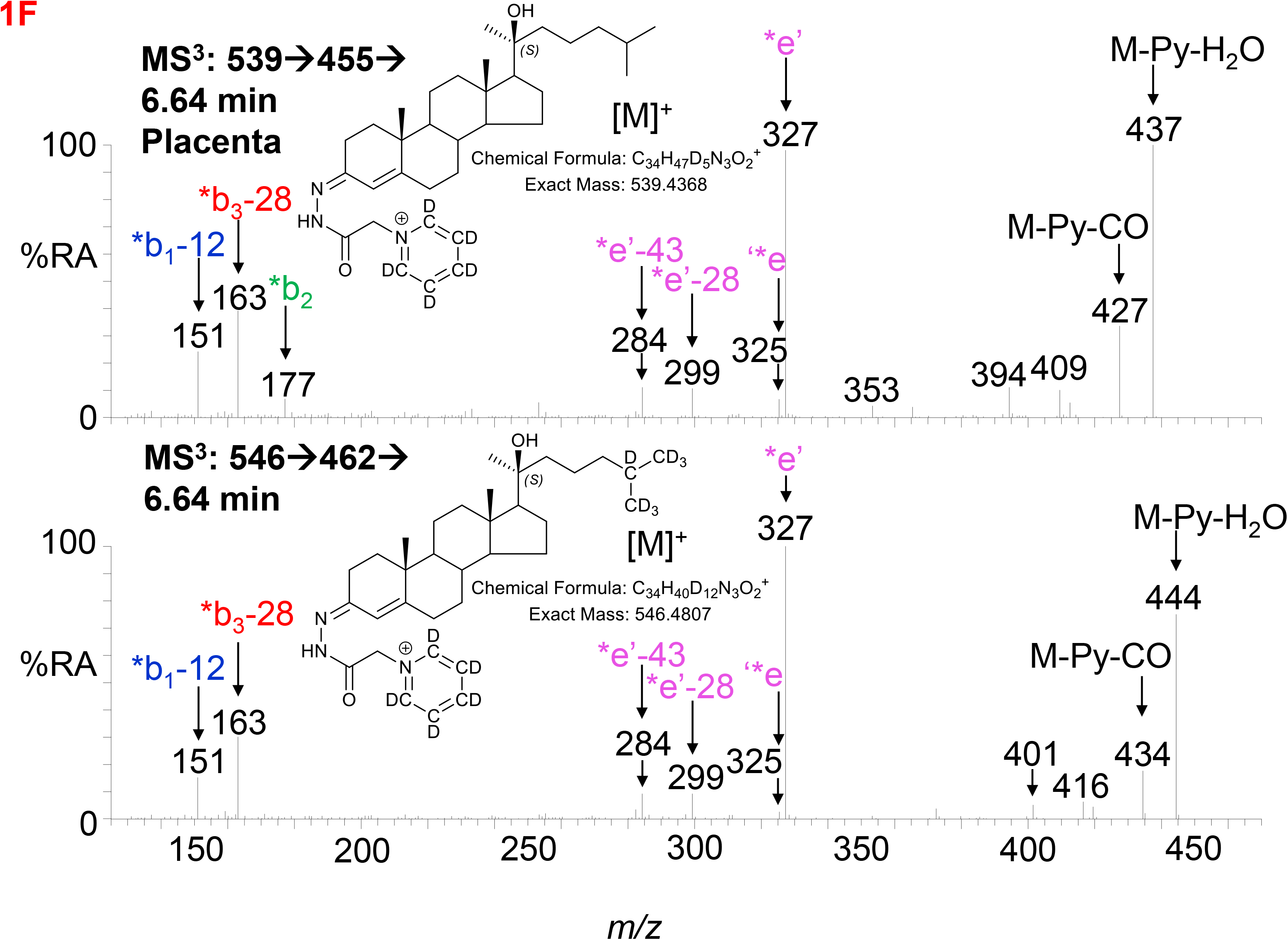
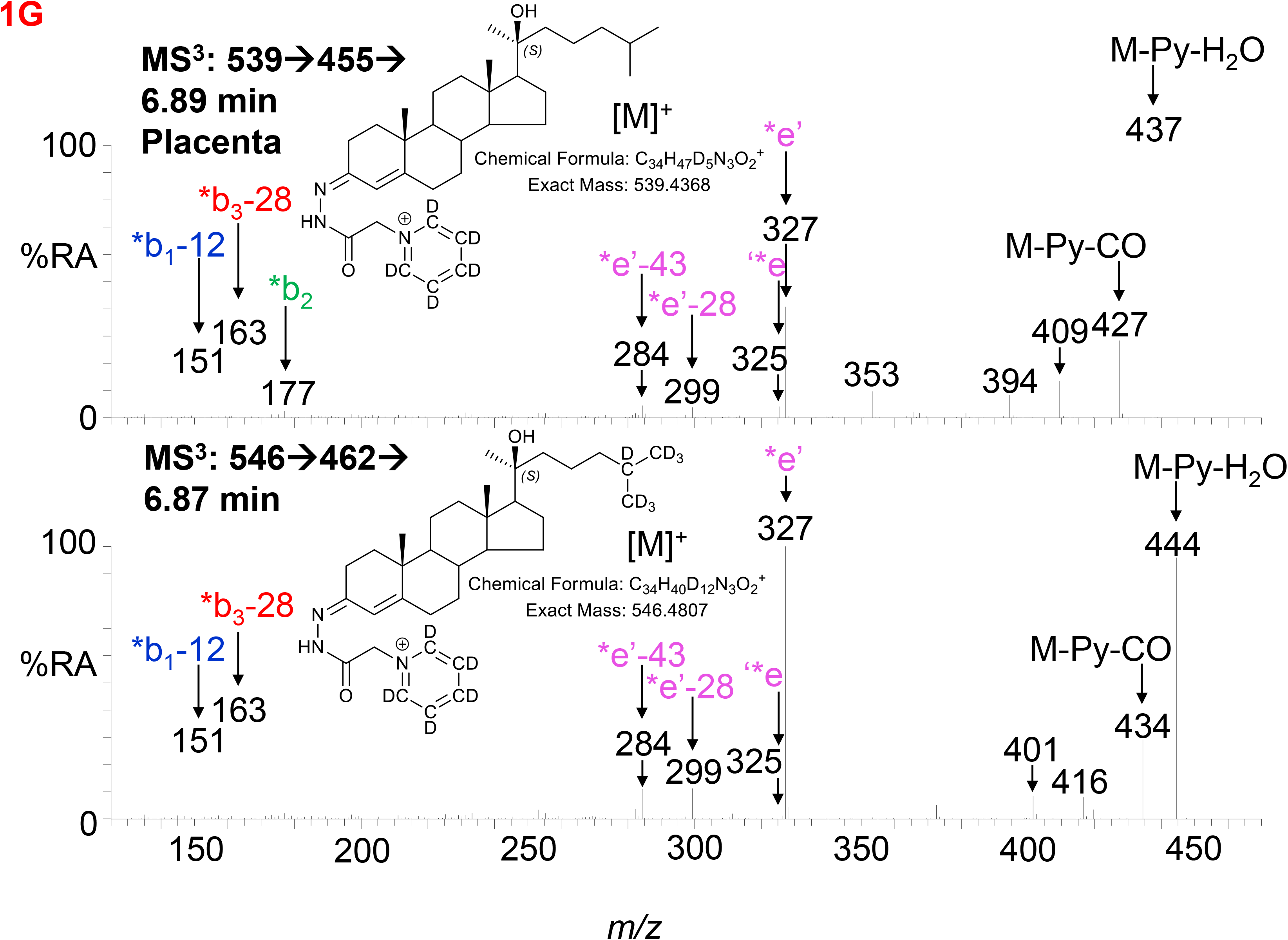
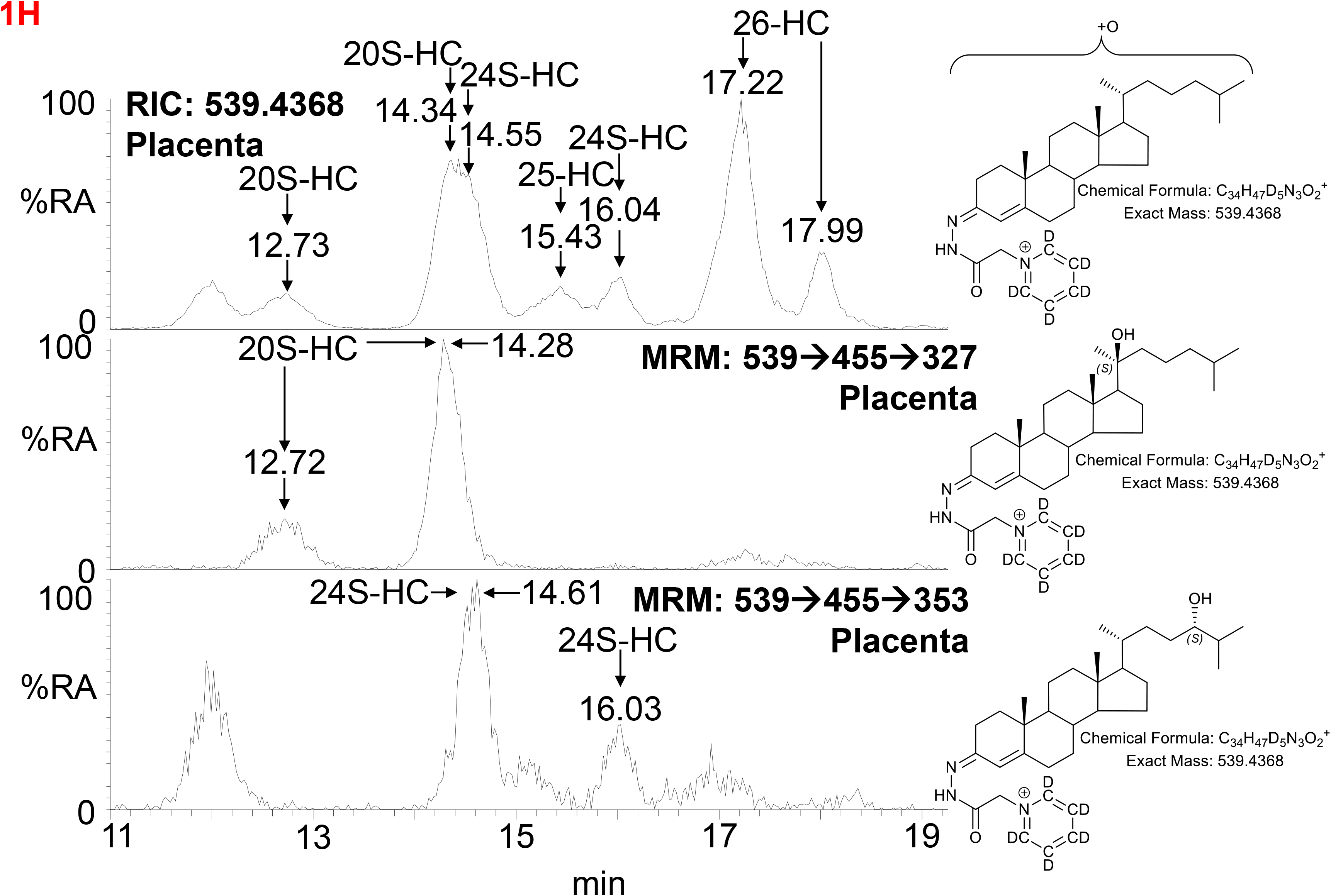
LC-MS(MS^n^) analysis of GP-derivatised monohydroxycholesterols (HC) in placenta and NIST SRM 1950 plasma. (A) Reconstructed ion chromatograms (RICs) for monohydroxycholesterols (HC, *m/z* 539.4368 ± 5 ppm) in placenta (upper panel), NIST SRM 1950 plasma (16) (central panel), and of [^2^H_7_]-labelled standards (546.4807 ± 5 ppm, lower panel). GP-derivatised oxysterols give *syn* and *anti* conformers which may or may not be resolved. (B) MS^3^ ([M]^+^→[M-Py]^+^→) spectra of 22R-HC identified in placenta (upper panel) and [^2^H_7_]22R-HC reference standard (lower panel). (C) Multiple reaction monitoring-like (MRM) chromatograms targeting 22-HC isomers ([M]^+^→[M-Py]^+^→355) found in placental (upper panel), and authentic standards of [^2^H_7_]22S-HC (central panel) and [^2^H_7_]22R-HC (lower panel). (D) MS^3^ ([M]^+^→[M-Py]^+^→) spectra of 22S-HC identified in placenta (upper panel) and [^2^H_7_]22S-HC reference standard (lower panel). (E) MRM-like chromatograms targeting 20S-HC ([M]^+^→[M-Py]^+^→327) in placenta (upper panel) and [^2^H_7_]20S-HC authentic standard (lower panel). (F & G) MS^3^ ([M]^+^→[M-Py]^+^→) spectra of *syn* and *anti* conformers of 20S-HC identified in placenta (upper panels) and of [^2^H_7_]20S-HC reference standard (lower panels). (H) 20S-HC and 24S-HC can be resolved via MRM but not by chromatography alone even when using an extended chromatographic gradient (37 min). Peaks corresponding to 20S-HC and 24S-HC coalesce in the RIC for their [M]^+^ ions (upper panel), but are resolved by their specific MRMs, i.e. 20S-HC ([M]^+^→[M-Py]→327, central panel) and 24S-HC ([M]^+^→[M-Py]^+^→353, lower panel). See Supplemental Figure S3 for MS^3^ spectra of 24S-HC. As data was collected during different sessions, 17 min gradient chromatograms (A, C & E), have been aligned to the peak corresponding to 26-HC in NIST SRM 1950 plasma.

In many LC-MS/MS studies oxysterols are identified by MRM where the transition is often non-specific (15, 20). This provides high sensitivity but relies on chromatographic separation of isomers and co-elution with isotopic labelled standards (21). In an initial interpretation of the data presented in Figure 1A (upper panel), the peak at 6.90 min was assumed to be 24S-HC, presumably from contaminating blood. However, closer scrutiny of the chromatogram and relevant MS^3^ spectra suggested that the earlier eluting peak 6.64 min was 20S-HC. 20S-HC has a characteristic MS^3^ fragmentation spectrum with a major fragment ion at 327.2 (*e’, Figure 1F & Supplemental Figure S2B). By generating a MRM chromatogram for [M]^+^→[M-Py]^+^→327.2, two peaks corresponding to the *syn* and *anti* conformers of 20S-HC are revealed (Figure 1E). The second peak co-elutes with 24S-HC, of which minor quantities are present as revealed by the 24S-HC characteristic fragment ion at *m/z* 353.3 (‘*f, Figure 1G upper panel & Supplemental Figure S1D). However, by extending the chromatographic gradient and by exploiting the specific MRM chromatograms i.e. [M]^+^→[M-Py]^+^→327.2 for 20-HC and [M]^+^→[M- Py]^+^→353.3 for 24-HC, the two isomers can be partially resolved (Figure 1H, see Supplemental Figure S3 for the MS^3^ spectrum of 24S-HC). Fortuitously, as both 20S-HC and 24S-HC give *syn* and *anti* conformers following GP-derivatisation, the first peak of 20S-HC is completely resolved from both peaks of 24S-HC and it is only the second peak of 20S-HC and the first peak of 24S-HC that partially co-elute, leaving the second peak of 24S-HC completely resolved from 20S-HC.

As is evident from Figure 1A, other monohydroxycholesterols were also observed in placenta including (25R)26-hydroxycholesterol (26-HC, also known by the non-systematic name 27-hydroxycholesterol, 27-HC) and 7α-hydroxycholesterol (7α-HC). MS^3^ spectra are shown in Supplemental Figure S3. 7β- Hydroxycholesterol (7β-HC) was also observed, this like 7α-HC may be endogenous, but can also be formed during sample preparation by *ex vivo* oxidation (22).

#### 3.1.2. Dihydroxycholesterols, Trihydroxycholesterols, Pregnenolone and Progesterone

The second step in the conversion of cholesterol to pregnenolone by CYP11A1 is the generation of 20R,22R-diHC from 22R-HC. The RIC (*m/z* 555.4317 ± 5 ppm) for dihydroxycholesterols reveals two major peaks which appear at retention times, and give identical MS^3^ ([M]^+^→[M-Py]^+^→) spectra, to the authentic standard of 20R,22R-diHC and are identified as the *syn* and *anti* conformers of the GP-derivative (Figure 2A & B). The MS^3^ fragmentation pattern bears features of both 20-HC (*e’, *m/z* 327.2, Supplemental Figure S2C) and 22-HC (*f’-16, *m/z* 355.3, Supplemental Figure S2D) and to search for possible epimers of 20R,22R-diHC, MRM-like chromatograms were constructed for the transitions [M]^+^→[M-Py]^+^→327.2 and ([M]^+^→[M-Py]^+^→355.3 (Figure 2A). A minor peak was observed at 5.04 min, but the MS^3^ fragmentation pattern corresponded to a different isomer (x,y-diHC), possibly 22,23- dihydroxycholesterol (22,23-diHC, Figures 2C upper panel, and Supplemental Figure S2E). Unsurprisingly, when a search was made for pregnenolone by constructing the appropriate RIC (*m/z* 450.3115) this steroid was evident as was progesterone (*m/z* 448.2959), its HSD3B1 oxidised metabolite, in the fraction not treated with cholesterol oxidase i.e. fraction B (Figure 2D & 2E). Besides 20R,22R-diHC and the presumptively identified 22,23-diHC, low levels of 7α,(25R)26- dihydroxycholesterol (7α,(25R)26-diHC) were also found in placenta (Figure 2A & 2C).

**Figure 2.**
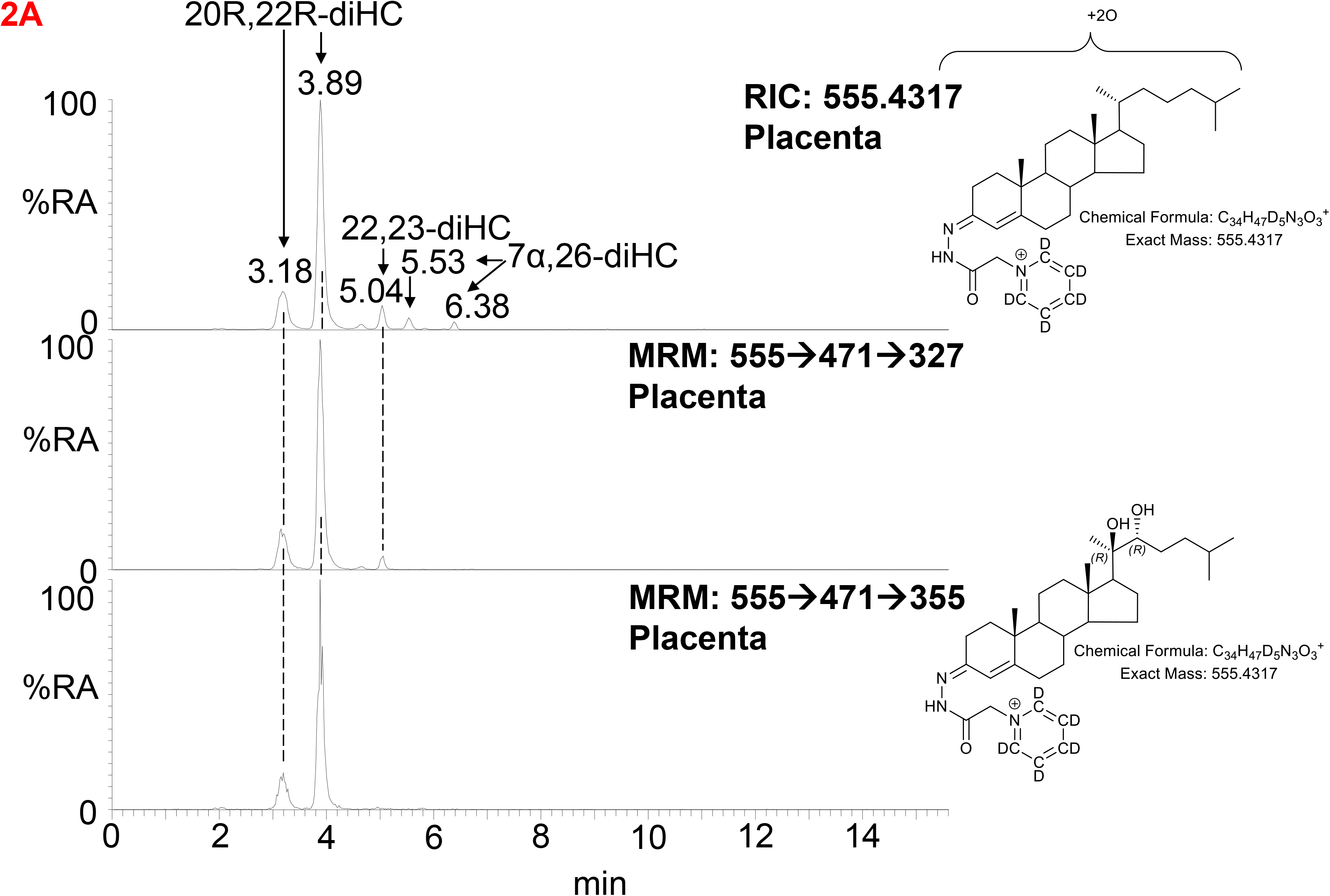
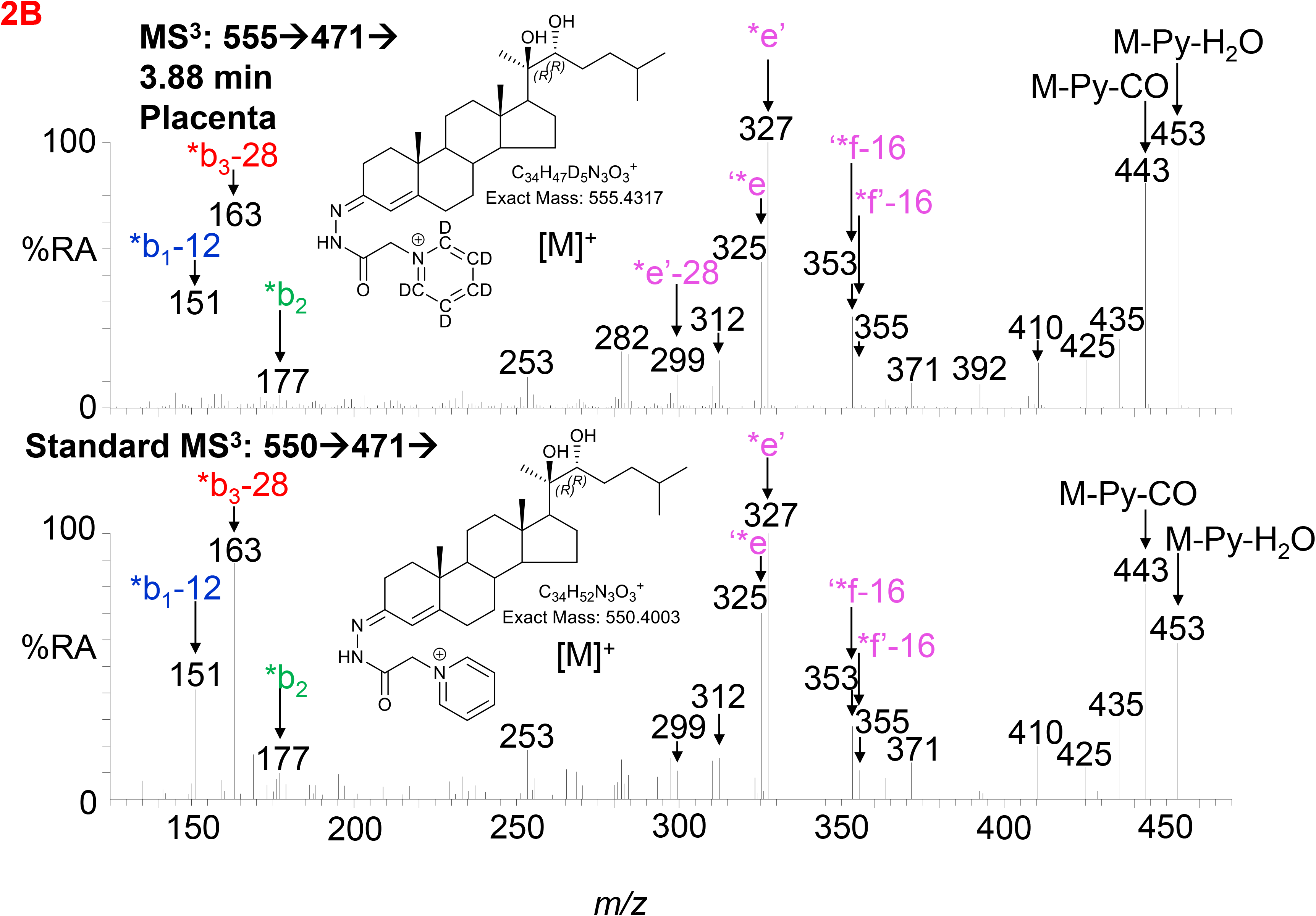
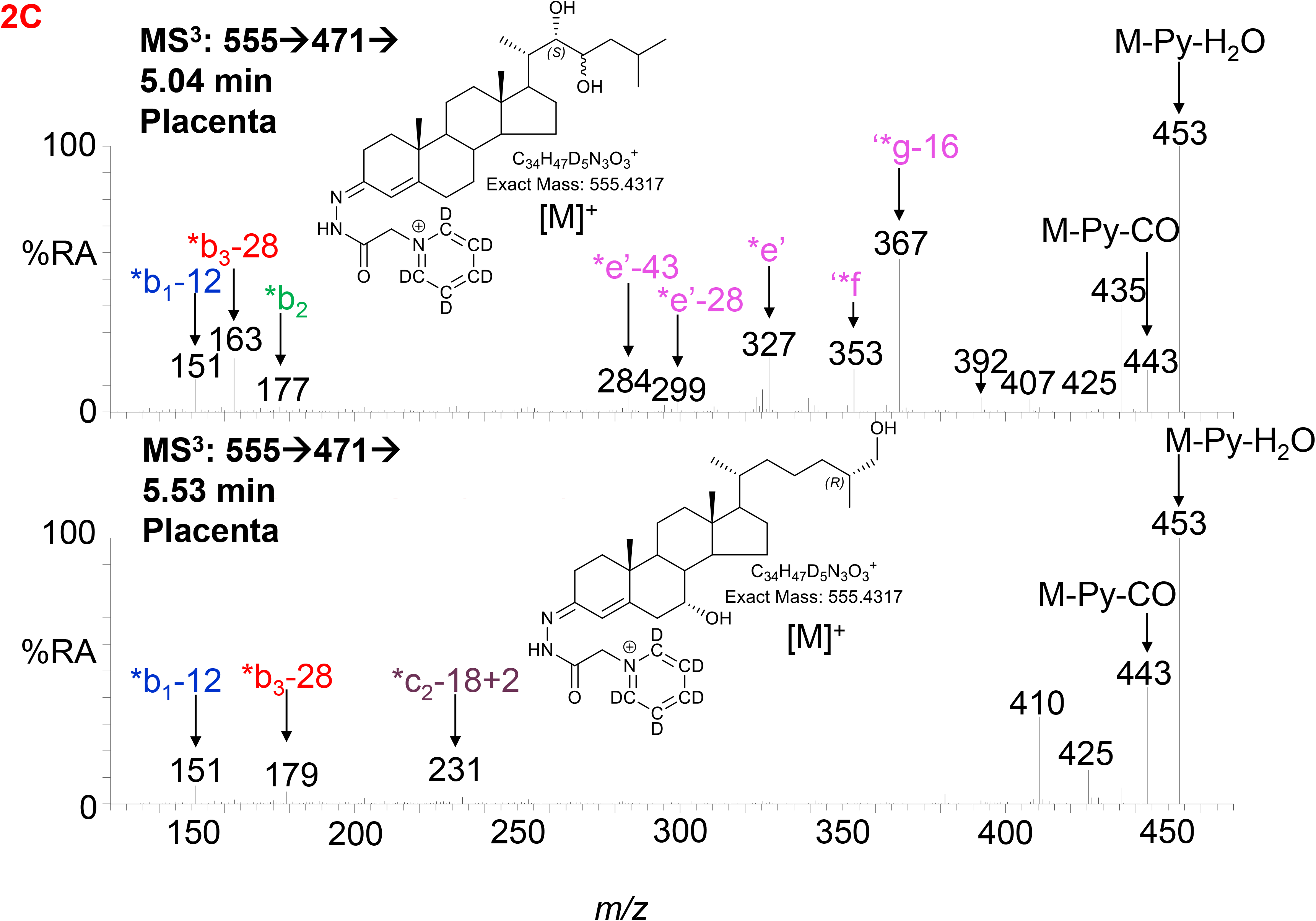
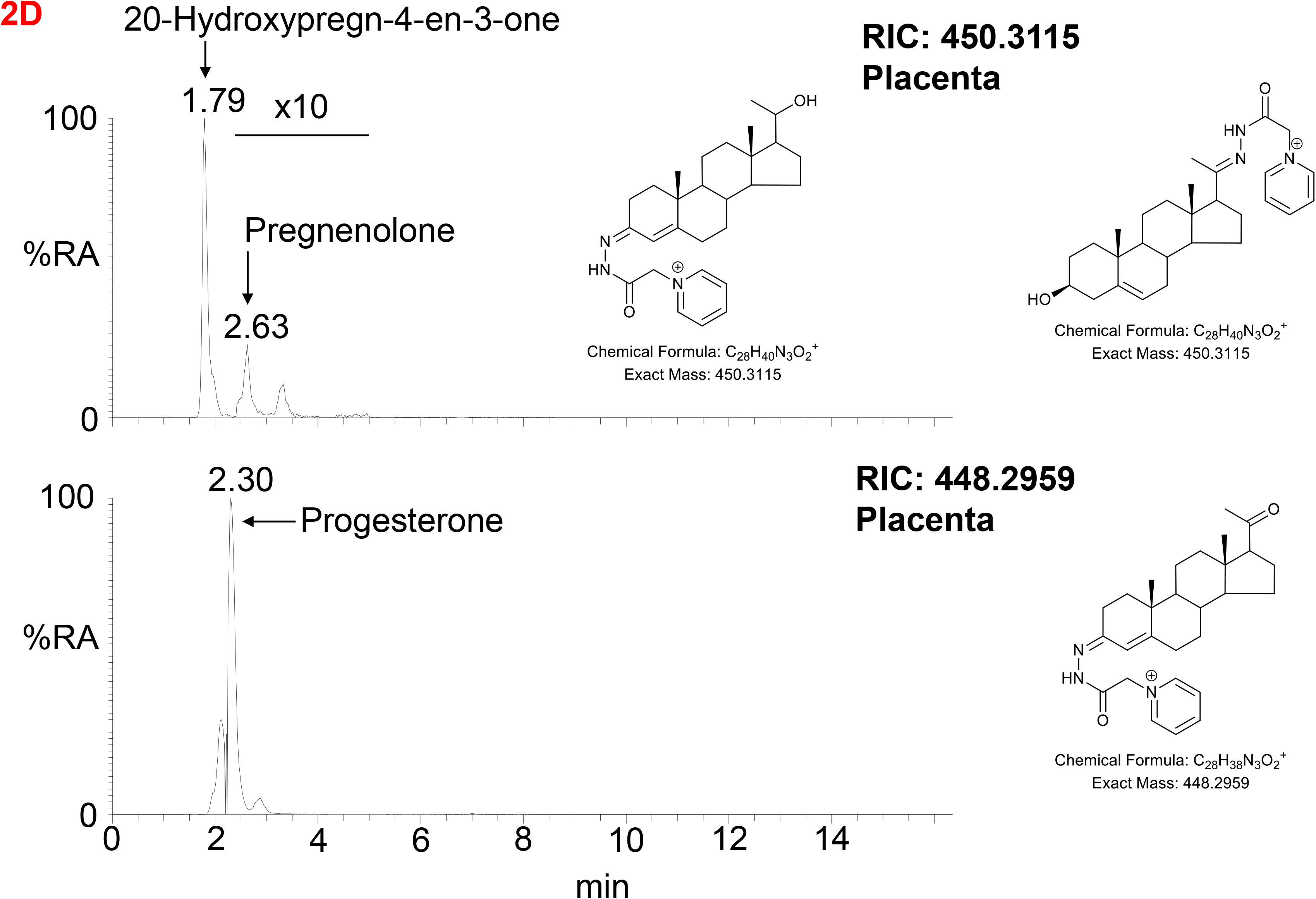
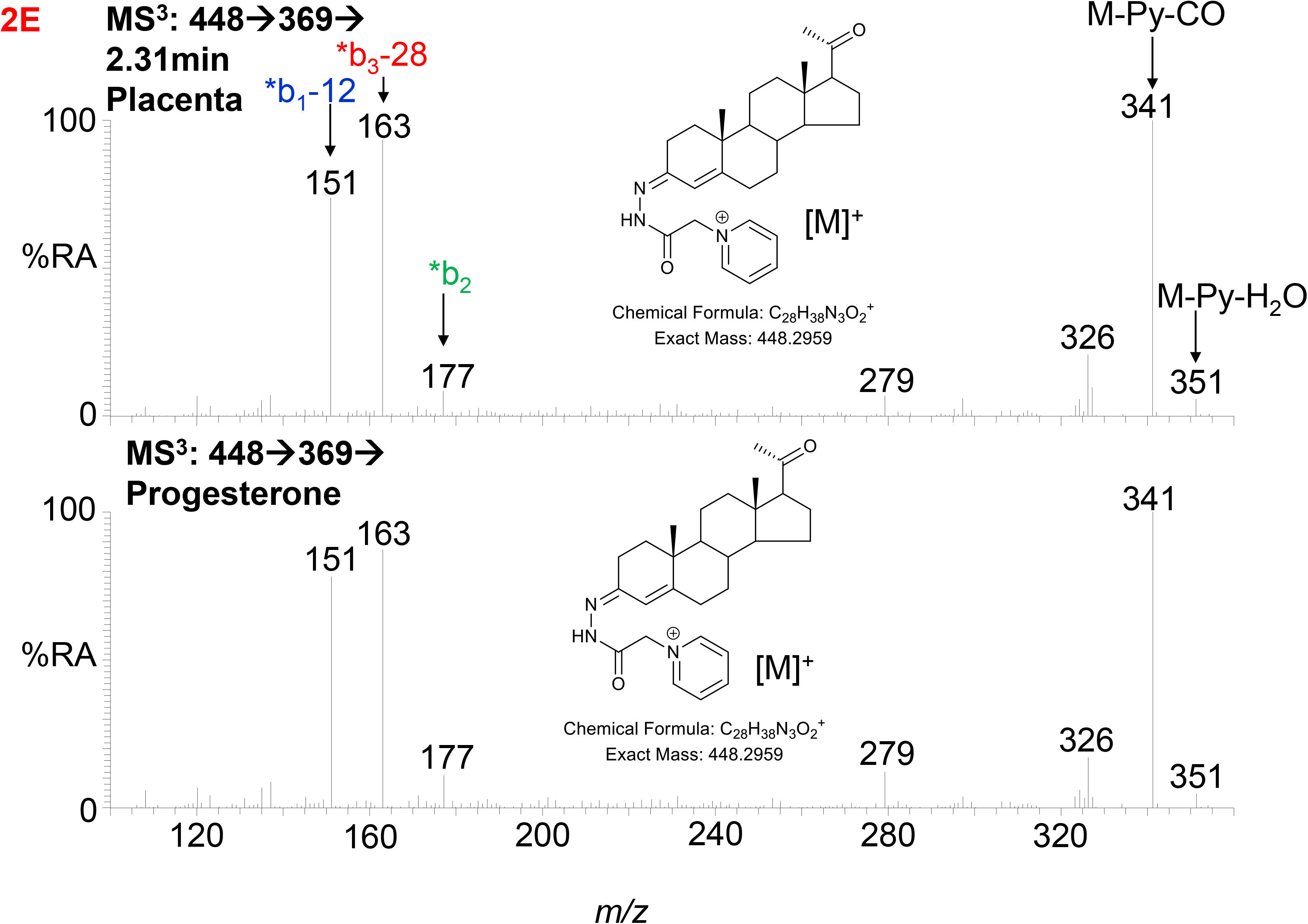
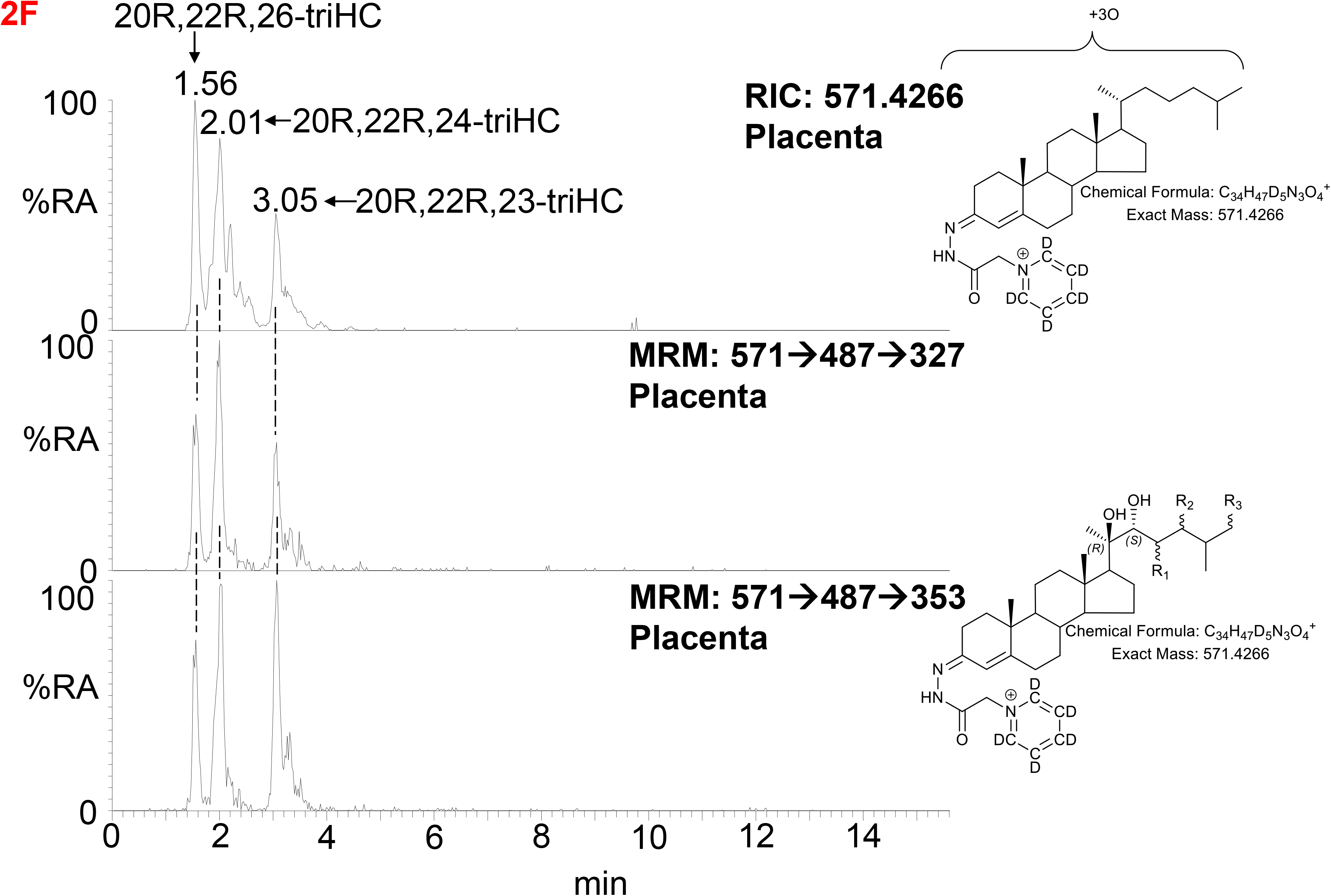
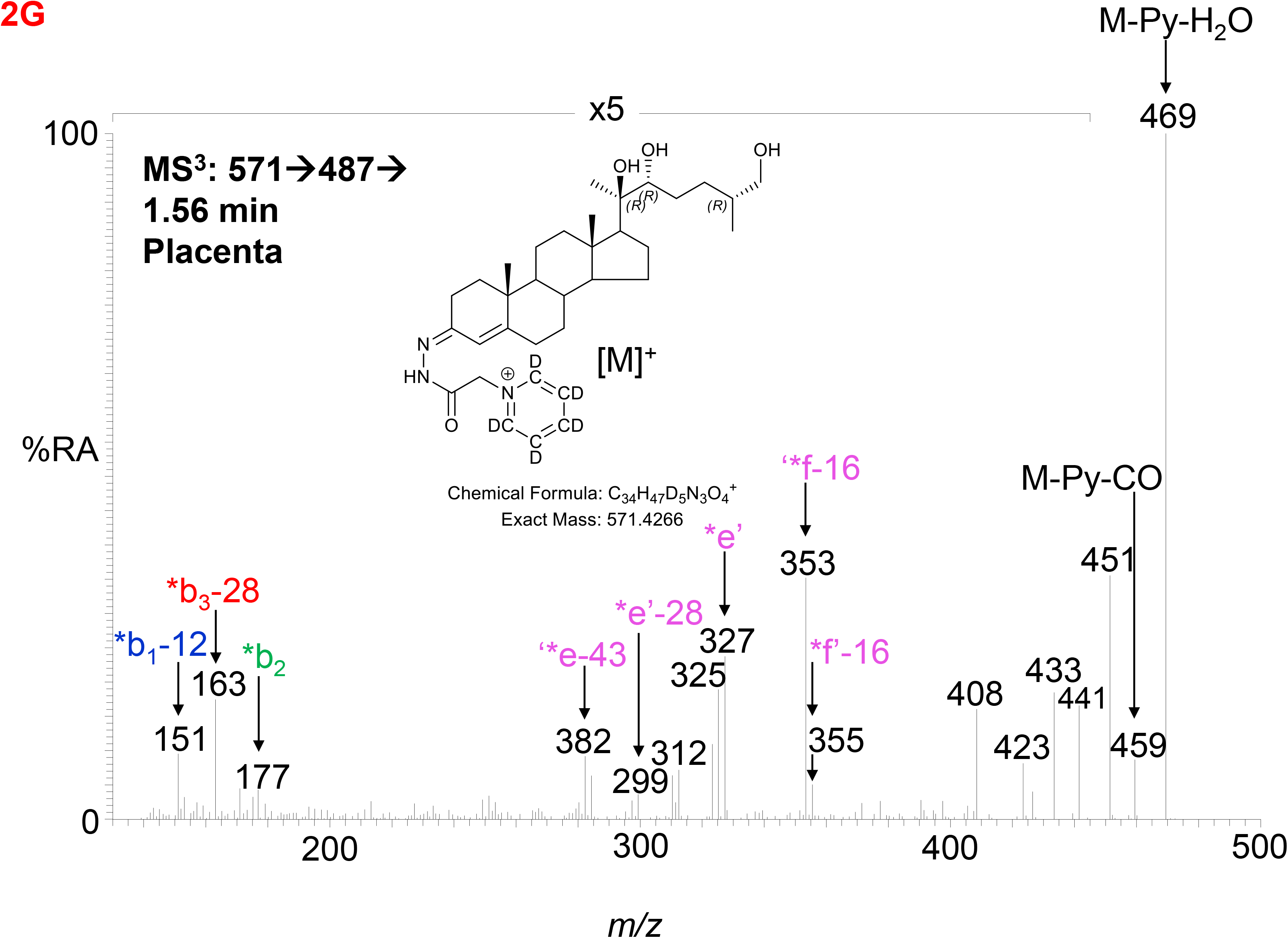
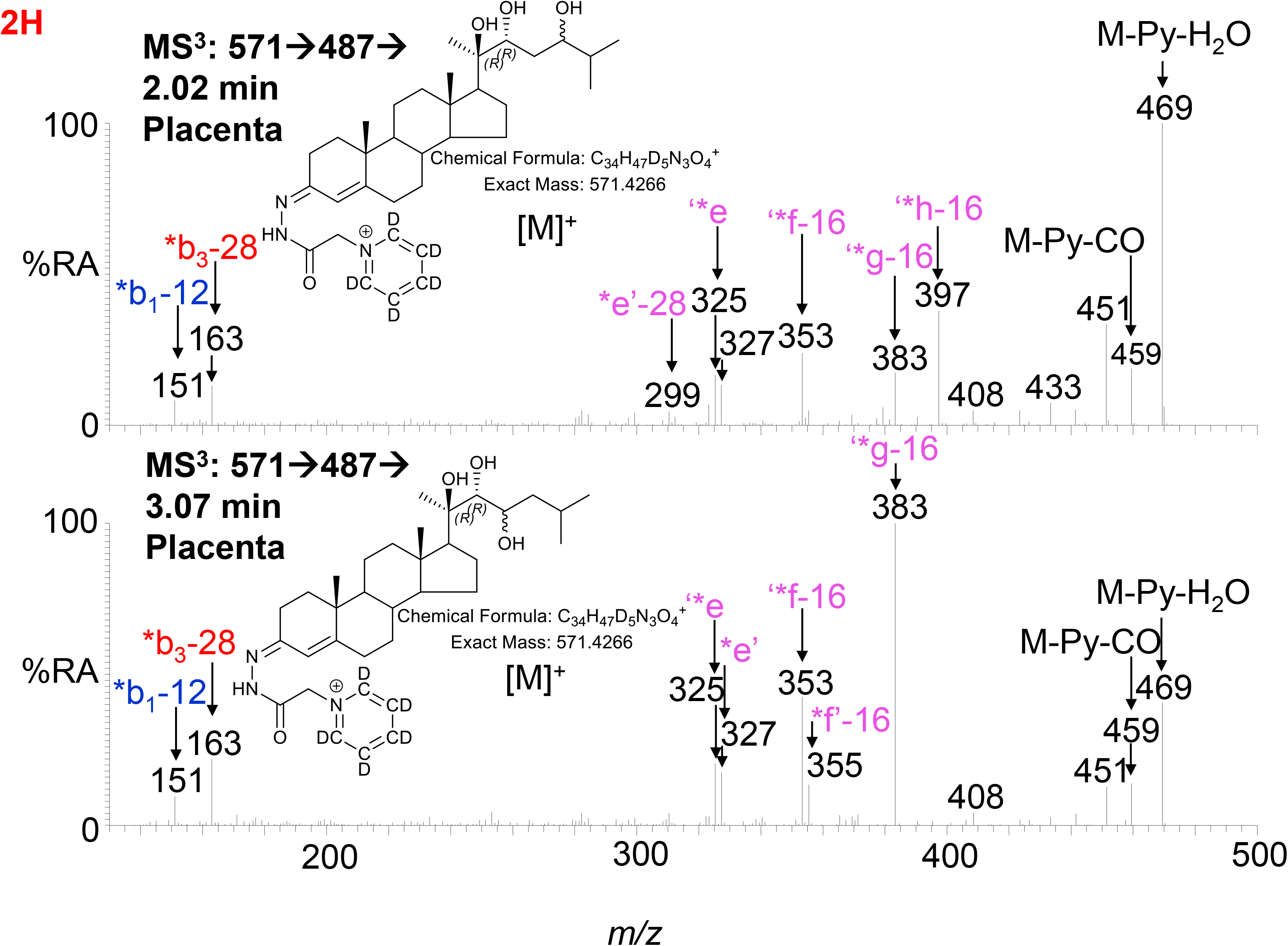
LC-MS(MS^n^) analysis of GP-derivatised 20R,22R-diHC and its metabolites in placenta. (A) RIC (*m/z* 555.4317 ± 5 ppm) corresponding to the [M]^+^ ion of dihydroxycholesterols (diHC, upper panel). MRM-like chromatograms targeting 20R,22R-diHC ([M]^+^→[M-Py]^+^→327, central panel) and ([M]^+^→[M-Py]^+^→355, lower panel). (B) MS^3^ ([M]^+^→[M-Py]^+^→) spectra of 20R,22R-diHC from placenta (upper panel) and of an authentic standard (lower panel). (C) MS^3^ ([M]^+^→[M-Py]^+^→) spectra of other dihydroxycholesterols found in placenta, possibly 22,23-diHC (upper panel) and 7α,26-diHC (lower panel). (D) RICs (*m/z* 450.3115) corresponding to pregnenolone (upper panel) and *m/z* 448.2959 corresponding to progesterone. (E) MS^3^ ([M]^+^→[M-Py]^+^→) spectra of progesterone from placenta (upper panel) and of the reference standard (lower panel). (F) RIC (*m/z* 571.4266 ± 5 ppm) corresponding to the [M]^+^ ion of trihydroxycholesterols (triHC, upper panel). MRM-like chromatograms targeting 20R,22R,x-triHC ([M]^+^→[M-Py]^+^→327, central panel) and ([M]^+^→[M- Py]^+^→353, lower panel). In the structure shown in the lower panel, one of R_1_, R_2_ or R_3_ is a hydroxy group, the other two are hydrogens. (G) MS^3^ ([M]^+^→[M-Py]^+^→) spectrum of the trihydroxycholesterol found in placenta and commensurate with the 20R,22R,26-triHC structure. (H) MS^3^ ([M]^+^→[M-Py]^+^→) spectra of other trihydroxycholesterols with possible structures of 20R,22R,24-triHC (upper panel) and 20R,22R,23-triHC.

20R,22R-diHC extracted from placenta gives intense signals in LC-MS and based on this and on the additional presumptive identification of 22,23-diHC, it is likely that other hydroxylase activities are present in placenta besides those normally associated with CYP11A1, potentially resulting in the formation of trihydroxycholesterols (triHC). The RIC for triHC (*m/z* 571.4266 ± 5 ppm) reveals three major peaks (Figure 2F upper panel). To tighten the search for hydroxylated metabolites of 20R,22R- diHC, MRM-like chromatograms were generated for the major side-chain cleavage fragment ions associated with the 20R,22R-diHC structure i.e. [M]^+^→[M-Py]^+^→327.2, [M]^+^→[M-Py]^+^→353.3 (cf. Figure 2B). Again, three major peaks were evident in these chromatograms. Interrogation of the respective MS^3^ spectra suggested isomers of 20,22,x-triHC, where x is the location of an additional hydroxy group on the side-chain. Considering first the spectrum of the latest eluting isomer at 3.05 min (Figure 2F & 2H lower panel), besides pairs of fragment ions at *m/z* 325/327 (‘*e/*e’), *m/z* 353/355 (‘*f-16/*f’-16) a dominating fragment ion is observed at *m/z* 383 (‘*g-16), this pattern is consistent with a 20R,22R,23-triHC isomer (Supplemental Figure S2F cf. S2C). Considering the isomer eluting second at 2.01 min (Figure 2F), the MS^3^ spectrum shows an additional fragment ion at *m/z* 397 (‘*h- 16, Figure 2H upper panel), the structure that explains this fragment ion formation most easily is 20R,22R,24-triHC (Supplemental Figure S2G, cf. S2C & S2D). The MS^3^ spectrum of the first eluting peak at 1.56 min (Figure 2G) does not show the fragment-ions at *m/z* 383 or 397 (Figure cf. 2H), suggesting that the extra hydroxy group does not encourage additional side-chain fragmentation. The most likely structures for this isomer are 20R,22R,26-triHC or perhaps 20R,22R,25-triHC (Supplemental Figure S2H & S2I, cf. S2C & S2D).

#### 3.1.3. Cholestenoic Acids

Sterol 27-hydroxylase (CYP27A1) is the enzyme that introduces both (25R)26-hydroxy and (25R)26- carboxy groups to sterols (23), it is expressed in trophoblast cells of the placenta (24). We identify 26- HC (Figure 1A & 1H, Supplemental Figure S3D) and 3β-hydroxycholest-5-en-(25R)26-oic acid (3β-HCA, Supplemental Figure S4) in placenta, and there is the possibility that 20R,22R-diHC may be a substrate for CYP27A1 and be metabolised via 20R,22R,26-triHC to the C_27_ bile acid 3β,20R,22R- trihydroxycholest-5-en-(25R)26-oic acid (3β,20R,22R-triHCA) in placenta. The RIC appropriate for 3β,20R,22R-triHCA (*m/z* 585.4059 ± 5ppm) reveals two major and a minor peak (Figure 3A) of which only the first gives an MS^3^ spectrum compatible with a 3β,20R,22R-triHCA structure (Figure 3B & Supplemental Figure S2J & S2K, cf. Supplemental Figure S2H & S2I). The similarity between the MS^3^ spectra of 20R,22R-diHC, and the presumptively identified 20R,22R,26-triHC and 3β,20R,22R-triHCA can be visualised in Figure 3C where the low-middle *m/z* range of the three spectra are shown on the same *m/z* scale. While the low-middle *m/z* range provides evidence for the 20R,22R-dihydroxy structural motif, the high *m/z* range (Figure 3B) is indicative of a C-26 acid. Sterol acids show characteristic neutral losses from the [M-Py]^+^ ion corresponding to the net loss of H_2_CO_2_ + n(H_2_O), where n is the number of OH groups on the sterol beyond that derivatised at C-3 (19). In Figure 3B we see such losses giving fragments at *m/z* 437 (M-Py-H_2_O-H_2_CO_2_ i.e. ‘*j-18) and *m/z* 419 (M-Py-2H_2_O-H_2_CO_2_ i.e. ‘*j-36). These fragment ions are associated with satellite peaks at *m/z* 440 ([M-Py-61]^+^ and 422 ([M-Py-79]^+^) providing a pattern characteristic of a trihydroxycholestenoic acid (See Supplemental Figure S2L). Note, the third hydroxy group is the site of derivatisation. The other two chromatographic peaks give almost identical MS^3^ spectra (Figure 3D) presumably *syn* and *anti* conformers of a second isomer whose structure is not obvious from the MS^3^ spectra, although fragment-ions at *m/z* 385, 397 and 471 are probably structurally significant.

**Figure 3.**
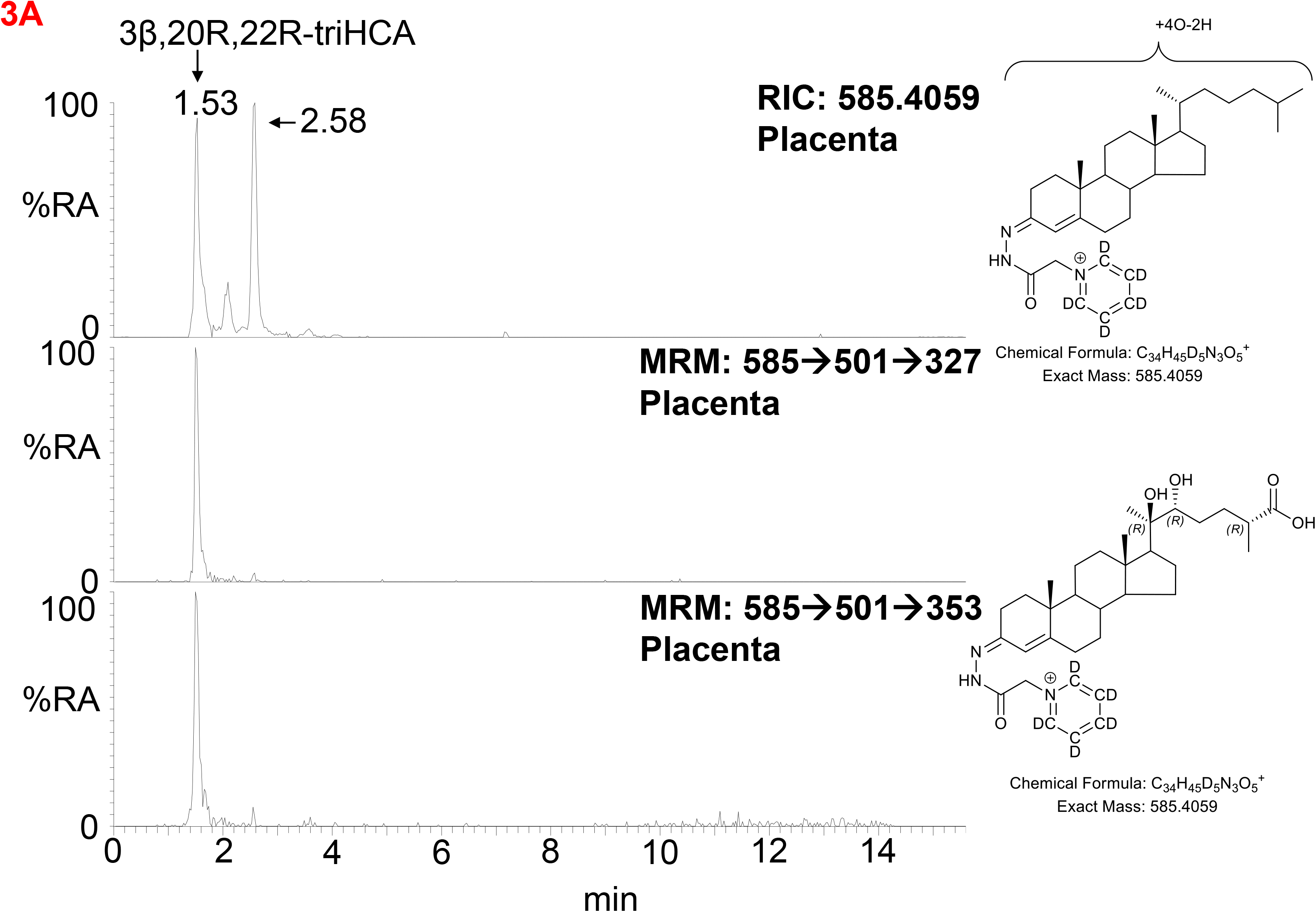
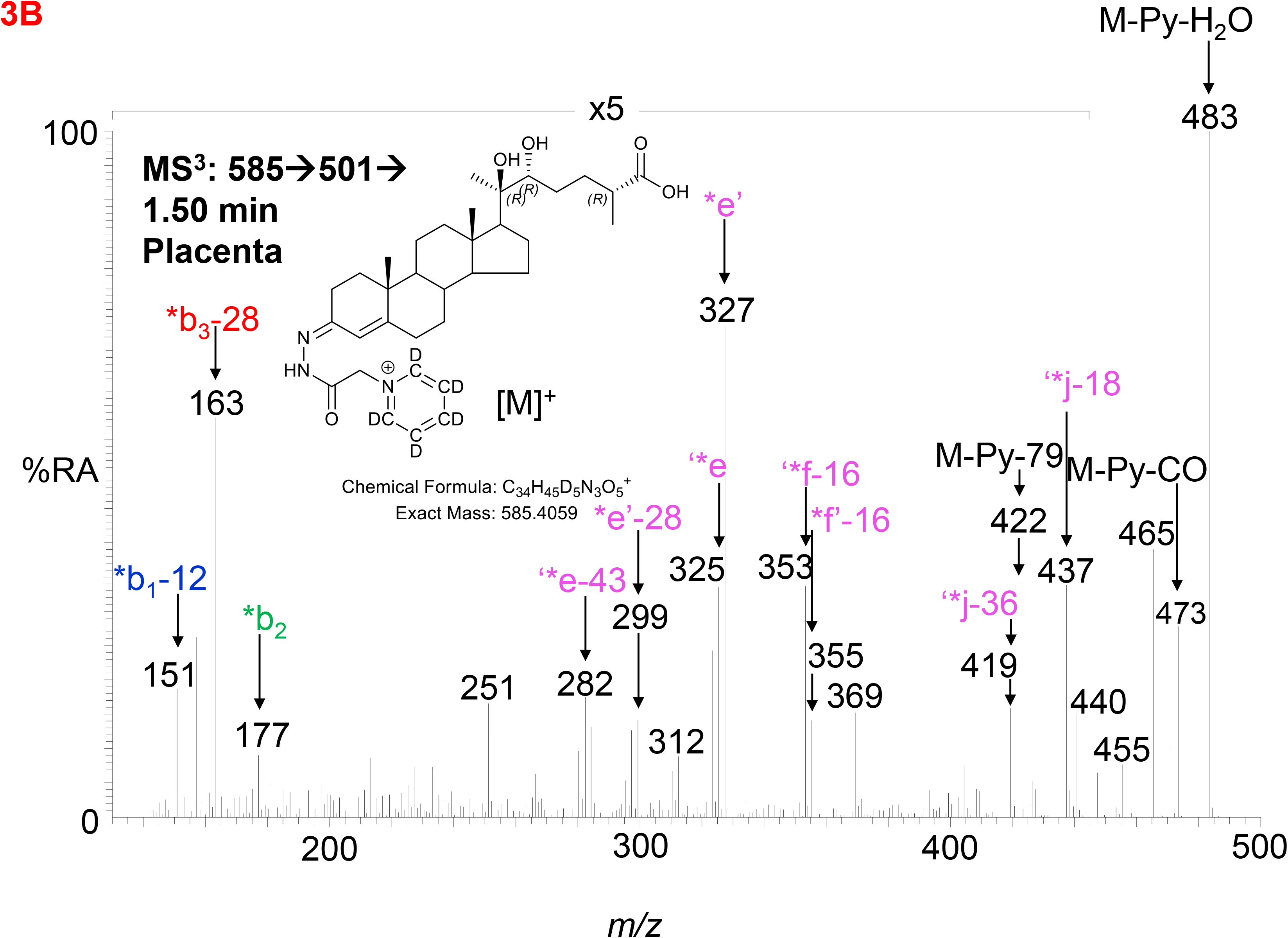
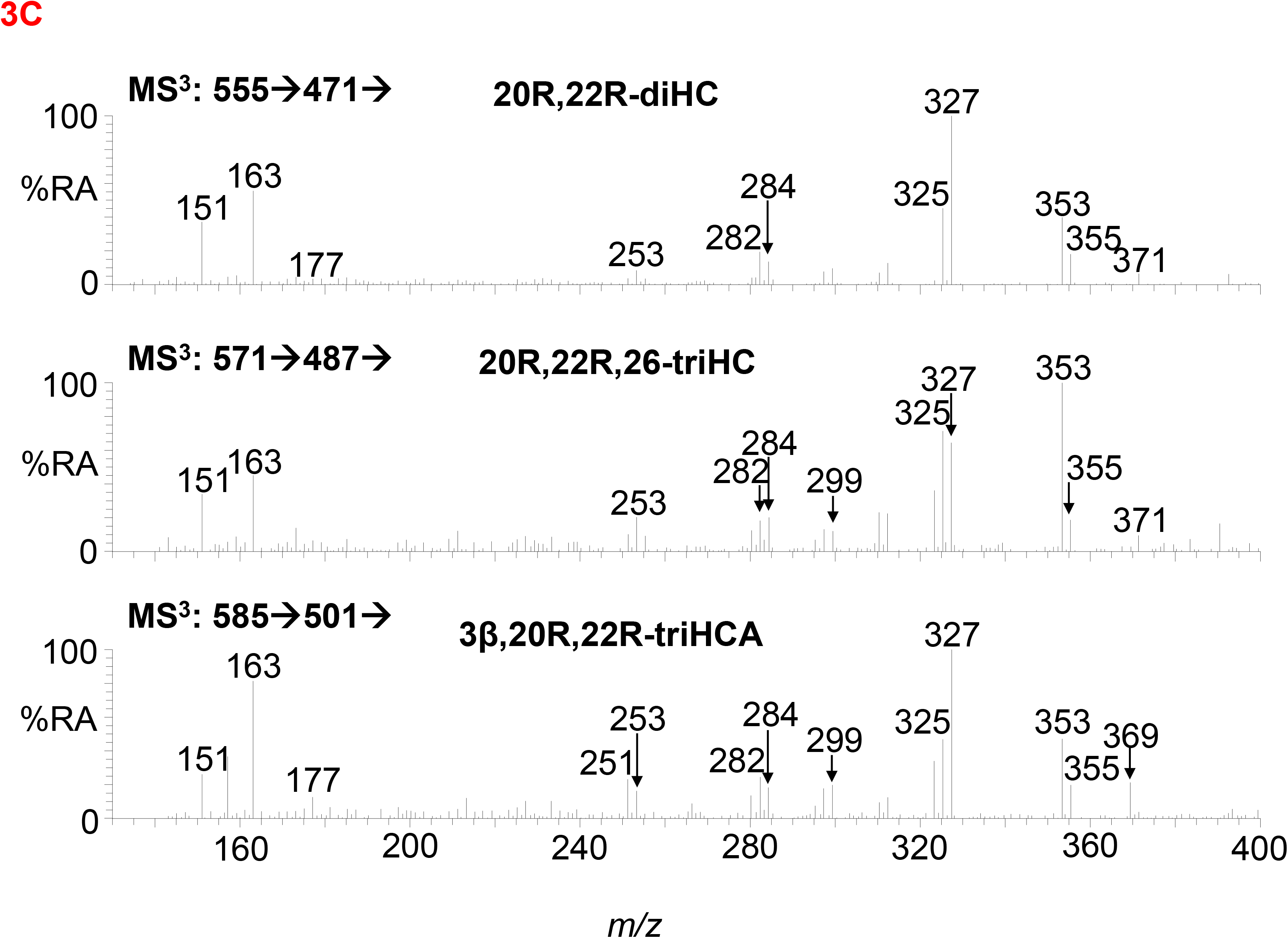
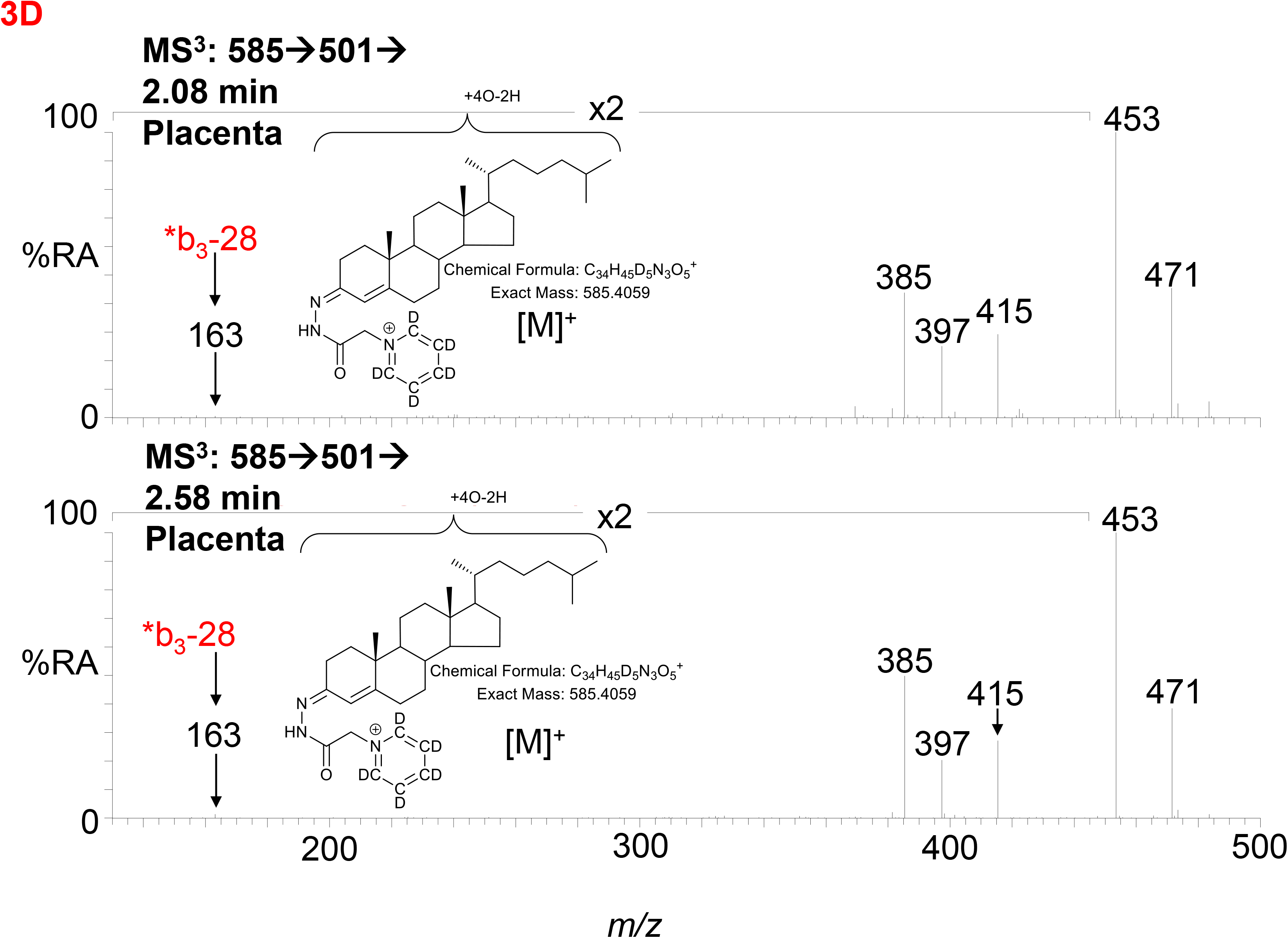
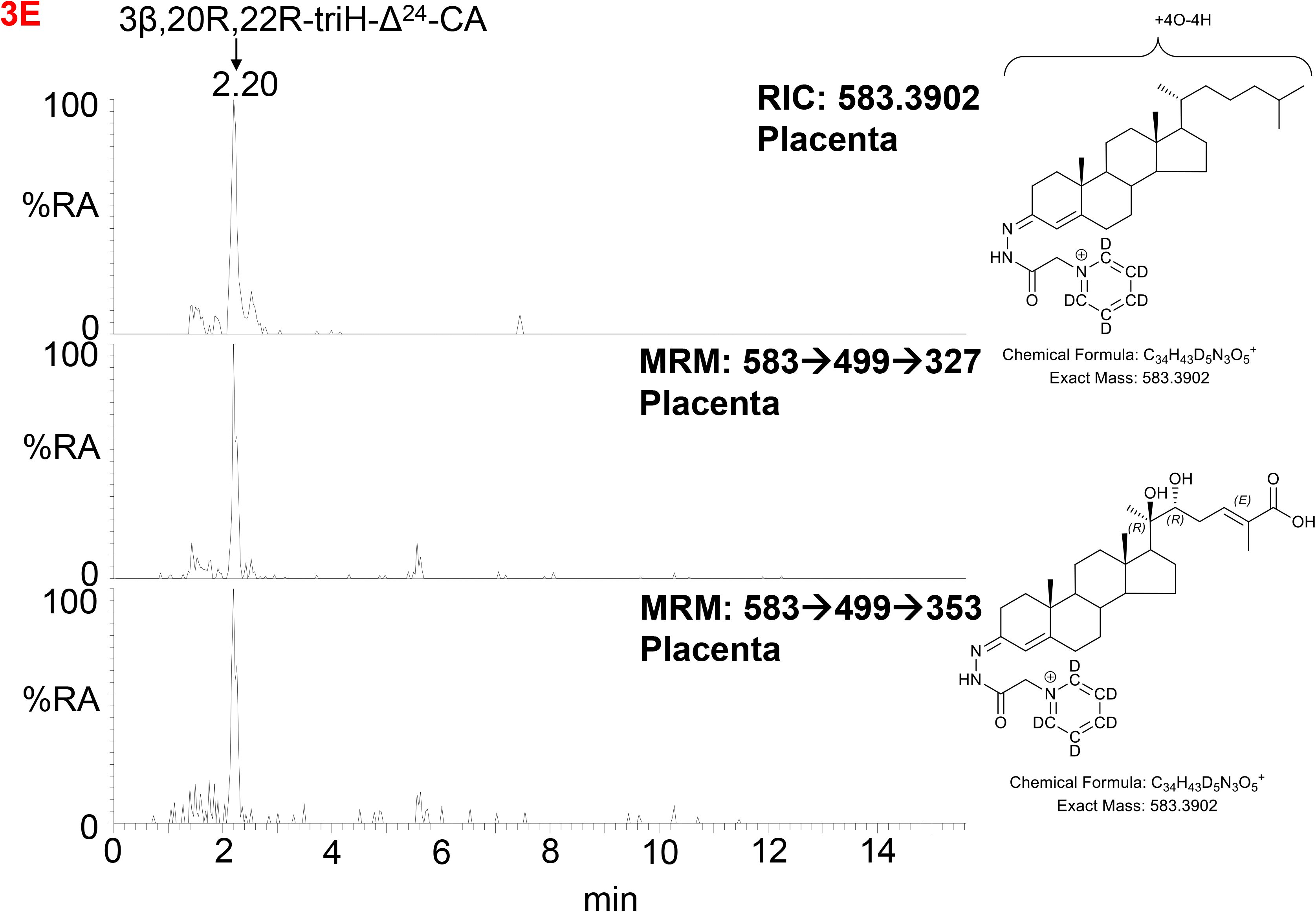
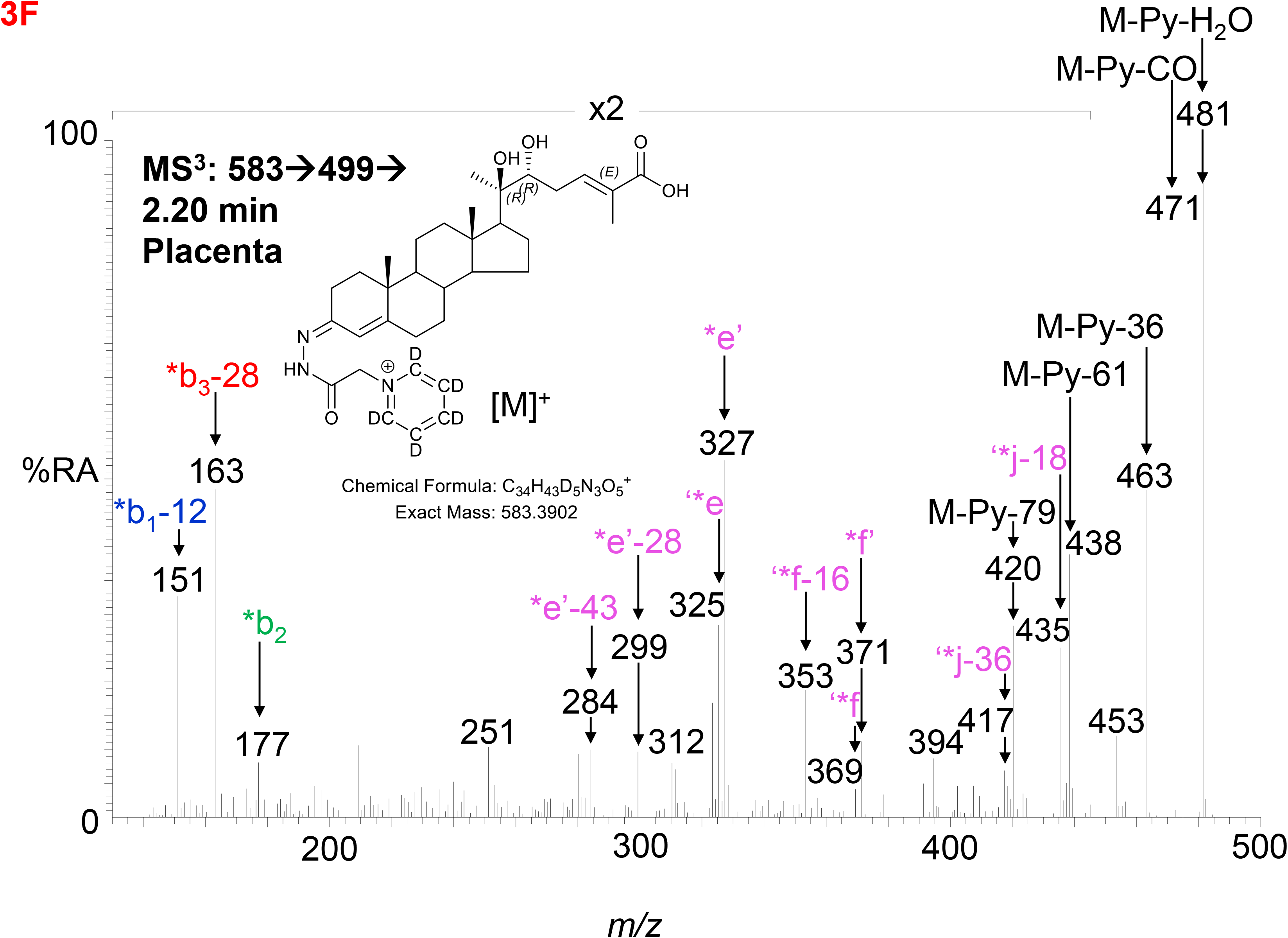
LC-MS(MS^n^) analysis of GP-derivatised 3β,20R,22R-triHCA, its isomers and 3β,20R,22R-triH- Δ^24^-CA in placenta. (A) RIC (*m/z* 585.4059 ± 5 ppm) corresponding to the [M]^+^ ion of trihydroxycholestenoic acids (upper panel). MRM-like chromatograms targeting 3β,20R,22R-triHCA ([M]^+^→[M-Py]^+^→327, central panel) and ([M]^+^→[M-Py]^+^→353, lower panel). (B) MS^3^ ([M]^+^→[M- Py]^+^→) spectrum postulated to correspond to 3β,20R,22R-triHCA. (C) Comparison of MS^3^ ([M]^+^→[M- Py]^+^→) spectra of 20R,22R-diHC (upper panel) with spectra postulated to correspond to 20R,22R,26- triHC (central panel) and 3β,20R,22R-triHCA (lower panel) over the *m/z* range 130 – 400. (D) MS^3^ spectra of two other isomers of 3β,20R,22R-triHCA, the spectra do not appear to be of cholestenoic acids. (E) RIC (*m/z* 583.3902 ± 5 ppm) corresponding to the [M]^+^ ion of a trihydroxycholestadienoic acids (upper panel). MRM-like chromatograms targeting 3β,20R,22R-triH-Δ^24^-CA ([M]^+^→[M- Py]^+^→327, central panel) and ([M]^+^→[M-Py]^+^→353, lower panel). (F) MS^3^ ([M]^+^→[M-Py]^+^→) spectrum postulated to correspond to 3β,20R,22R-triH-Δ^24^-CA.

Cholestenoic acids are intermediates in bile acid biosynthesis pathways (25–27). In the route towards C_24_ acids, C_27_ acids become converted to their CoA-thioesters by bile acid CoA-synthetase (BACS, SLC27A5), stereochemistry at C-25 is inverted by α-methylacyl-CoA racemase (AMACR) and a double bond introduced at Δ^24^ with *trans* geometry by acyl-CoA oxidase 2 (ACOX2) (26). The next steps are catalysed by D-bifunctional protein (DBP, HSD17B4) lead to a C-24 oxo group. Beta-oxidation by sterol carrier protein 2 (SCP2) then gives a C_24_ CoA-thioester which is finally amidated with glycine or taurine or hydrated to give the C_24_ acid (26). In bioanalysis, intermediates are almost always observed as the carboxylic acids rather than the thioesters (27). Following this pathway, the CoA-thioester of 3β,20R,22R-triHCA would be isomerised from the 25R-epimer to one with 25S-stereochemistry which would then be oxidised to the CoA-thioester of 3β,20R,22R-trihydoxycholest-5,24-dienoic acid (3β,20R,22R-triH-Δ^24^-CA). Shown in Figure 3E is the RIC (*m/z* 583.3902 ± 5 ppm) appropriate to the [M]^+^ ion of 3β,20R,22R-triH-Δ^24^-CA, along with MRM-like chromatograms characteristic of sterols with 20- and 22-hydroxylation of the side-chain. One major chromatographic peak is evident at 2.20 min, and the MS^3^ ([M]^+^→[M-Py]^+^→) spectrum associated with this peak (Figure 3F) resembles, in the low to middle *m/z* range, that of 3β,20R,22R-triHCA (Figure 3B), however, the fragment-ion observed at *m/z* 355 (*f’-16) in spectrum of 3β,20R,22R-triHCA is replaced by one at *m/z* 371 (*f’) in the spectrum shown Figure 3F. This spectrum is compatible with the 3β,20R,22R-triH-Δ^24^-CA structure (see Supplemental Figure S2M - O). Further evidence for the MS^3^ spectrum presented in Figure 3F corresponding to 3β,20R,22R-triH-Δ^24^-CA is the presence of fragment ions characteristic of sterol acids, ‘*j-18 (M-Py-H_2_O-H_2_CO_2_) at *m/z* 435 and ‘*j-36 (M-Py-2H_2_O-H_2_CO_2_) at *m/z* 417. These fragment ions are associated with satellite peaks at *m/z* 438 ([M-Py-61]^+^ and 420 ([M-Py-79]^+^, see Supplemental Figure S2N). However, as is the case with other postulated structures definitive identification awaits synthesis of the authentic standard.

### 3.2. Identification of Oxysterols in Cord and Maternal Plasma

To investigate if the placental oxysterols are transported to the fetus, umbilical cord plasma derived from umbilical cord blood was analysed for oxysterols. The data was compared to the oxysterol profiles in plasma from maternal blood, taken 1 - 2 day before elective caesarean section and plasma from “control” non-pregnant females. As might be expected, the oxysterol profile of cord plasma resembles that of non-pregnant females, but with the additional presence of CYP11A1-derived oxysterols. 22R-HC is present in both cord and maternal plasma but is absent from controls (Figure 4A & 4B, Figure 5A - C, Table 1). If present, 20S-HC is at levels in cord, maternal and control plasma samples below the limit of detection (0.1 ng/mL). 20R,22R-diHC is the dominant oxysterol in cord plasma, it is also a major oxysterol in maternal plasma but is absent from control plasma (Figure 4C – D, Figure 5D - F). Presumptively identified 20R,22R,26-triHC was near the limit of quantification in both cord and maternal plasma but was absent from controls (Figure 4E & 4F, Figure 5G - H). The presumptively identified C_27_ bile acid 3β,20R,22R-triHCA was evident in cord plasma and just detected in maternal plasma but was absent from control plasma (Figure 4G & 4H, Figure 6A - C). Presumptively identified 3β,20R,22R-triH-Δ^24^-CA was only detected in cord plasma (Supplemental Figure S5A & S5B).

**Figure 4.**
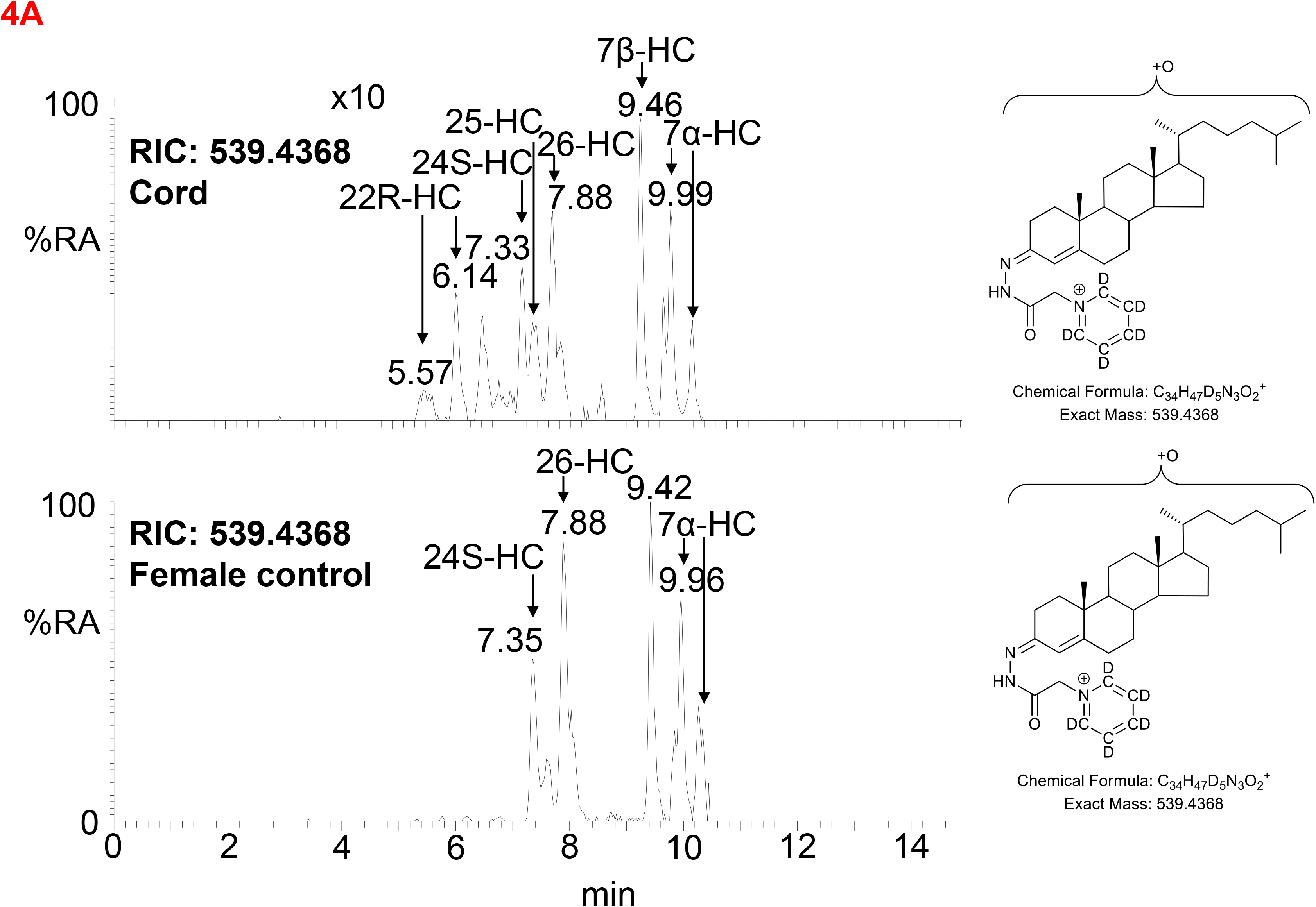
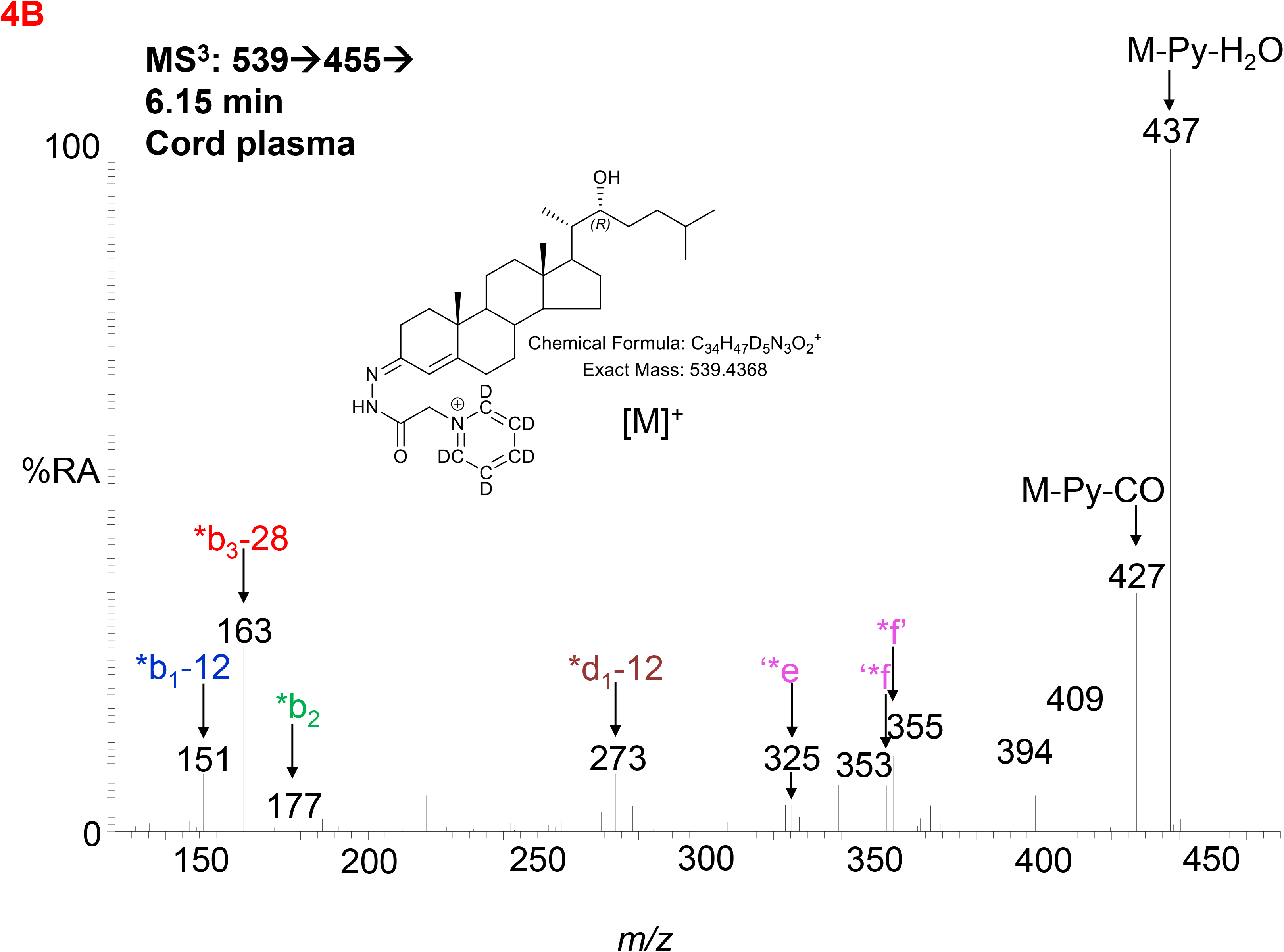
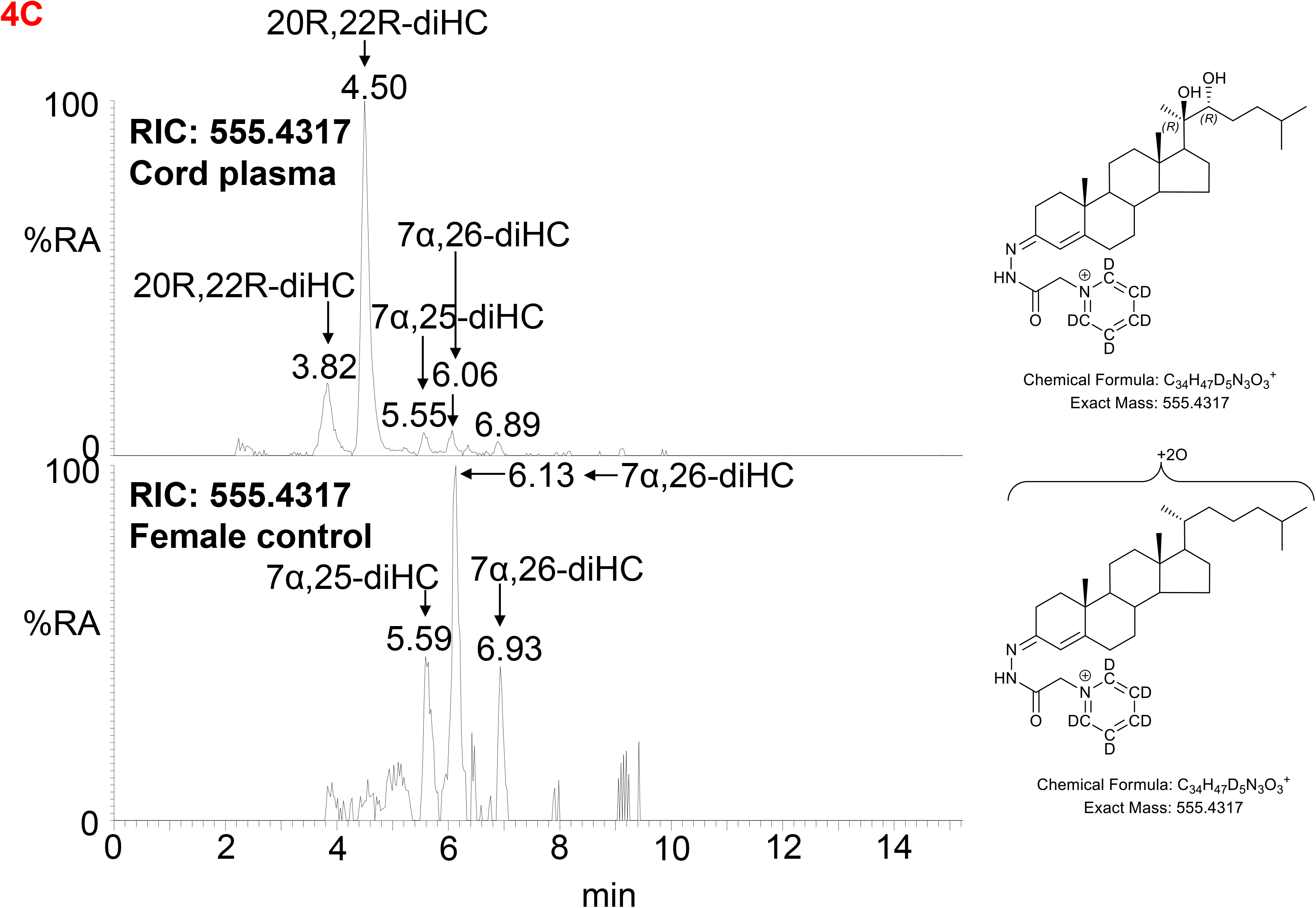
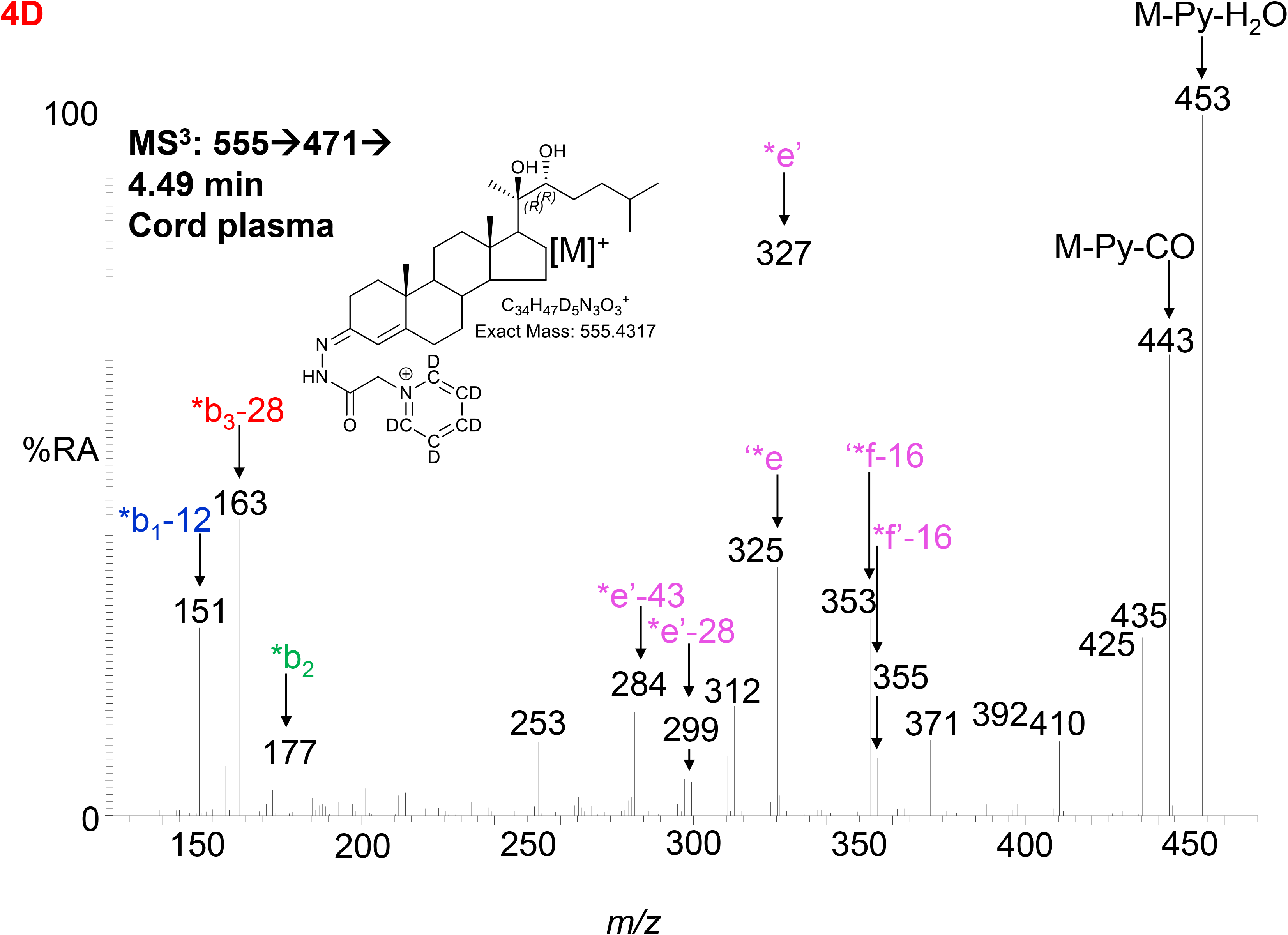
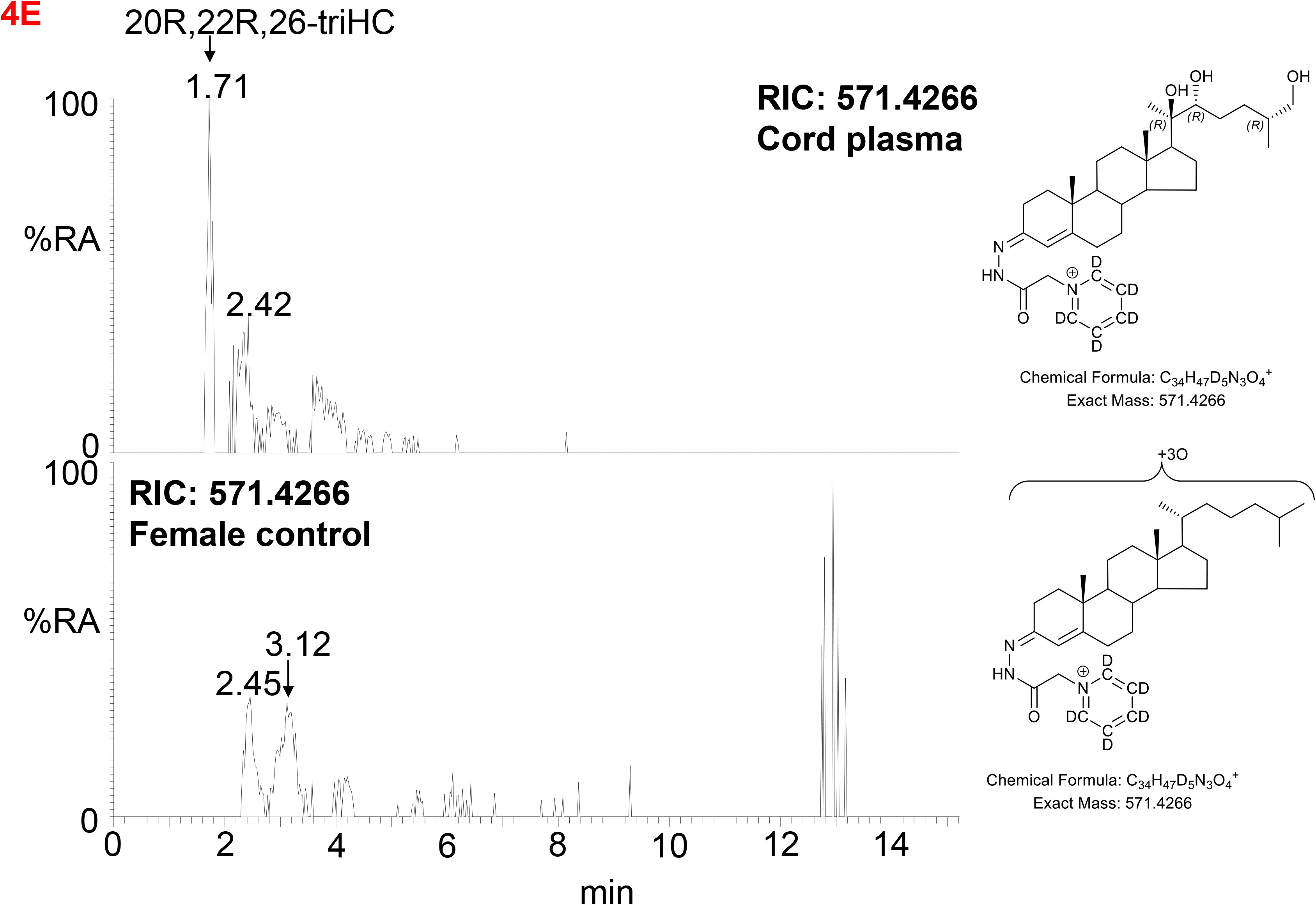
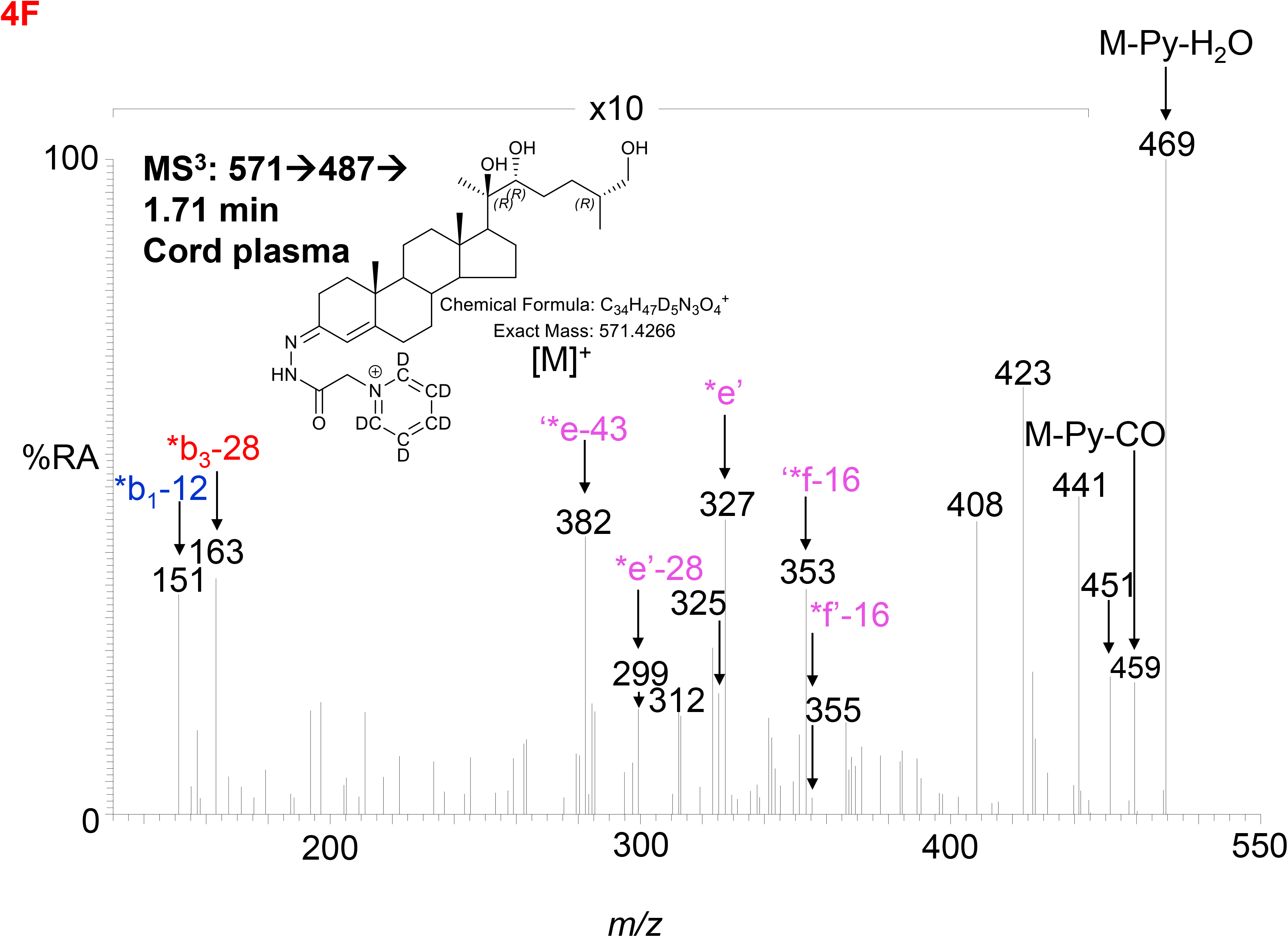
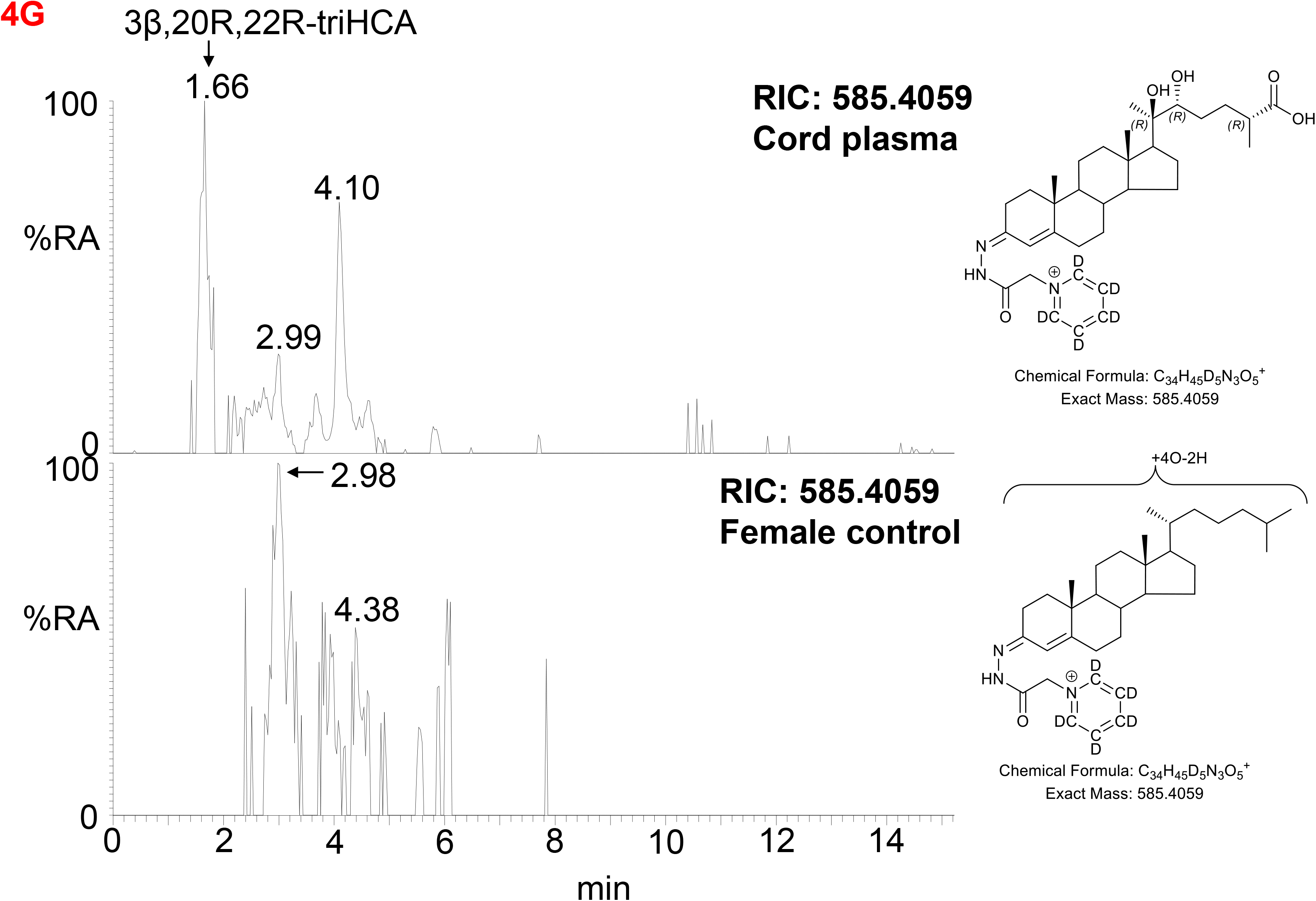
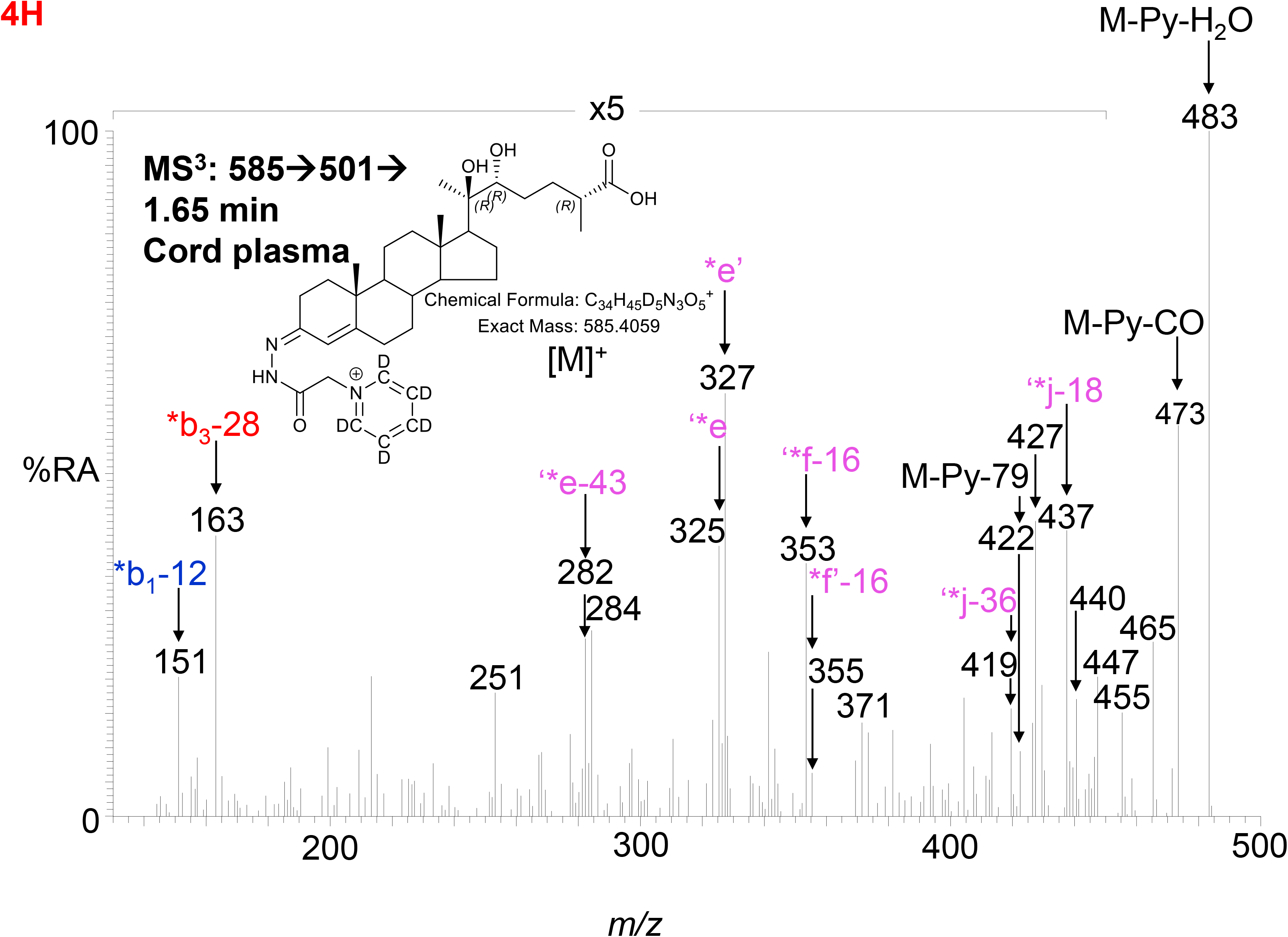
LC-MS(MS^n^) analysis of GP-derivatised oxysterols in umbilical cord plasma. (A) RICs for monohydroxycholesterols (*m/z* 539.4368 ± 5 ppm) in cord (upper panel) and control non-pregnant female plasma (lower panel). (B) MS^3^ ([M]^+^→[M-Py]^+^→) spectrum of 22R-HC from cord plasma. (C) RICs for dihydroxycholesterols (*m/z* 555.4317 ± 5 ppm) in cord (upper panel) and control female plasma (lower panel). (D) MS^3^ ([M]^+^→[M-Py]^+^→) spectrum of 20R,22R-diHC from cord plasma. (E) RICs for trihydroxycholesterols (*m/z* 571.4266 ± 5 ppm) in cord (upper panel) and control female plasma (lower panel). (F) MS^3^ ([M]^+^→[M-Py]^+^→) spectrum postulated to correspond to 20R,22R,26-triHC. (G) RIC (*m/z* 585.4059 ± 5 ppm) corresponding to the [M]^+^ ion of trihydroxycholestenoic acids in cord plasma (upper panel) and control female plasma. (H) MS^3^ ([M]^+^→[M-Py]^+^→) spectrum postulated to correspond to 3β,20R,22R-triHCA. Note, cord and control female plasma were analysed using a different LC column (same type, different batch) to placenta.

**Figure 5.**
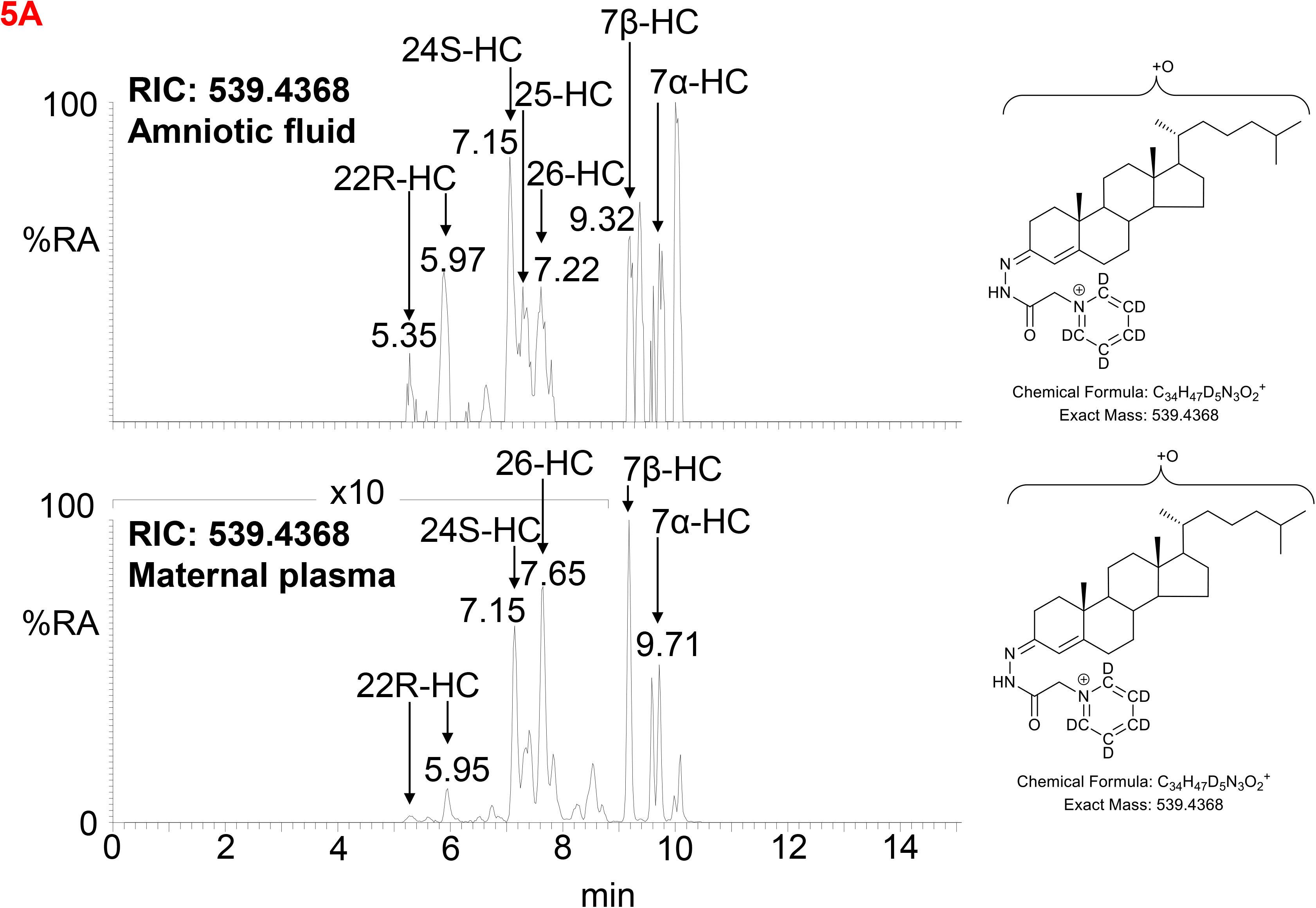
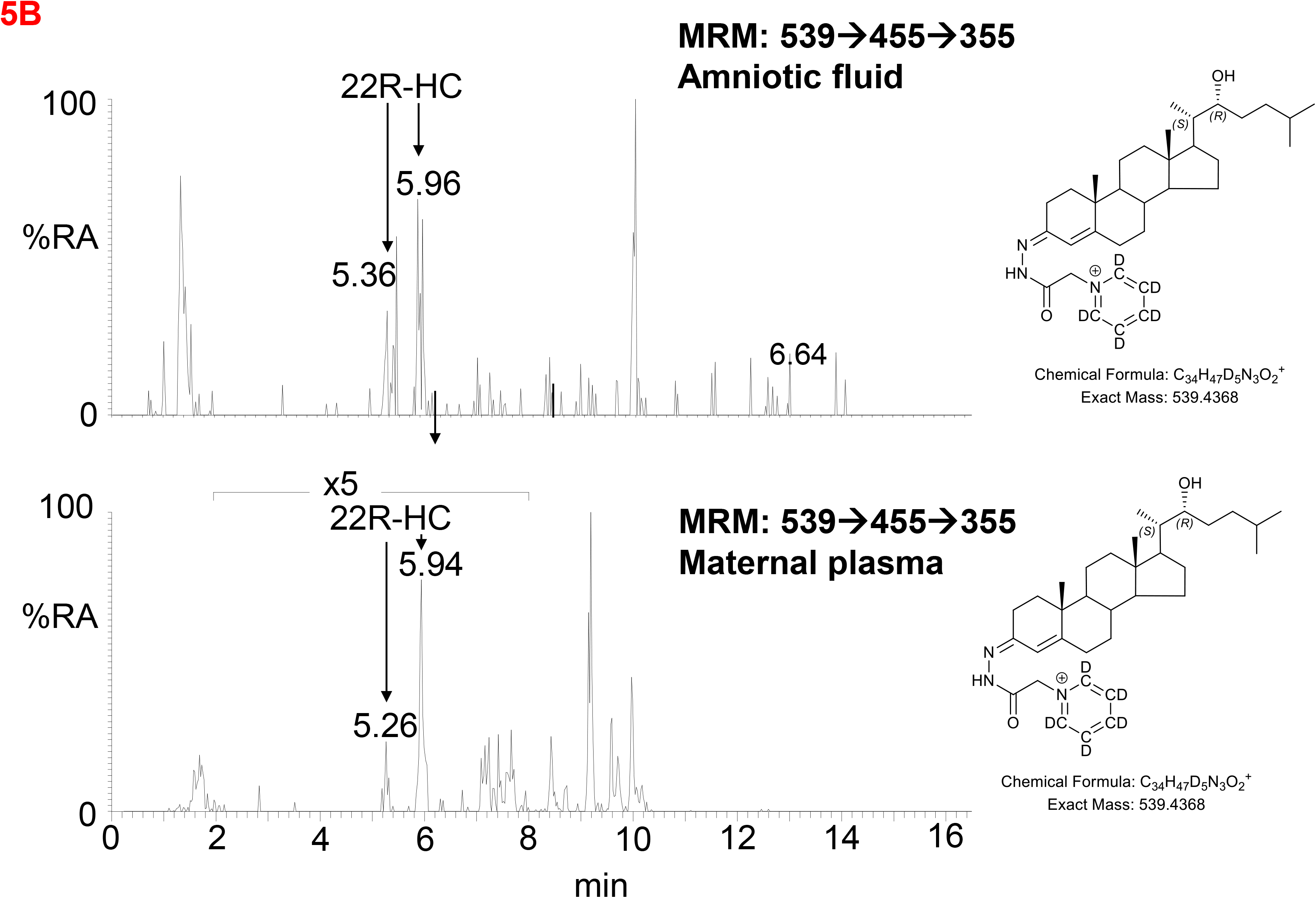
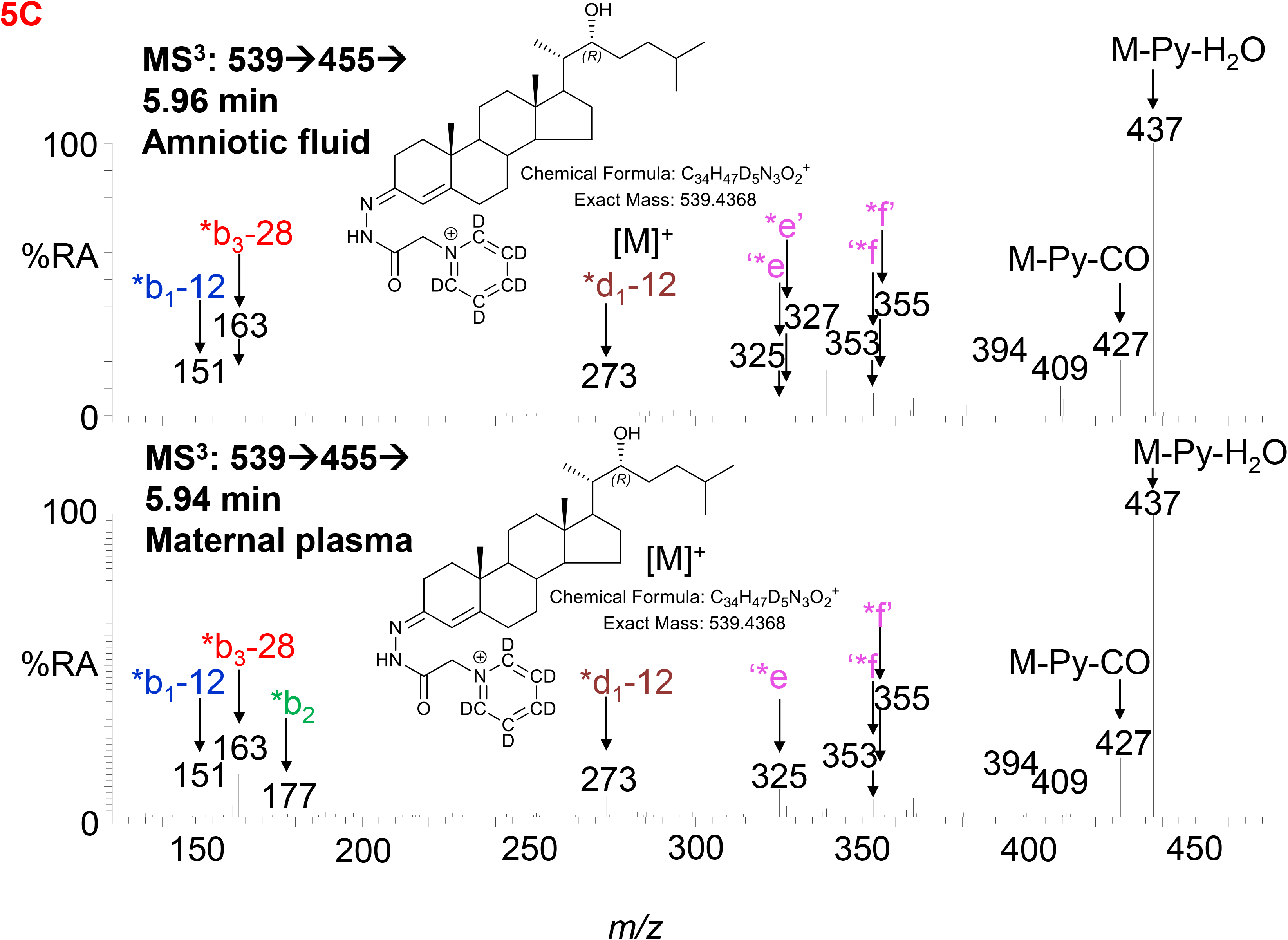
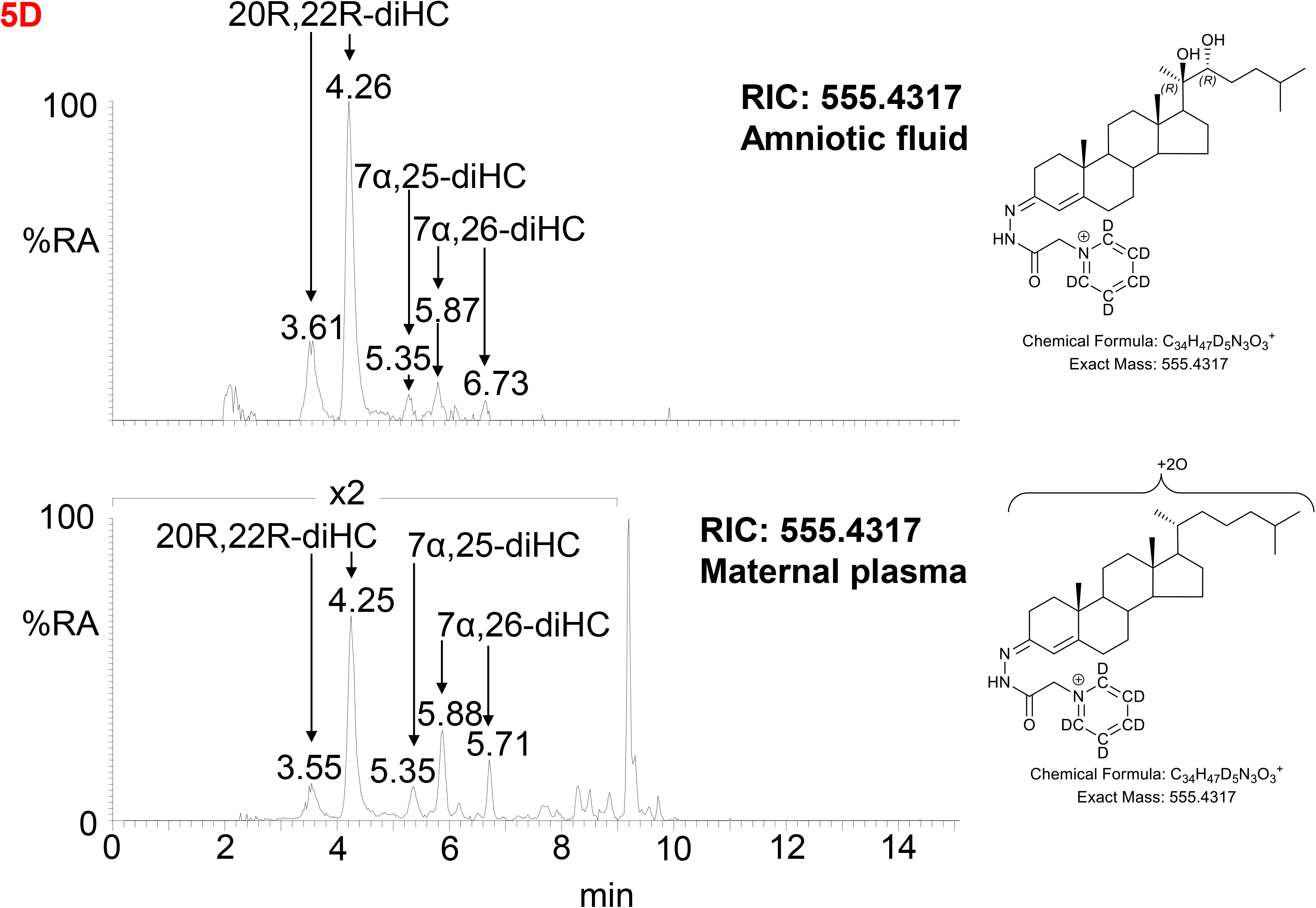
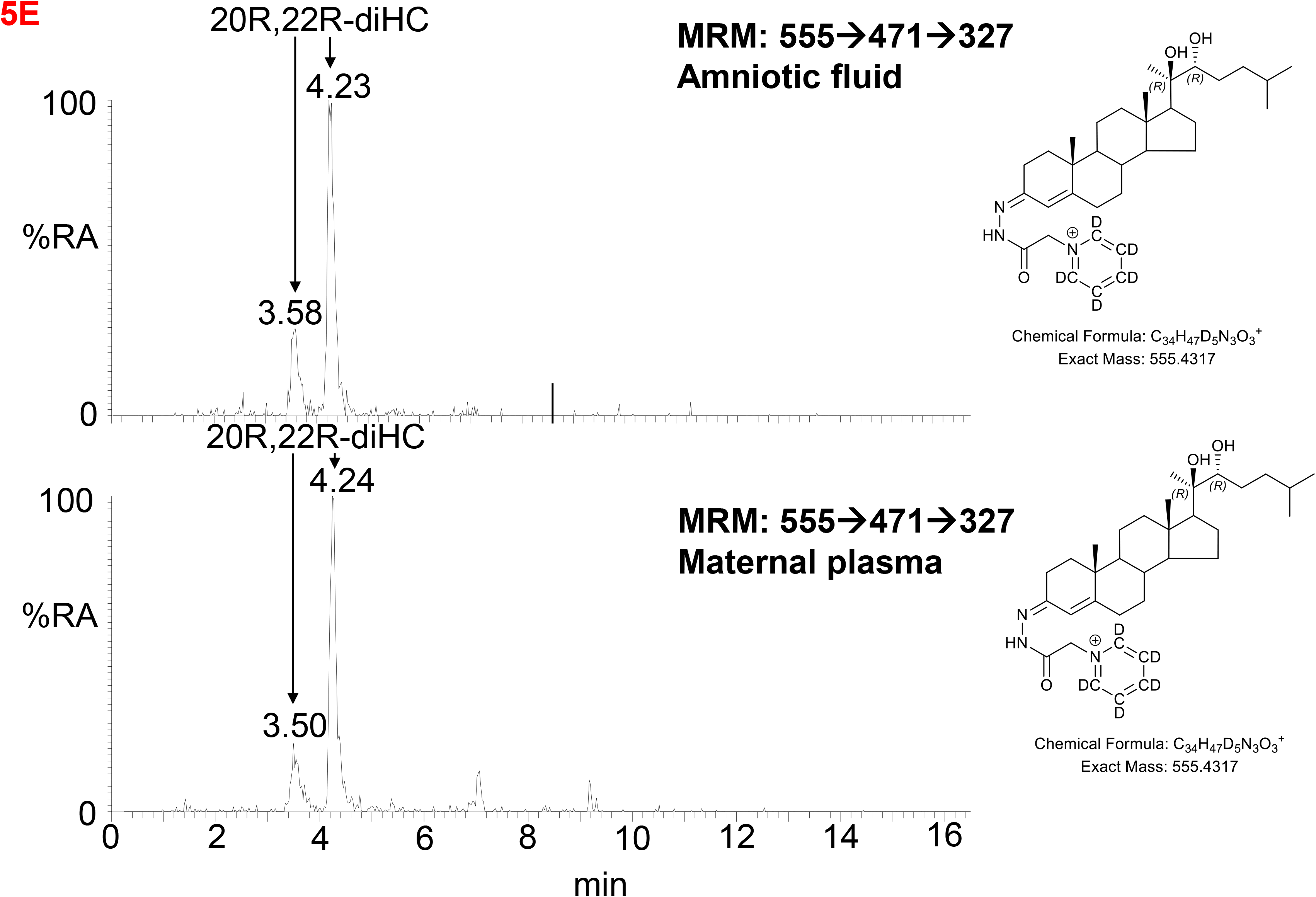
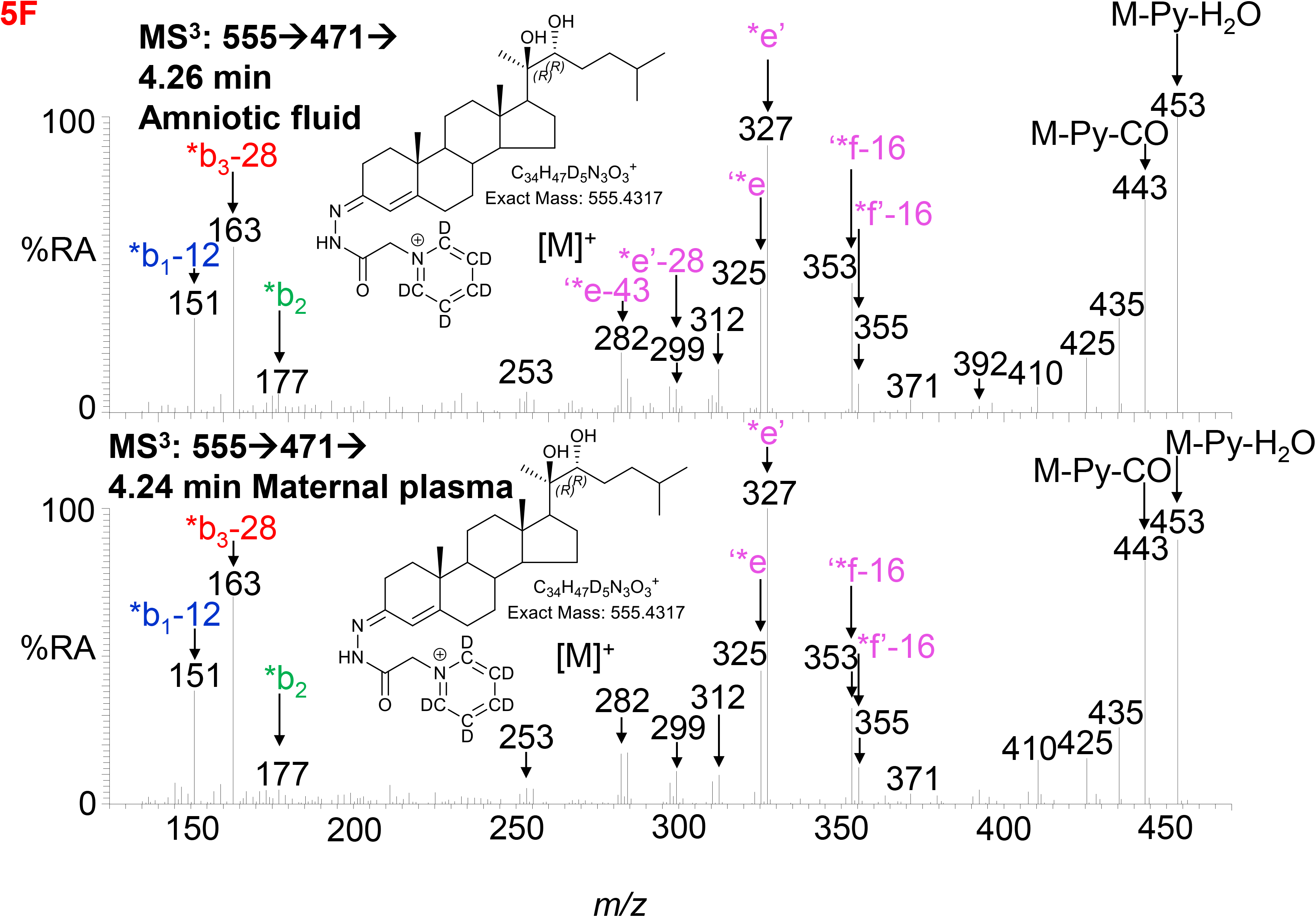
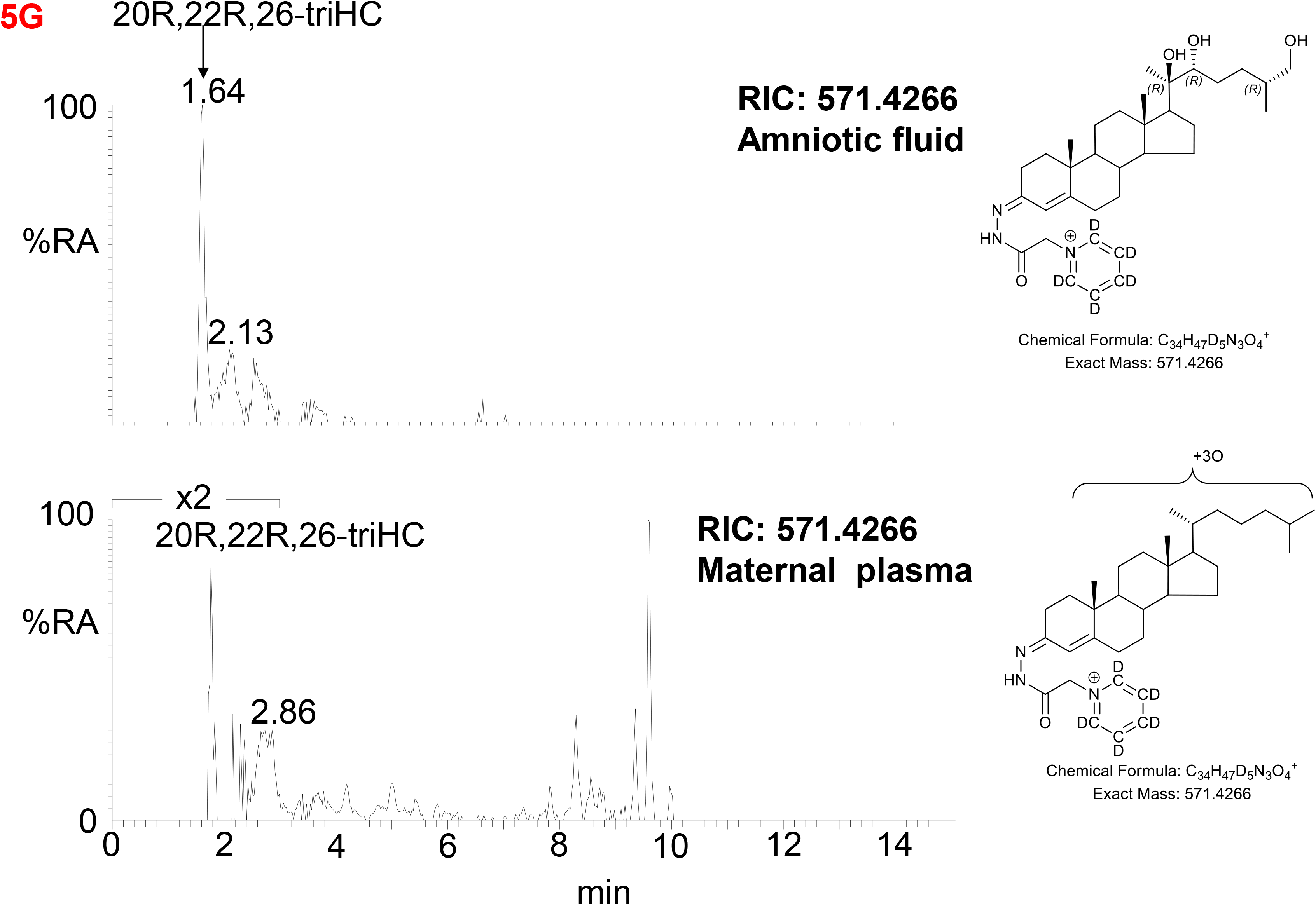
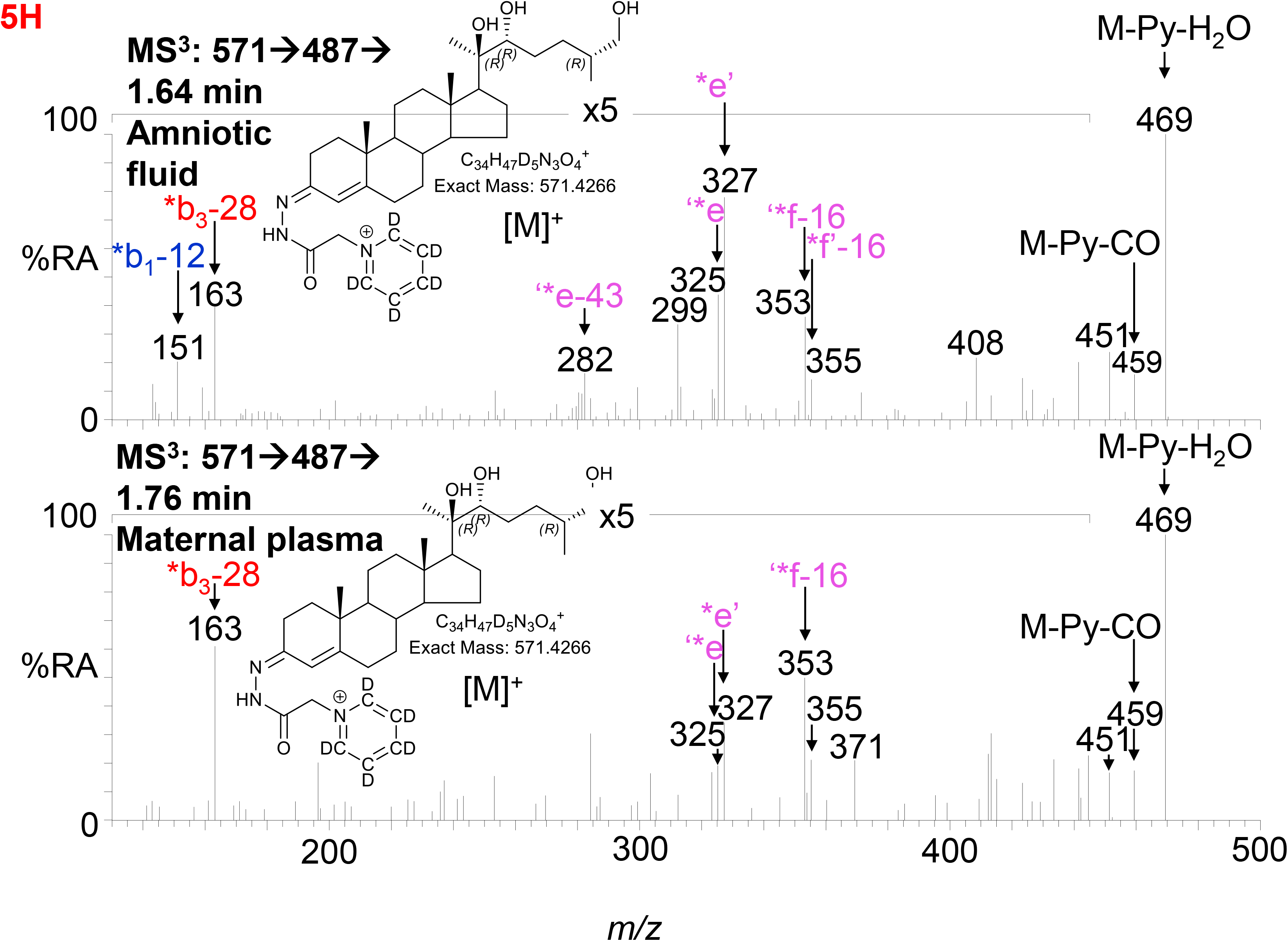
LC-MS(MS^n^) analysis of GP-derivatised oxysterols in amniotic fluid and plasma from pregnant females (maternal plasma). (A) RICs for monohydroxycholesterols (*m/z* 539.4368 ± 5 ppm) in amniotic fluid (upper panel), and in plasma from pregnant females (lower panel). (B) MRM-like ([M]^+^→[M- Py]^+^→355) chromatograms targeting 22-HC in amniotic fluid (upper panel) and in plasma from pregnant females (lower panel). (C) MS^3^ ([M]^+^→[M-Py]^+^→) spectra of 22R-HC identified in amniotic fluid (upper panel) and in plasma from pregnant females (lower panel). (D) RIC (*m/z* 555.4317 ± 5 ppm) corresponding to the [M]^+^ ion of dihydroxycholesterols identified in amniotic fluid (upper panel) and in plasma from pregnant females (lower panel). (E) MRM-like chromatograms targeting 20R,22R-diHC ([M]^+^→[M-Py]^+^→327) in amniotic fluid (upper panel) and in plasma from pregnant females (lower panel). (F) MS^3^ ([M]^+^→[M-Py]^+^→) spectra of 20R,22R-diHC in amniotic fluid (upper panel) and in plasma from pregnant females (lower panel). (G) RICs for trihydroxycholesterols (*m/z* 571.4266 ± 5 ppm) in amniotic fluid (upper panel) and in plasma from pregnant females (lower panel). (H) MS^3^ ([M]^+^→[M-Py]^+^→) spectra of the trihydroxycholesterols found in amniotic fluid (upper panel) and in plasma from pregnant females (lower panel) commensurate with the 20R,22R,26-triHC structure.

**Figure 6.**
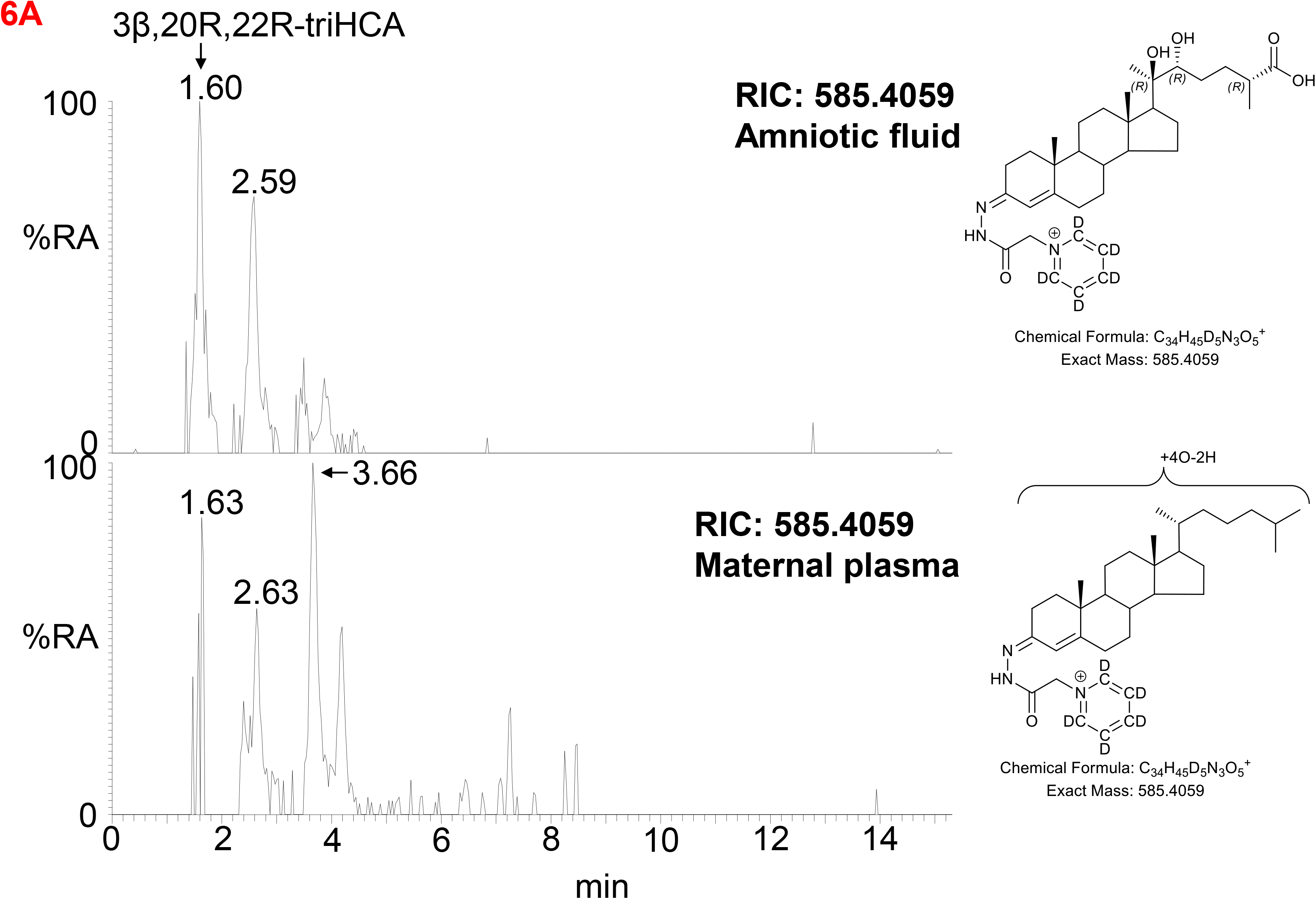
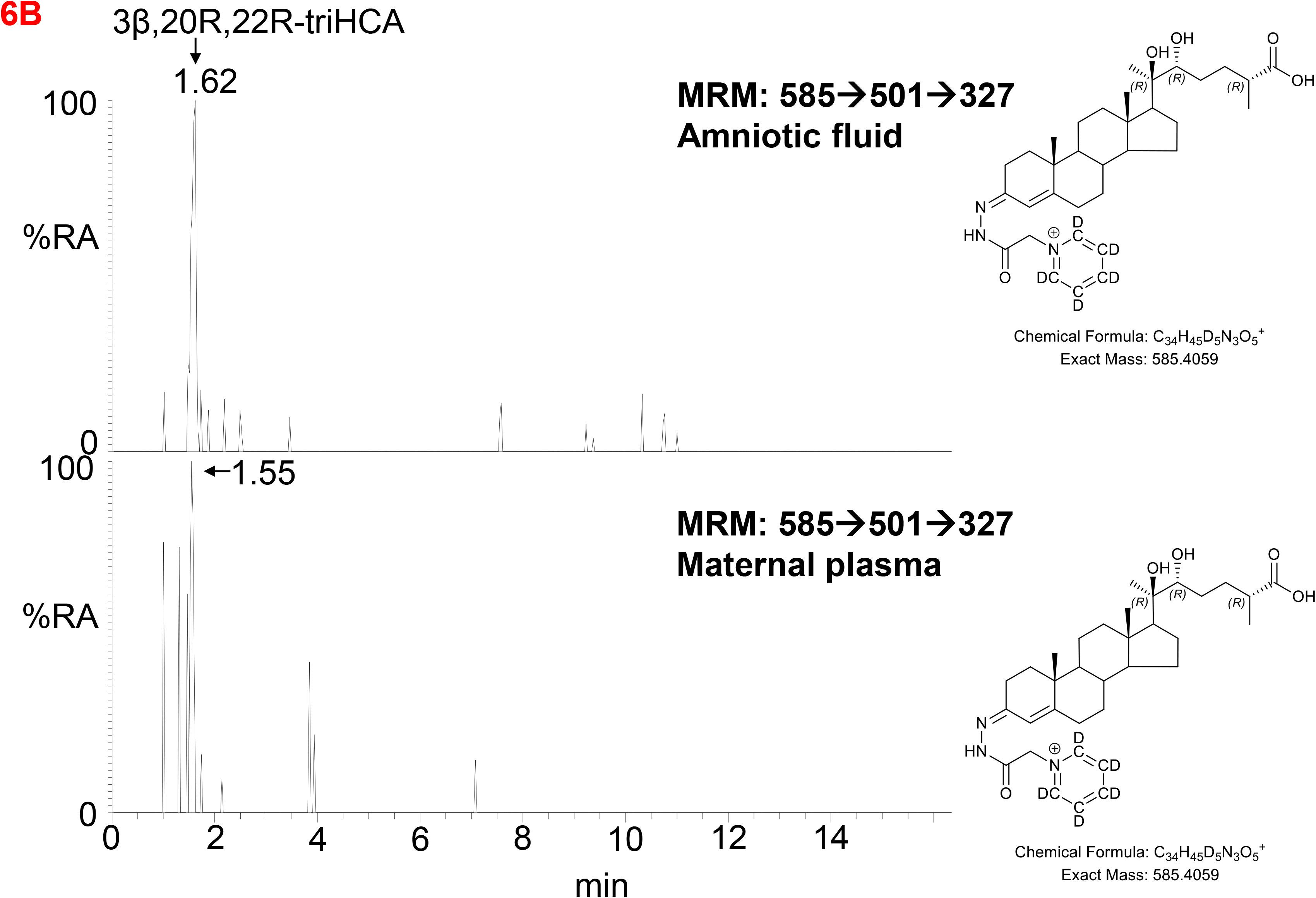
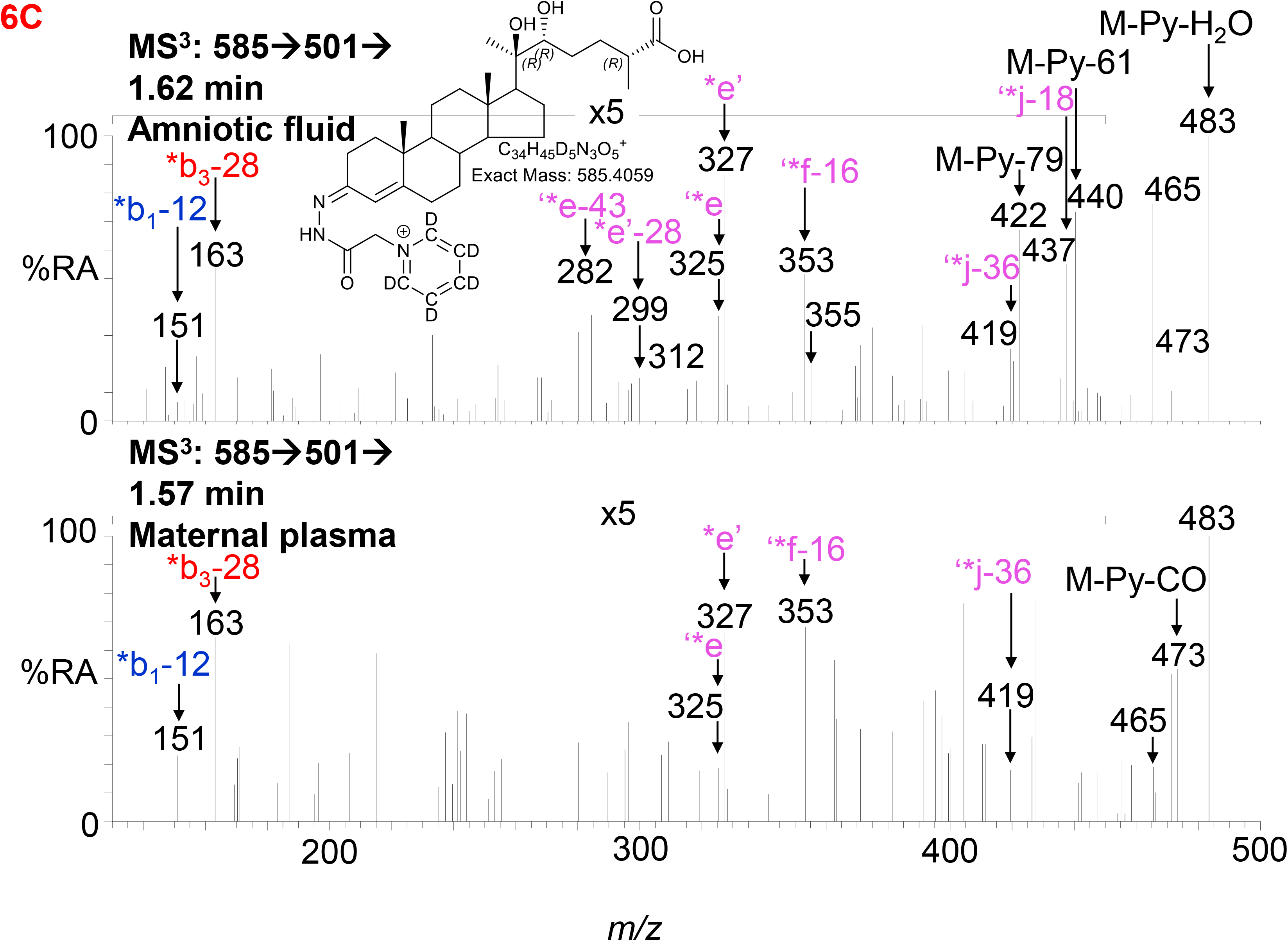
LC-MS(MS^n^) analysis of GP-derivatised trihydroxycholestenoic acids found in amniotic fluid and plasma from pregnant females (maternal plasma). (A) RICs (*m/z* 585.4059 ± 5 ppm) corresponding to the [M]^+^ ion of trihydroxycholestenoic acids found in amniotic fluid (upper panel) and in plasma from pregnant females (lower panel). (B) MRM-like ([M]^+^→[M-Py]^+^→327) chromatograms targeting 3β,20R,22R-triHCA in amniotic fluid and in plasma from pregnant females. (C) MS^3^ ([M]^+^→[M-Py]^+^→) spectra postulated to correspond to 3β,20R,22R-tiHCA in amniotic fluid (upper panel) and in plasma from pregnant females (lower panel).

Besides the oxysterols discussed above, the profile of cord and maternal plasma was investigated for other oxysterols and sterol acids routinely found in adult plasma, this data is included in Table S1. While it was possible to make quantitative measurements for mono- and di-hydroxycholesterols thanks to the availability of authentic standards, the absence of standards for 20R,22R,26-triHC and 3β,20R,22R-diHCA means that the values determined for these metabolites are semi-quantitative, but by using the same internal standard for quantification across samples should give reliable measurements for relative quantification across the sample groups.

### 3.3. Identification of Oxysterols in Amniotic Fluid

Amniotic fluid is derived from maternal plasma but also contains progressively more fetal urine as pregnancy continues. One of its functions is to facilitate the exchange of biochemicals between mother and fetus. Based on the data from analysis of placenta and cord plasma, it is reasonable to expect to find 22R-HC and its down-stream metabolites in amniotic fluid. As in cord and maternal blood 22R-HC (Figure 5A – C), 20R,22R-diHC (Figure 5D - F), 20R,22R,26-triHC (Figure 5G – H) and 3β,20R,22R-triHCA (Figure 6A – C) were identified and quantified in amniotic fluid (Table 1). Presumptively identified 3β,20R,22R-triH-Δ^5^-CA was also detected but not quantified (Supplemental Figure S5C & 5D).

### 3.4. Relative Quantification Between Samples Groups

As mentioned above although the measurement of some of the metabolites of 22R-HC is only semi-quantitative, relative values are likely to be accurate (Table 1). Shown in Figure 7 are plots displaying the quantities determined in the different samples for 22R-HC and its dominant metabolites. With respect to these metabolites, cord plasma is very different to control plasma from non-pregnant females with statistical differences also observed between cord and maternal plasma for 20R,22R- diHC and 3β,20R,22R-triHCA.

**Figure 7.**
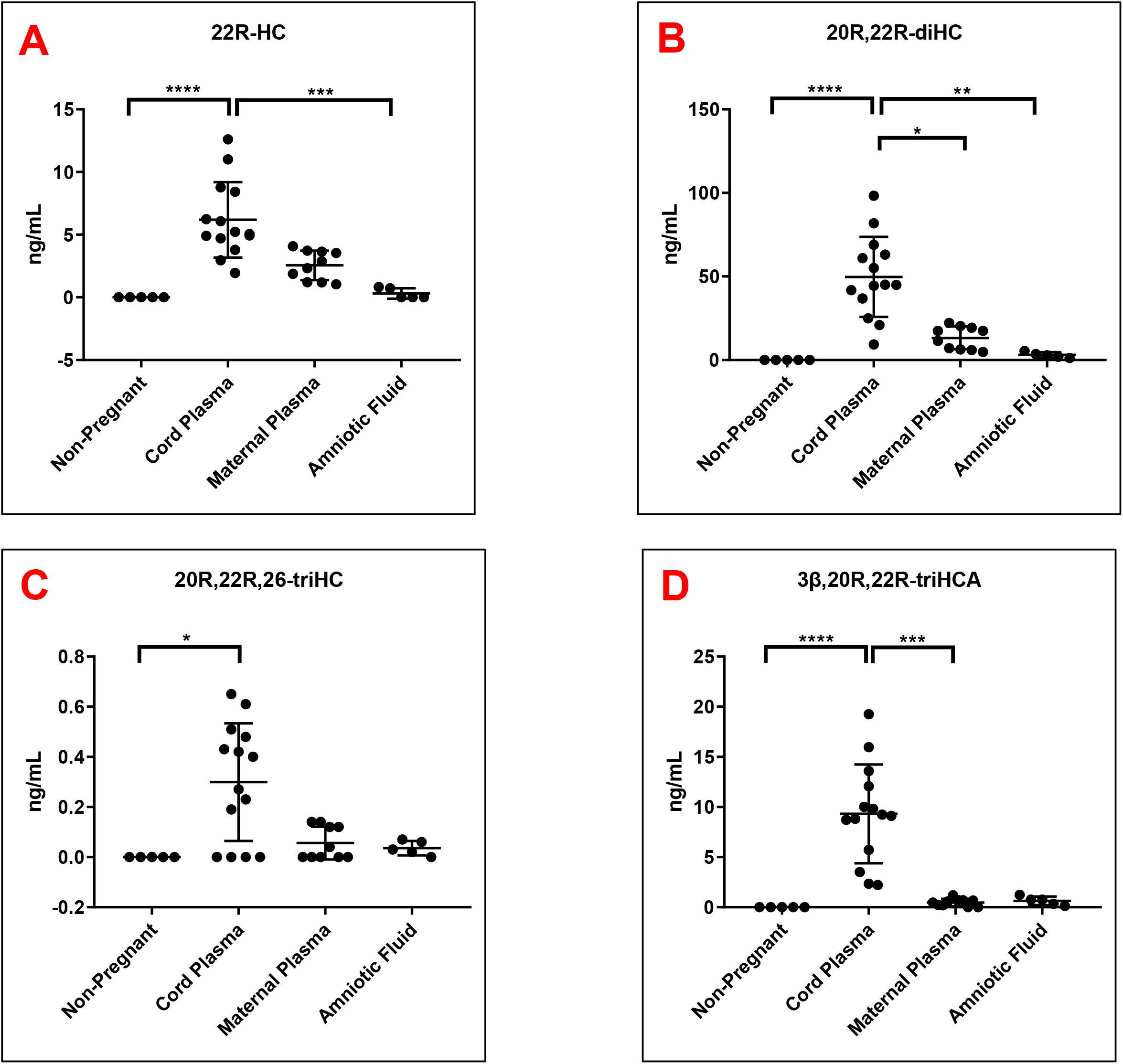
Concentration of 22R-HC and downstream metabolites. For each sample type: Control non- pregnant female plasma (plasma, n = 5); cord plasma (n = 14); maternal (pregnant female) plasma (n = 10); and amniotic fluid (n = 5). Concentrations of (A) 22R-HC, (B) 20R,22R-diHC, (C) 20R,22R,26-triHC and (D) 3β,20R,22R-triHCA. Concentrations were determined by LC-MS exploiting charge-tagging utilising the GP reagent. Vales in (A) and (B) are quantitative, those in (C) and (D) are semi-quantitative. The band represents the median where the whiskers extend to the most extreme upper and lower data points which are no more than 1.5 times the range between the first and third quartile. Non- parametric Kruskal-Wallis multiple comparisons test was used for comparison of data. *, P<0.05; **, P<0.01, ***P<0.001.

### 3.5. Quantification of Other Oxysterols

Besides the oxysterols discussed in the previous section, other oxysterols typically measured by the charge-tagging approach were also measured and are presented in Table S1 (11, 14, 19). Note the values for the oxysterols and sterol acids for which there is no authentic standard were quantified against [^2^H_7_]24R/S-HC and are only semi-quantitative values.

## 4. Discussion

In the current study we have investigated the oxysterol profile of placenta, cord plasma, maternal plasma, non-pregnant female plasma (control plasma) and amniotic fluid. In each of the pregnancy samples we identify metabolites derived from CYP11A1 which are essentially absent from non-pregnant females (Figure 7, Table 1). There are two significant findings from the current study. Firstly, the rediscovery of 20S-HC and the discovery of 22S-HC in human placenta (7), and secondly the uncovering of a shunt pathway for 22R-HC metabolism to C_27_ bile acids.

20S-HC is a controversial oxysterol as it has been detected in very few analytical studies (7, 8, 28, 29) despite being biologically active *in vitro*. 20S-HC, like 22R-HC, is a ligand to the liver X receptors α and β (LXRα, LXRβ) (30) and to the retinoic acid receptor-related orphan receptor γ (RORγ) (31), but unlike 22R-HC, activates the G protein-coupled receptor (GPCR) Smoothened (SMO), a key protein in the hedgehog signalling pathway, required for proper cell differentiation in the embryo (32, 33). 20S-HC also inhibits the processing of SREBP-2 (sterol regulatory element-binding protein 2) to its active form as the master transcription factor regulating cholesterol biosynthesis (34, 35), presumably by binding to INSIG (insulin induced gene) in a manner similar to other side-chain hydroxycholesterols (36). Recently, 20S-HC has been identified as a ligand to the sigma 2 (σ2) receptor (37), also known as transmembrane protein 97 (Tmem97), which is expressed in the central nervous system (38), and has been suggested to be a chaperone protein for NPC1 (Niemann Pick C1), the lysosomal cholesterol transport protein (37). The enzyme required to biosynthesise 20S-HC has not been identified, although CYP11A1 has been reported to generate both 20-hydroxyvitamin D_3_ and 20,22-dihydroxyvitamin D_3_ or 20,23-dihydroxyvitamin D_3_ from vitamin D_3_ (39, 40). The high level of CYP11A1 in placenta (17), makes this a good candidate enzyme for biosynthesis of 20S-HC. Like 20S-HC, there are few reports of the detection of 22S-HC in biological systems (29), however, 22S-HC has been identified as the sulphate ester in human meconium (10), the earliest stool of a mammalian infant, and in the human cell lines HCT-15 and HCT-116 (41). Unlike 20S-HC and most other side-chain oxysterols, 22S-HC is not an LXR agonist (42), behaving more like an antagonist (43), neither does it activate the Hh signalling pathway through SMO (32).

22R-HC and 20R,22R-diHC are abundant oxysterols in cord plasma and placenta. 20R,22R-diHC, like 22R-HC and 20S-HC, is an LXR ligand and all three appears to have similar activating capacity (44). Although the primary function of the LXRs is considered to be the regulation of cellular cholesterol (45), LXRs also appear to have developmental functions, being required for the development of dopaminergic neurons in midbrain (46). In fact, LXRβ also appears to have a protective role towards dopaminergic neurons, as the synthetic agonist GW3965 protects against the loss of dopaminergic neurons in a Parkinson’s disease mouse model (47).

CYP11A1 is an inner mitochondrial membrane protein and catalyses the side-chain cleavage of cholesterol to pregnenolone. The intermediates in this reaction scheme i.e. 22R-HC and 20R,22R-diHC, bind more tightly to CYP11A1 and are converted to pregnenolone at a greater rate than cholesterol (48). It is generally considered that 22R-HC and 20R,22R-diHC remain in the active site until all three oxidation steps are complete (3), however, the abundance of 20R-HC and of 20R,22R-diHC observed in cord plasma and placenta in this study would argue that this is not always the case. Pregnenolone is converted to progesterone by HSD3B1 which is localised in both mitochondria and the endoplasmic reticulum (49, 50), and is highly expressed in placenta (51). Progesterone has many roles associated with the establishment and maintenance of pregnancy, including ovulation, uterine and mammary gland development and the onset of labour (52). Progesterone suppresses spontaneous uterine contractility during pregnancy and, in most mammals, a fall in systemic progesterone is required for the initiation of labour at term. However, in humans, labour occurs in the presence of elevated circulating levels of progesterone. Despite this, disruption of progesterone signalling by the progesterone receptor (PR) antagonist RU486 at any stage of pregnancy results in myometrial contractions and labour, strongly suggesting that reduced progesterone signalling is responsible for labour in women (53). In the current study we have uncovered a shunt pathway that operates in parallel to pregnenolone/progesterone biosynthesis in the placenta (Figure 8). Beyond 20R,22R-diHC we identified three trihydroxycholesterol isomers, one of which gives an MS^3^ fragmentation pattern consistent with 20R,22R,26-triHC, and two dihydroxycholestenoic acid isomers one of which gives a fragmentation pattern we assign to 3β,20R,22R-triHCA. A down-stream metabolite 3β,20R,22R-triH- Δ^24^-CA was also presumptively identified. We awaited the chemical synthesis of these metabolites to definitively confirm their identification but their presence during pregnancy would define a new pathway of C_27_ bile acid biosynthesis (Figure 8). Most of the metabolites of this pathway are also observed in cord plasma, maternal plasma and amniotic fluid (Table 1). Notably, the amniotic fluid samples were from 16 – 18 weeks of gestation and the other pregnancy samples 37+ weeks indicating a pathway operational throughout pregnancy. CYP27A1 is the likely sterol hydroxylase which will convert 20R,22R-diHC to 20R,22R,26-triHC and on to 3β,20R,22R-triHCA. Like CYP11A1, CYP27A1 is an inner mitochondrial membrane enzyme and is expressed in placenta (6, 24). Although the C_27_ bile acid 3β,20R,22R-diHCA has not previously been identified, a C_27_ bile acid with 22R-hydroxylation has been identified in a patient with Zellweger’s syndrome (54). It should be noted that 20S-HC will also act as a substrate for pregnenolone formation via CYP11A1 catalysed reactions (29), presumably via 20R,22R-diHC (55). Thus, a potential route for 20S-HC metabolism is through 20R,22R-diHC and on to C_27_ bile acids.

**Figure 8.**
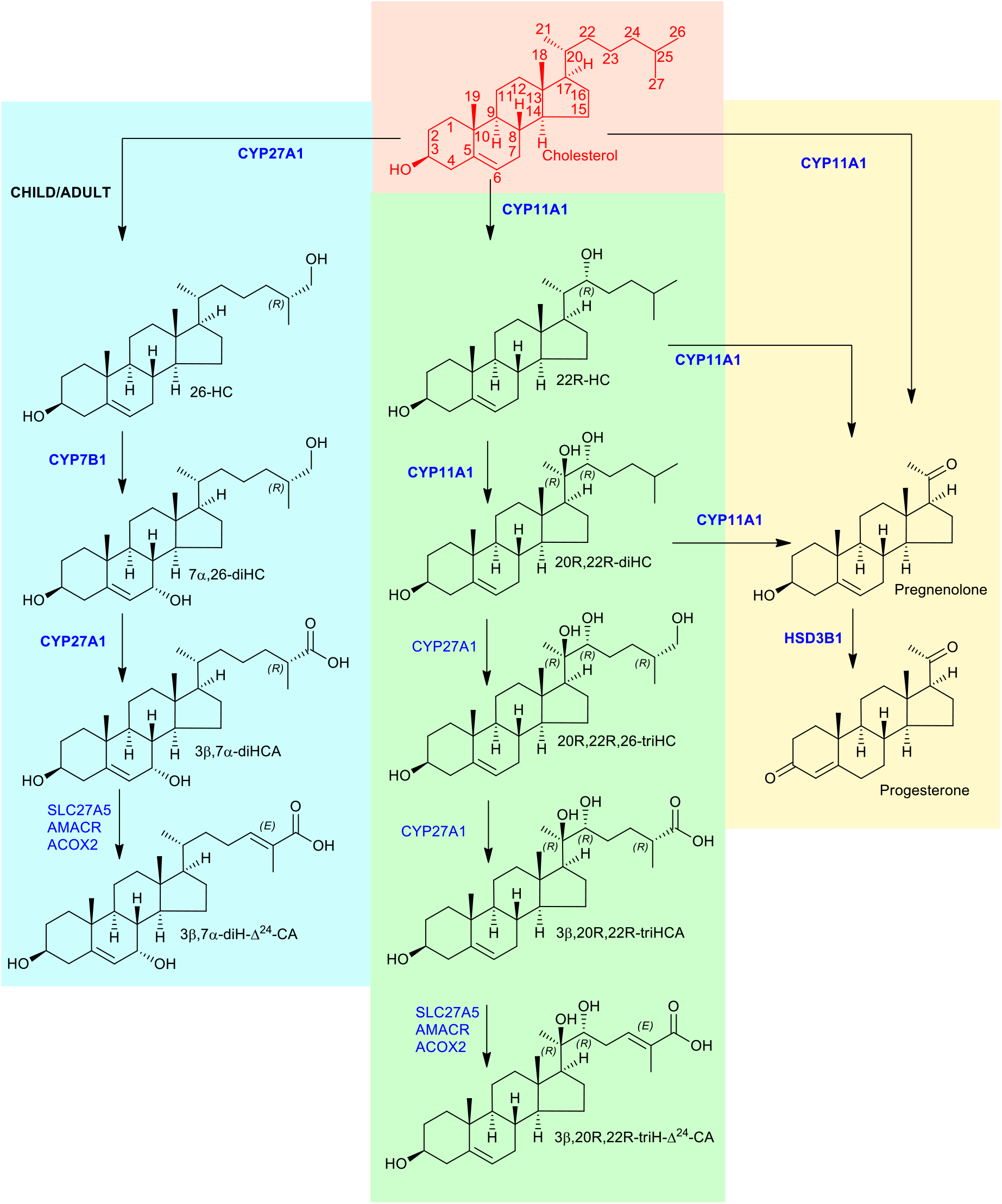
Proposed metabolism of cholesterol to the C_27_ bile acid 3β,20R,22R-triH-Δ^24^-CA (green background). For comparison the acidic pathway to the C_27_ bile acid 3β,7α-diH-Δ^24^-CA is shown (blue background) as is the pathway to progesterone (yellow background). Enzymes are indicated in blue. When shown in bold the enzymatic reactions can be found in the literature, when shown in normal typeface they are postulated. Cholesterol is shown in red with full stereochemistry and numbering system on a salmon background. For simplicity the 3β,20R,22R-triH-Δ^24^-CA and 3β,7α-diH-Δ^24^-CA are shown as the acids rather than the CoA-thioesters.

How important is the 20R,22R-diHC shunt pathway? At present we can only speculate, but the LXR- activating capacity of 22R-HC and 20R,22R-diHC and the expression of both LXRα and β during mammalian development makes it tempting to speculate that 20R-HC, 20R,22R-diHC and also 20S-HC by activating LXR, and in the case of 20S-HC by binding to Smo and Tmem97, are important for development of the embryo (Figure 9). In this regard, biallelic variants of Tmem97 were recently associated with fetal abnormalities (56). Little is known about the biological activities of trihydroxycholesterols and trihydroxycholestenoic acids and it is unknown whether 20R,22R,26-triHC, 3β,20R,22R-triHCA and 3β,20R,22R-triH-Δ^24^-CA are simply inactive intermediates on the road to bile acids or biologically active molecules themselves. A final point of note, during the course of this study we found evidence that HSD3B1 can oxidise sterols as well as steroids. Further details will be reported elsewhere.

**Figure 9.**
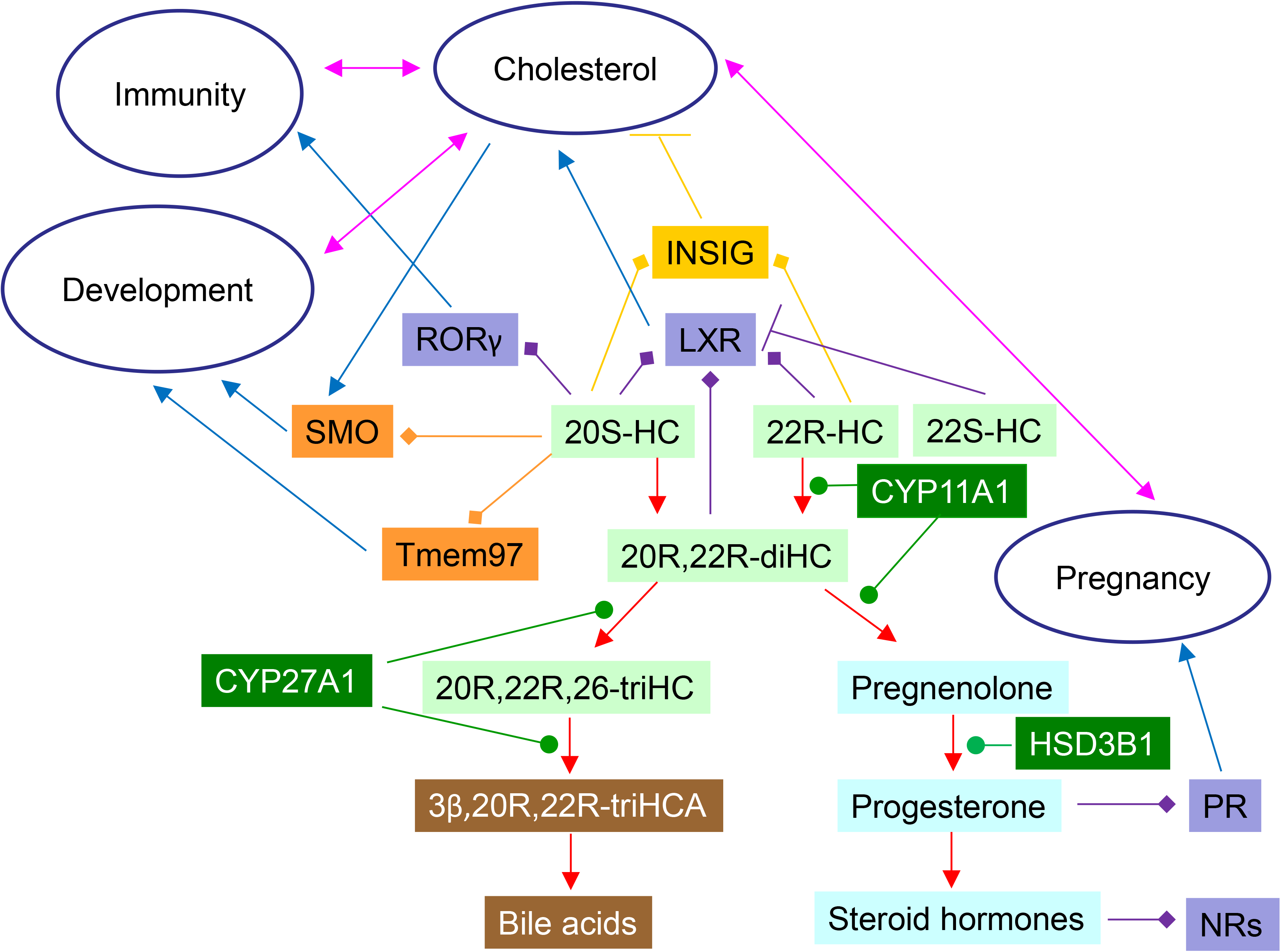
Schematic showing cholesterol metabolites identified in placenta and their interactions with protein receptors. 22R-HC, 22S-HC and 20S-HC are formed from cholesterol, 22R-HC by CYP11A1 which may also be the enzyme that catalyses the formation of 20S-HC. Oxysterols are shown on a light green background, steroids on a light blue background, bile acids on a brown background and enzymes on a dark green background. Nuclear receptors are shown on a purple background, GPCRs on an orange background and INSIG on a mustard background. Blue arrows indicate a “process”, red arrows a chemical reaction, T signifies inhibition of a process, arrows with a diamond arrowhead indicate activation of a receptor, and green oval arrowheads indicate catalysis. Pink double headed arrows link processes.

## Acknowledgements

This work was supported by the Biotechnology and Biological Sciences Research Council (BBSRC, grant numbers BB/I001735/1, BB/N015932/1 and BB/S019588/1 to WJG, BB/L001942/1 to YW), the European Union through European Structural Funds (ESF), as part of the Welsh Government funded Academic Expertise for Business project (to WJG and YW). ALD was supported via a KESS2 award in association with Markes International from the Welsh Government and the European Social Fund. Dr Peter Grosshans and Steve Smith of Markes International are thanked for helpful discussions. Members of the European Network for Oxysterol Research (ENOR, https://www.oxysterols.net/) are thanked for informative discussions.

## Supplemental Figures

**Supplemental Figure S1.** (A) Schematic illustration of the EADSA process exemplified by 20R,22R-diHC, 7α,(25R)26-dihydroxycholesterol (7α,26-diHC) and 7α,(25R)26-dihydroxycholest-4-en-3-one (7α,26- diHCO). (B) *Z* (*syn*) and *E* (*anti*) conformers of the GP-derivative of 22R-HC. The insets show minimum energy structures calculated by Chem 3DPro (Perkin Elmer), lone pairs are in pink. (C) Characteristic MS^3^ fragmentation of GP-derivatised 3β-hydroxy-5-ene sterols. The top box shows sterol numbering and fragmentation nomenclature (57). An asterisk indicates that the pyridine ring has been lost. A prime to the left of the fragment describing letter, e.g. ‘*e, indicates that homolytic fragmentation proceeds with loss of an additional hydrogen atom from the ion. A prime to the right, e.g. *e’, indicates that homolytic fragmentation proceeds with the addition of a hydrogen atom to the fragment ion. The central box shows the major ring-fragment ions. The lower box shows the changed pattern of ring fragment ions when a hydroxy group is present on C-7. (D) GP-derivatised 24S-HC gives a major side-chain fragment ion at *m/z* 353.3 (‘*f) and a minor fragment ion at 325.2 (‘*e). See Supplemental Figure S3 for MS^3^ spectrum. Coloured arrows suggest possible mechanisms and the minimum energy structure, showing lone-pairs, of the intermediate ion at *m/z* 437.4 is shown in the inset. Square brackets indicate different excited states.

**Supplemental Figure S2.** Suggested mechanisms for the formation of side-chain fragment ions in GP-derivatised 20-, and 22-hydroxysterols. (A) 22R-HC. (B) 20S-HC. (C-D) 20R,22R-diHC. (E) 22S,23-diHC. (F) 20R,22R,23-triHC. (G) 20R,22R,24-triHC. (H-I) 20R,22R,26-triHC. (J-L) 3β,20R,22R-triHCA. (M-O) 3β,20R,22R-triH-Δ^24^-CA. Square brackets indicate different tautomers or excited states. Coloured arrows suggest possible mechanisms. Minimum energy structures, showing lone pairs, of key intermediate ions are shown in insets. Major side-chain fragmentations are driven by the stability of ketene, ketone or enol products. MS^3^ spectra can be found in Figures 1B (22R-HC), 1F (20S-HC), 2B (20R,22R-diHC), 2C (22,23-diHC), 2G (20R,22R,26-triHC), 2H (20R,22R,23-triHC & 20R,22R,24-triHC), 3B (3β,20R,22R-triHCA) and 3F (3β,20R,22R-triH-Δ^24^-CA).

**Supplemental Figure S3.** Partial resolution by LC-MS(MS^n^) of GP-derivatised 20S-HC from 24S-HC and other hydroxycholesterols in placental samples. (A) RIC using the extended chromatographic gradient (37 min) for 20S-HC, 24S-HC and 26-HC (*m/z* 539.4368 ± 5 ppm, upper panel). MRM-like chromatograms targeting 22S-HC ([M]^+^→[M-Py]^+^→327, 2^nd^ panel), 24S-HC ([M]^+^→[M-Py]^+^→353, 3^rd^ panel) and [^2^H_7_]24R/S-HC ([M]^+^→[M-Py]^+^→353, bottom panel). Note GP derivatives each give *syn* and *anti* conformers resulting in twin peaks. (B-C) MS^3^ ([M]^+^→[M-Py]^+^→) spectra of 24S-HC (upper panels) and [^2^H_7_]24S-HC (lower panels). Note deuterated compounds elute slightly earlier than their hydrogen equivalents. (D) MS^3^ ([M]^+^→[M-Py]^+^→) spectra of the 26-HC. (E) RIC using the 17 min chromatographic gradient for monohydroxycholesterols (*m/z* 539.4368 ± 5 ppm, upper panel) and TIC generated by MS^3^ ([M]^+^→[M-Py]^+^→, lower panel). MS^3^ ([M]^+^→[M-Py]^+^→) spectra of (F) 7β-HC and (G) 7α-HC. Chromatograms using the 17 min gradient were aligned to the 26-HC peak in the NIST SRM 1950 plasma as described for Figure 1. Spectra of authentic standards can be found in (19).

**Supplemental Figure S4.** LC-MS(MS^n^) of GP-derivatised cholestenoic acids in placenta. (A) RIC (*m/z* 553.4161 ± 5 ppm, upper panel) corresponding to the [M]^+^ ion of 3β-HCA and TIC generated by MS^3^ ([M]^+^→[M-Py]^+^→, lower panel). (B) MS^3^ ([M]^+^→[M-Py]^+^→) spectrum of 3β-HCA. Chromatograms using the 17 min gradient were aligned to the 26-HC peak in the NIST SRM 1950 plasma as described for Figure 1. Spectra of authentic standards can be found in (19).

**Supplemental Figure S5.** LC-MS(MS^n^) analysis of GP-derivatised trihydroxycholestadienoic acids found in cord plasma and amniotic fluid. (A) RIC (*m/z* 583.3902 ± 5 ppm, upper panel) appropriate to 3β,20R,22R-triH-Δ^24^-CA and MRM-like chromatograms targeting 20,22-dihydroxysterols, [M]^+^→[M- Py]^+^→327 (centre panel) and [M]^+^→[M-Py]^+^→353 (lower panel) from cord plasma. (B) MS^3^ ([M]^+^→[M- Py]^+^→) spectrum postulated to correspond to 3β,20R,22R-triH-Δ^24^-CA in cord plasma. (C) RIC (*m/z* 583.3902 ± 5 ppm, upper panel) appropriate to 3β,20R,22R-triH-Δ^24^-CA and MRM-like chromatogram targeting 20,22-dihydroxysterols [M]^+^→[M-Py]^+^→327 (lower panel) from amniotic fluid. (D) MS^3^ ([M]^+^→[M-Py]^+^→) spectrum postulated to correspond to 3β,20R,22R-triH-Δ^24^-CA in amniotic fluid.

## References

1. Woollett, L. A. 2008. Where does fetal and embryonic cholesterol originate and what does it do? Annu Rev Nutr 28: 97–114.

2. Chatuphonprasert, W., K. Jarukamjorn, and I. Ellinger. 2018. Physiology and Pathophysiology of Steroid Biosynthesis, Transport and Metabolism in the Human Placenta. Front Pharmacol 9: 1027.

3. Mast, N., A. J. Annalora, D. T. Lodowski, K. Palczewski, C. D. Stout, and I. A. Pikuleva. 2011. Structural basis for three-step sequential catalysis by the cholesterol side chain cleavage enzyme CYP11A1. J Biol Chem 286: 5607–5613.

4. Tuckey, R. C., and K. J. Cameron. 1993. Catalytic properties of cytochrome P-450scc purified from the human placenta: comparison to bovine cytochrome P-450scc. Biochim Biophys Acta 1163: 185–194.

5. Sun, Y., S. Kopp, J. Strutz, C. C. Gali, M. Zandl-Lang, E. Fanaee-Danesh, A. Kirsch, S. Cvitic, S. Frank, R. Saffery, I. Björkhem, G. Desoye, C. Wadsack, and U. Panzenboeck. 2018. Gestational diabetes mellitus modulates cholesterol homeostasis in human fetoplacental endothelium. Biochim Biophys Acta Mol Cell Biol Lipids 1863: 968–979.

6. Mistry, H. D., L. O. Kurlak, Y. T. Mansour, L. Zurkinden, M. G. Mohaupt, and G. Escher. 2017. Increased maternal and fetal cholesterol efflux capacity and placental CYP27A1 expression in preeclampsia. J Lipid Res 58: 1186–1195.

7. Lin, Y. Y., M. Welch, and S. Lieberman. 2003. The detection of 20S-hydroxycholesterol in extracts of rat brains and human placenta by a gas chromatograph/mass spectrometry technique. J Steroid Biochem Mol Biol 85: 57–61.

8. Roberg-Larsen, H., M. F. Strand, S. Krauss, and S. R. Wilson. 2014. Metabolites in vertebrate Hedgehog signaling. Biochem Biophys Res Commun 446: 669–674.

9. Gustafsson, J. A., and P. Eneroth. 1972. Steroids in meconium and faeces from newborn infants. Proc R Soc Lond B Biol Sci 180: 179–186.

10. Eneroth, P., and J. Gustafsson. 1969. Steroids in newborns and infants. Hydroxylated cholesterol derivatives in the steroid monosulphate fraction from meconium. FEBS Lett 3: 129–132.

11. Crick, P. J., T. William Bentley, J. Abdel-Khalik, I. Matthews, P. T. Clayton, A. A. Morris, B. W. Bigger, C. Zerbinati, L. Tritapepe, L. Iuliano, Y. Wang, and W. J. Griffiths. 2015. Quantitative charge- tags for sterol and oxysterol analysis. Clin Chem 61: 400–411.

12. Raselli, T., T. Hearn, A. Wyss, K. Atrott, A. Peter, I. Frey-Wagner, M. R. Spalinger, E. M. Maggio, A. W. Sailer, J. Schmitt, P. Schreiner, A. Moncsek, J. Mertens, M. Scharl, W. J. Griffiths, M. Bueter, A. Geier, G. Rogler, Y. Wang, and B. Misselwitz. 2019. Elevated oxysterol levels in human and mouse livers reflect nonalcoholic steatohepatitis. J Lipid Res 60: 1270–1283.

13. Griffiths, W. J., P. J. Crick, A. Meljon, S. Theofilopoulos, J. Abdel-Khalik, E. Yutuc, J. E. Parker, D. E. Kelly, S. L. Kelly, E. Arenas, and Y. Wang. 2019. Additional pathways of sterol metabolism: Evidence from analysis of Cyp27a1-/- mouse brain and plasma. Biochim Biophys Acta Mol Cell Biol Lipids 1864: 191–211.

14. Abdel-Khalik, J., E. Yutuc, P. J. Crick, J. A. Gustafsson, M. Warner, G. Roman, K. Talbot, E. Gray, W. J. Griffiths, M. R. Turner, and Y. Wang. 2017. Defective cholesterol metabolism in amyotrophic lateral sclerosis. J Lipid Res 58: 267–278.

15. Quehenberger, O., A. M. Armando, A. H. Brown, S. B. Milne, D. S. Myers, A. H. Merrill, S. Bandyopadhyay, K. N. Jones, S. Kelly, R. L. Shaner, C. M. Sullards, E. Wang, R. C. Murphy, R. M. Barkley, T. J. Leiker, C. R. Raetz, Z. Guan, G. M. Laird, D. A. Six, D. W. Russell, J. G. McDonald, S. Subramaniam, E. Fahy, and E. A. Dennis. 2010. Lipidomics reveals a remarkable diversity of lipids in human plasma. J Lipid Res 51: 3299–3305.

16. Phinney, K. W., G. Ballihaut, M. Bedner, B. S. Benford, J. E. Camara, S. J. Christopher, W. C. Davis, N. G. Dodder, G. Eppe, B. E. Lang, S. E. Long, M. S. Lowenthal, E. A. McGaw, K. E. Murphy, B. C. Nelson, J. L. Prendergast, J. L. Reiner, C. A. Rimmer, L. C. Sander, M. M. Schantz, K. E. Sharpless, L. T. Sniegoski, S. S. Tai, J. B. Thomas, T. W. Vetter, M. J. Welch, S. A. Wise, L. J. Wood, W. F. Guthrie, C. R. Hagwood, S. D. Leigh, J. H. Yen, N. F. Zhang, M. Chaudhary-Webb, H. Chen, Z. Fazili, D. J. LaVoie, L. F. McCoy, S. S. Momin, N. Paladugula, E. C. Pendergrast, C. M. Pfeiffer, C. D. Powers, D. Rabinowitz, M. E. Rybak, R. L. Schleicher, B. M. Toombs, M. Xu, M. Zhang, and A. L. Castle. 2013. Development of a Standard Reference Material for metabolomics research. Anal Chem 85: 11732–11738.

17. Chung, B. C., K. J. Matteson, R. Voutilainen, T. K. Mohandas, and W. L. Miller. 1986. Human cholesterol side-chain cleavage enzyme, P450scc: cDNA cloning, assignment of the gene to chromosome 15, and expression in the placenta. Proc Natl Acad Sci U S A 83: 8962–8966.

18. Griffiths, W. J., T. Hearn, P. J. Crick, J. Abdel-Khalik, A. Dickson, E. Yutuc, and Y. Wang. 2017. Charge-tagging liquid chromatography-mass spectrometry methodology targeting oxysterol diastereoisomers. Chem Phys Lipids 207: 69–80.

19. Yutuc, E., A. L. Dickson, M. Pacciarini, L. Griffiths, P. R. S. Baker, L. Connell, A. Öhman, L. Forsgren, M. Trupp, S. Vilarinho, Y. Khalil, P. T. Clayton, S. Sari, B. Dalgic, P. Höflinger, L. Schöls, W. J. Griffiths, and Y. Wang. 2021. Deep mining of oxysterols and cholestenoic acids in human plasma and cerebrospinal fluid: Quantification using isotope dilution mass spectrometry. Anal Chim Acta 1154: 338259.

20. Sidhu, R., H. Jiang, N. Y. Farhat, N. Carrillo-Carrasco, M. Woolery, E. Ottinger, F. D. Porter, J. E. Schaffer, D. S. Ory, and X. Jiang. 2015. A validated LC-MS/MS assay for quantification of 24(S)- hydroxycholesterol in plasma and cerebrospinal fluid. J Lipid Res 56: 1222–1233.

21. Stiles, A. R., J. Kozlitina, B. M. Thompson, J. G. McDonald, K. S. King, and D. W. Russell. 2014. Genetic, anatomic, and clinical determinants of human serum sterol and vitamin D levels. Proc Natl Acad Sci U S A 111: E4006–4014.

22. Griffiths, W. J., E. Yutuc, J. Abdel-Khalik, P. J. Crick, T. Hearn, A. Dickson, B. W. Bigger, T. Hoi-Yee Wu, A. Goenka, A. Ghosh, S. A. Jones, D. F. Covey, D. S. Ory, and Y. Wang. 2019. Metabolism of Non-Enzymatically Derived Oxysterols: Clues from sterol metabolic disorders. Free Radic Biol Med 144: 124–133.

23. Cali, J. J., and D. W. Russell. 1991. Characterization of human sterol 27-hydroxylase. A mitochondrial cytochrome P-450 that catalyzes multiple oxidation reaction in bile acid biosynthesis. J Biol Chem 266: 7774–7778.

24. Uhlen, M., L. Fagerberg, B. M. Hallstrom, C. Lindskog, P. Oksvold, A. Mardinoglu, A. Sivertsson, C. Kampf, E. Sjostedt, A. Asplund, I. Olsson, K. Edlund, E. Lundberg, S. Navani, C. A. Szigyarto, J. Odeberg, D. Djureinovic, J. O. Takanen, S. Hober, T. Alm, P. H. Edqvist, H. Berling, H. Tegel, J. Mulder, J. Rockberg, P. Nilsson, J. M. Schwenk, M. Hamsten, K. von Feilitzen, M. Forsberg, L. Persson, F. Johansson, M. Zwahlen, G. von Heijne, J. Nielsen, and F. Ponten. 2015. Proteomics. Tissue-based map of the human proteome. Science 347: 1260419.

25. Axelson, M., and J. Sjovall. 1990. Potential bile acid precursors in plasma--possible indicators of biosynthetic pathways to cholic and chenodeoxycholic acids in man. J Steroid Biochem 36: 631–640.

26. Russell, D. W. 2003. The enzymes, regulation, and genetics of bile acid synthesis. Annu Rev Biochem 72: 137–174.

27. Griffiths, W. J., and Y. Wang. 2020. Oxysterols as lipid mediators: Their biosynthetic genes, enzymes and metabolites. Prostaglandins Other Lipid Mediat 147: 106381.

28. Yutuc, E., R. Angelini, M. Baumert, N. Mast, I. Pikuleva, J. Newton, M. R. Clench, D. O. F. Skibinski, O. W. Howell, Y. Wang, and W. J. Griffiths. 2020. Localization of sterols and oxysterols in mouse brain reveals distinct spatial cholesterol metabolism. Proc Natl Acad Sci U S A 117: 5749–5760.

29. Schroepfer, G. J., Jr. 2000. Oxysterols: modulators of cholesterol metabolism and other processes. Physiol Rev 80: 361–554.

30. Lehmann, J. M., S. A. Kliewer, L. B. Moore, T. A. Smith-Oliver, B. B. Oliver, J. L. Su, S. S. Sundseth, D. A. Winegar, D. E. Blanchard, T. A. Spencer, and T. M. Willson. 1997. Activation of the nuclear receptor LXR by oxysterols defines a new hormone response pathway. J Biol Chem 272: 3137–3140.

31. Jin, L., D. Martynowski, S. Zheng, T. Wada, W. Xie, and Y. Li. 2010. Structural basis for hydroxycholesterols as natural ligands of orphan nuclear receptor RORgamma. Mol Endocrinol 24: 923–929.

32. Nachtergaele, S., L. K. Mydock, K. Krishnan, J. Rammohan, P. H. Schlesinger, D. F. Covey, and R. Rohatgi. 2012. Oxysterols are allosteric activators of the oncoprotein Smoothened. Nat Chem Biol 8: 211–220.

33. Nachtergaele, S., D. M. Whalen, L. K. Mydock, Z. Zhao, T. Malinauskas, K. Krishnan, P. W. Ingham, D. F. Covey, C. Siebold, and R. Rohatgi. 2013. Structure and function of the Smoothened extracellular domain in vertebrate Hedgehog signaling. Elife 2: e01340.

34. Goldstein, J. L., R. A. DeBose-Boyd, and M. S. Brown. 2006. Protein sensors for membrane sterols. Cell 124: 35–46.

35. Abrams, M. E., K. A. Johnson, S. S. Perelman, L. S. Zhang, S. Endapally, K. B. Mar, B. M. Thompson, J. G. McDonald, J. W. Schoggins, A. Radhakrishnan, and N. M. Alto. 2020. Oxysterols provide innate immunity to bacterial infection by mobilizing cell surface accessible cholesterol. Nat Microbiol 5: 929–942.

36. Radhakrishnan, A., Y. Ikeda, H. J. Kwon, M. S. Brown, and J. L. Goldstein. 2007. Sterol-regulated transport of SREBPs from endoplasmic reticulum to Golgi: oxysterols block transport by binding to Insig. Proc Natl Acad Sci U S A 104: 6511–6518.

37. Cheng, Y.-S., T. Zhang, X. Ma, S. Pratuangtham, G. C. Zhang, A. A. Ondrus, A. Mafi, B. Lomenick, J. J. Jones, and A. E. Ondrus. 2021. A proteome-wide map of 20(S)-hydroxycholesterol interactors in cell membranes. Nature Chemical Biology 17: 1271–1280.

38. Yu, W., and J. M. Baskin. 2021. There is a lock for every key. Nature Chemical Biology 17: 1214–1216.

39. Guryev, O., R. A. Carvalho, S. Usanov, A. Gilep, and R. W. Estabrook. 2003. A pathway for the metabolism of vitamin D_3_: Unique hydroxylated metabolites formed during catalysis with cytochrome P450scc (CYP11A1). Proceedings of the National Academy of Sciences 100: 14754–14759.

40. Tuckey, R. C., W. Li, J. K. Zjawiony, M. A. Zmijewski, M. N. Nguyen, T. Sweatman, D. Miller, and A. Slominski. 2008. Pathways and products for the metabolism of vitamin D3 by cytochrome P450scc. Febs j 275: 2585–2596.

41. Roberg-Larsen, H., M. F. Strand, A. Grimsmo, P. A. Olsen, J. L. Dembinski, F. Rise, E. Lundanes, T. Greibrokk, S. Krauss, and S. R. Wilson. 2012. High sensitivity measurements of active oxysterols with automated filtration/filter backflush-solid phase extraction-liquid chromatography-mass spectrometry. J Chromatogr A 1255: 291–297.

42. Spencer, T. A., D. Li, J. S. Russel, J. L. Collins, R. K. Bledsoe, T. G. Consler, L. B. Moore, C. M. Galardi, D. D. McKee, J. T. Moore, M. A. Watson, D. J. Parks, M. H. Lambert, and T. M. Willson. 2001. Pharmacophore Analysis of the Nuclear Oxysterol Receptor LXRα. Journal of Medicinal Chemistry 44: 886–897.

43. Tranheim Kase, E., B. Andersen, H. I. Nebb, A. C. Rustan, and G. Hege Thoresen. 2006. 22- Hydroxycholesterols regulate lipid metabolism differently than T0901317 in human myotubes. Biochimica et Biophysica Acta (BBA) - Molecular and Cell Biology of Lipids 1761: 1515–1522.

44. Forman, B. M., B. Ruan, J. Chen, G. J. Schroepfer, and R. M. Evans. 1997. The orphan nuclear receptor LXRα is positively and negatively regulated by distinct products of mevalonate metabolism. Proceedings of the National Academy of Sciences 94: 10588–10593.

45. Korach-André, M., and J. Gustafsson. 2015. Liver X receptors as regulators of metabolism. Biomol Concepts 6: 177–190.

46. Sacchetti, P., K. M. Sousa, A. C. Hall, I. Liste, K. R. Steffensen, S. Theofilopoulos, C. L. Parish, C. Hazenberg, L. A. Richter, O. Hovatta, J. A. Gustafsson, and E. Arenas. 2009. Liver X receptors and oxysterols promote ventral midbrain neurogenesis in vivo and in human embryonic stem cells. Cell Stem Cell 5: 409–419.

47. Dai, Y.-b., X.-j. Tan, W.-f. Wu, M. Warner, and J.-Å. Gustafsson. 2012. Liver X receptor β protects dopaminergic neurons in a mouse model of Parkinson disease. Proceedings of the National Academy of Sciences 109: 13112–13117.

48. Tuckey, R. C., and K. J. Cameron. 1993. Side-chain specificities of human and bovine cytochromes P-450scc. Eur J Biochem 217: 209–215.

49. Thomas, J. L., R. P. Myers, and R. C. Strickler. 1989. Human placental 3 beta-hydroxy-5-ene- steroid dehydrogenase and steroid 5----4-ene-isomerase: purification from mitochondria and kinetic profiles, biophysical characterization of the purified mitochondrial and microsomal enzymes. J Steroid Biochem 33: 209–217.

50. Chapman, J. C., J. R. Polanco, S. Min, and S. D. Michael. 2005. Mitochondrial 3 beta-hydroxysteroid dehydrogenase (HSD) is essential for the synthesis of progesterone by corpora lutea: An hypothesis. Reproductive Biology and Endocrinology 3: 11.

51. Simard, J., M. L. Ricketts, S. Gingras, P. Soucy, F. A. Feltus, and M. H. Melner. 2005. Molecular biology of the 3beta-hydroxysteroid dehydrogenase/delta5-delta4 isomerase gene family. Endocr Rev 26: 525–582.

52. Astle, S., D. M. Slater, and S. Thornton. 2003. The involvement of progesterone in the onset of human labour. Eur J Obstet Gynecol Reprod Biol 108: 177–181.

53. Nadeem, L., O. Shynlova, E. Matysiak-Zablocki, S. Mesiano, X. Dong, and S. Lye. 2016. Molecular evidence of functional progesterone withdrawal in human myometrium. Nat Commun 7: 11565.

54. Une, M., K. Tsujimura, K. Kihira, and T. Hoshita. 1989. Identification of (22R)-3 alpha,7 alpha,12 alpha,22- and (23R)-3 alpha,7 alpha,12 alpha,23-tetrahydroxy-5 beta-cholestanoic acids in urine from a patient with Zellweger’s syndrome. J Lipid Res 30: 541–547.

55. Lambeth, J. D., S. E. Kitchen, A. A. Farooqui, R. Tuckey, and H. Kamin. 1982. Cytochrome P- 450scc-substrate interactions. Studies of binding and catalytic activity using hydroxycholesterols. J Biol Chem 257: 1876–1884.

56. Al-Hamed, M. H., N. Alsahan, M. Tulbah, W. Kurdi, W. Ali, J. A. Sayer, and F. Imtiaz. 2020. Fetal Anomalies Associated with Novel Pathogenic Variants in TMEM94. Genes (Basel) 11.

57. Karu, K., M. Hornshaw, G. Woffendin, K. Bodin, M. Hamberg, G. Alvelius, J. Sjovall, J. Turton, Y. Wang, and W. J. Griffiths. 2007. Liquid chromatography-mass spectrometry utilizing multi-stage fragmentation for the identification of oxysterols. J Lipid Res 48: 976–987.

