## Supplemental for "Identification of Unusual Oxysterols Biosynthesised in Human Pregnancy by Charge-Tagging and Liquid Chromatography - Mass Spectrometry"

**Supplemental Material**

**Supplemental Materials and Methods**

*LC-MS Mobile Phase and Gradient*

LC was performed on an Ultimate 3000 UHPLC system (Dionex, now Thermo Fisher Scientific, Hemel Hempstead, UK) using a Hypersil GOLD revered phase column (1.9 µm particle size, 50 × 2.1 mm, Thermo Fisher). Mobile phase A consisted of 33.3% methanol, 16.7% acetonitrile and 0.1% formic acid. Mobile phase B consisted of 63.3% methanol, 31.7% acetonitrile and 0.1% formic acid. The 17 min chromatographic run started at 20% B for 1 min, before increasing the proportion of B to 80% over 7 minutes, and maintaining this for a further 5 min. The proportion of B was returned to 20% over 6 s and re-equilibration was for 3 min, 54 s to give a total run time of 17 min. To separate closely eluting oxysterols, the usual 17 min gradient was extended to 37 min. The mobile phase composition was initially at 20% B for 10 min, changed to 50% B over the next 10 min, maintained at this composition for 6 min, and then changed 80% B over the next 3 min. The mobile phase composition was held at 80% B for a further 3 min before returning to 20% B in 6 s and reconditioning the column for a further 4 min 54 s.

The flow rate was 200 µL/min and the eluent was directed to the atmospheric pressure ionization (API) source of an Orbitrap Elite MS (Thermo Fisher Scientific). The Orbitrap was calibrated externally, and the mass accuracy was better than 5 ppm.

*Scan Events on the Orbitrap Elite MS*

The scan events were as follows. For each LC-MS($MS^n$) injection the first scan event consisted of a Fourier Transform (FT)-MS scan in the Orbitrap at 120,000 resolution (full width at half-maximum height; FWHM) at *m/z* 400, simultaneous to which sequential $MS^3$ scan events were carried out in the linear ion-trap with normalised collision energies of 30 for $MS^2$ and 35 for $MS^3$ (instrument settings).

| Injection (run time) | Orbitrap *m/z* | LIT $MS^3$ | LIT $MS^3$ | LIT $MS^3$ | LIT $MS^3$ | LIT $MS^3$ |
|---|---|---|---|---|---|---|
| 1 (17 min) | 400-610 | 522.3→443.3→ | 527.4→443.3→ | 532.4→453.4→ | 537.4→453.4→ | |
| 2 (17 min) | 400-610 | 534.4→455.4→ | 539.4→455.4→ | 550.4→471.4→ | 555.4→471.4→ | |
| 3 (17 min) | 400-610 | 546.4→467.4→ | 551.4→467.3→ | 564.4→485.3→ | 569.4→485.3→ | |
| 4 (17 min) | 400-610 | 546.5→462.4→ | 548.4→469.3→ | 553.4→469.3→ | 580.4→501.3→ | 585.4→501.3→ |
| 5 (17 min) | 400-610 | 562.4→483.3→ | 566.4→487.4→ | 567.4→483.3→ | 571.4→487.4→ | |
| 6 (17 min) | 400-610 | 537.4→435.3→ | 543.5→441.4→ | 556.4→477.4→ | 561.5→477.4→ | |
| 7 (17 min) | 300-800 | 614.4→535.3→ | 619.4→535.3→ | 710.4→631.4→ | 715.5→631.4→ | |
| 8 (17 min) | 400-610 | 506.3→427.3→ | 511.4→427.3→ | 578.4→499.3→ | 583.4→499.3→ | |
| 9 (37 min) | 400-610 | 539.4→455.4→ | 546.5→462.4→ | 550.4→471.4→ | 555.4→471.4→ | |
| 10 (17 min) | 400-610 | 540.4→461.4→ | 541.5→462.4→ | 545.5→461.4→ | 546.5→462.4→ | |

**Supplemental Figures**

**Supplemental Figure S1.** (A) Schematic illustration of the EADSA process exemplified by 20R,22R-diHC, 7α,(25R)26-dihydroxycholesterol (7α,26-diHC) and 7α,(25R)26-dihydroxycholest-4-en-3-one (7α,26-diHCO). (B) *Z* (*syn*) and *E* (*anti*) conformers of the GP-derivative of 22R-HC. The insets show minimum energy structures calculated by Chem 3DPro (Perkin Elmer), lone pairs are in pink. (C) Characteristic $MS^3$ fragmentation of GP-derivatised 3β-hydroxy-5-ene sterols. The top box shows sterol numbering and fragmentation nomenclature (1). An asterisk indicates that the pyridine ring has been lost. A prime to the left of the fragment describing letter, e.g. '*e, indicates that homolytic fragmentation proceeds with loss of an additional hydrogen atom from the ion. A prime to the right, e.g. *e′, indicates that homolytic fragmentation proceeds with the addition of a hydrogen atom to the fragment ion. The central box shows the major ring-fragment ions. The lower box shows the changed pattern of ring fragment ions when a hydroxy group is present on C-7. (D) GP-derivatised 24S-HC gives a major side-chain fragment ion at *m/z* 353.3 (‘*f) and a minor fragment ion at 325.2 (‘*e). See Supplemental Figure S3 for $MS^3$ spectrum. Coloured arrows suggest possible mechanisms and the minimum energy structure, showing lone-pairs, of the intermediate ion at *m/z* 437.4 is shown in the inset. Square brackets indicate different excited states.

**Supplemental Figure S2.** Suggested mechanisms for the formation of side-chain fragment ions in GP-derivatised 20-, and 22-hydroxysterols. (A) 22R-HC. (B) 20S-HC. (C-D) 20R,22R-diHC. (E) 22S,23-diHC. (F) 20R,22R,23-triHC. (G) 20R,22R,24-triHC. (H-I) 20R,22R,26-triHC. (J-L) 3β,20R,22R-triHCA. (M-O) 3β,20R,22R-triH-$\Delta^{24}$-CA. Square brackets indicate different tautomers or excited states. Coloured arrows suggest possible mechanisms. Minimum energy structures, showing lone pairs, of key intermediate ions are shown in insets. Major side-chain fragmentations are driven by the stability of ketene, ketone or enol products. $MS^3$ spectra can be found in Figures 1B (22R-HC), 1F (20S-HC), 2B (20R,22R-diHC), 2C (22,23-diHC), 2G (20R,22R,26-triHC), 2H (20R,22R,23-triHC & 20R,22R,24-triHC), 3B (3β,20R,22R-triHCA) and 3F (3β,20R,22R-triH-$\Delta^{24}$-CA).

**Supplemental Figure S3.** Partial resolution by LC-MS($MS^n$) of GP-derivatised 20S-HC from 24S-HC and other hydroxycholesterols in placental samples. (A) RIC using the extended chromatographic gradient (37 min) for 20S-HC, 24S-HC and 26-HC (*m/z* 539.4368 ± 5 ppm, upper panel). MRM-like chromatograms targeting 22S-HC ($[M]^+$→$[M-Py]^+$→327, 2nd panel), 24S-HC ($[M]^+$→$[M-Py]^+$→353, 3rd panel) and [$^2H_7$]24R/S-HC ($[M]^+$→$[M-Py]^+$→353, bottom panel). Note GP derivatives each give *syn* and *anti* conformers resulting in twin peaks. (B-C) $MS^3$ ($[M]^+$→$[M-Py]^+$→) spectra of 24S-HC (upper panels) and [$^2H_7$]24S-HC (lower panels). Note deuterated compounds elute slightly earlier than their hydrogen equivalents. (D) $MS^3$ ($[M]^+$→$[M-Py]^+$→) spectra of the 26-HC. (E) RIC using the 17 min chromatographic gradient for monohydroxycholesterols (*m/z* 539.4368 ± 5 ppm, upper panel) and TIC generated by $MS^3$ ($[M]^+$→$[M-Py]^+$→, lower panel). $MS^3$ ($[M]^+$→$[M-Py]^+$→) spectra of (F) 7β-HC and (G) 7α-HC. Chromatograms using the 17 min gradient were aligned to the 26-HC peak in the NIST SRM 1950 plasma as described for Figure 1. Spectra of authentic standards can be found in (2).

**Supplemental Figure S4.** LC-MS($MS^n$) of GP-derivatised cholestenoic acids in placenta. (A) RIC (*m/z* 553.4161 ± 5 ppm, upper panel) corresponding to the $[M]^+$ ion of 3β-HCA and TIC generated by $MS^3$ ($[M]^+$→$[M-Py]^+$→, lower panel). (B) $MS^3$ ($[M]^+$→$[M-Py]^+$→) spectrum of 3β-HCA. Chromatograms using the 17 min gradient were aligned to the 26-HC peak in the NIST SRM 1950 plasma as described for Figure 1. Spectra of authentic standards can be found in (2).

**Supplemental Figure S5.** LC-MS($MS^n$) analysis of GP-derivatised trihydroxycholestadienoic acids found in cord plasma and amniotic fluid. (A) RIC (*m/z* 583.3902 ± 5 ppm, upper panel) appropriate to 3β,20R,22R-triH-$\Delta^{24}$-CA and MRM-like chromatograms targeting 20,22-dihydroxysterols, $[M]^+$→$[M-Py]^+$→327 (centre panel) and $[M]^+$→$[M-Py]^+$→353 (lower panel) from cord plasma. (B) $MS^3$ ($[M]^+$→$[M-Py]^+$→) spectrum postulated to correspond to 3β,20R,22R-triH-$\Delta^{24}$-CA in cord plasma. (C) RIC (*m/z* 583.3902 ± 5 ppm, upper panel) appropriate to 3β,20R,22R-triH-$\Delta^{24}$-CA and MRM-like chromatogram targeting 20,22-dihydroxysterols $[M]^+$→$[M-Py]^+$→327 (lower panel) from amniotic fluid. (D) $MS^3$ ($[M]^+$→$[M-Py]^+$→) spectrum postulated to correspond to 3β,20R,22R-triH-$\Delta^{24}$-CA in amniotic fluid.

1. Karu, K., M. Hornshaw, G. Woffendin, K. Bodin, M. Hamberg, G. Alvelius, J. Sjovall, J. Turton, Y. Wang, and W. J. Griffiths. 2007. Liquid chromatography-mass spectrometry utilizing multi-stage fragmentation for the identification of oxysterols. *J Lipid Res* **48**: 976-987.
2. Yutuc, E., A. L. Dickson, M. Pacciarini, L. Griffiths, P. R. S. Baker, L. Connell, A. Öhman, L. Forsgren, M. Trupp, S. Vilarinho, Y. Khalil, P. T. Clayton, S. Sari, B. Dalgic, P. Höflinger, L. Schöls, W. J. Griffiths, and Y. Wang. 2021. Deep mining of oxysterols and cholestenoic acids in human plasma and cerebrospinal fluid: Quantification using isotope dilution mass spectrometry. *Anal Chim Acta* **1154**: 338259.

**Figure S1A**

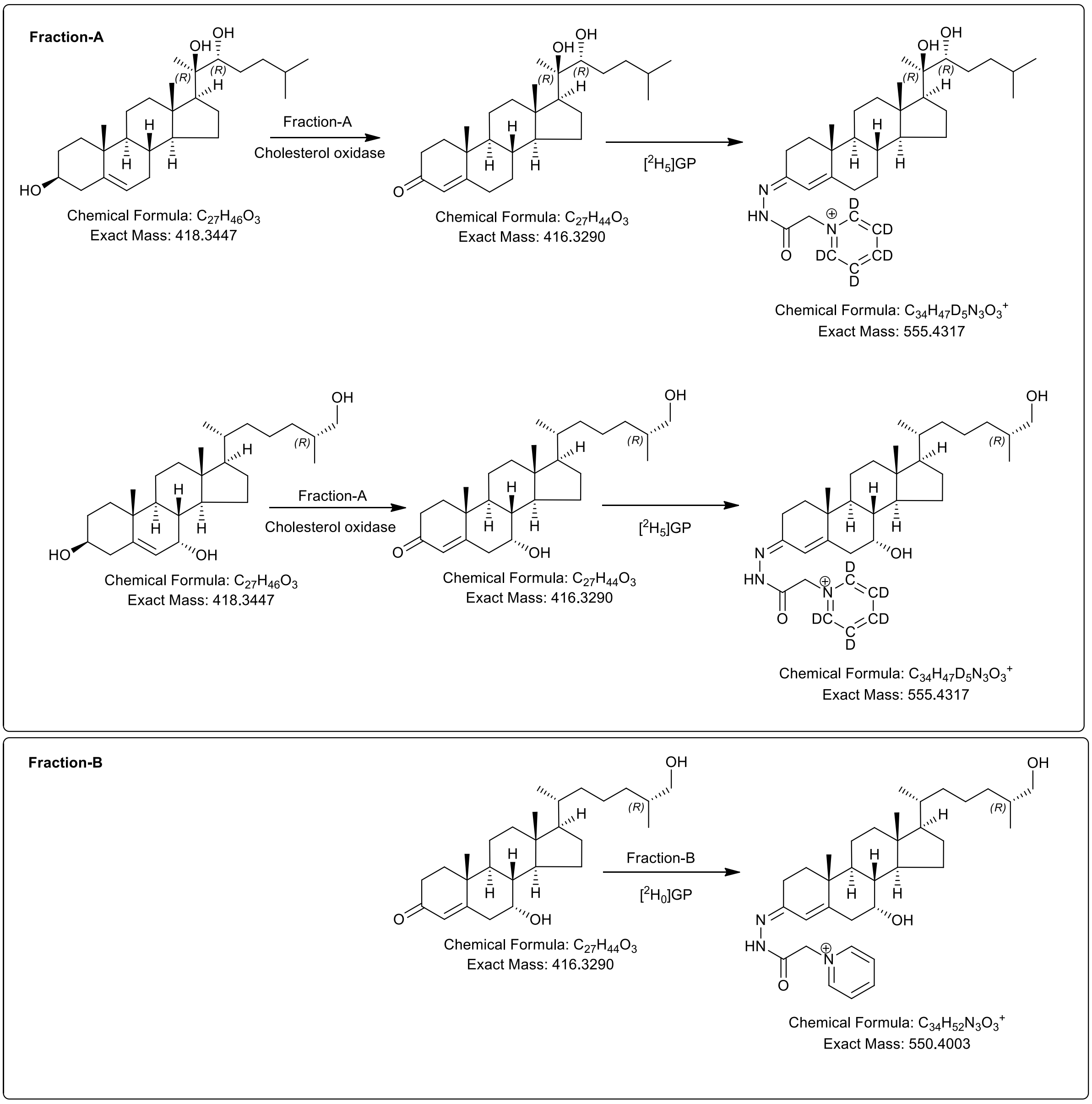
Fraction-A
OH
OH
(R)
(R)
H
HO
Chemical Formula: $C_{27}H_{46}O_3$
Exact Mass: 418.3447
Fraction-A
Cholesterol oxidase
Chemical Formula: $C_{27}H_{44}O_3$
Exact Mass: 416.3290
[$^2H_5$]GP
N
HN
O
D
DC
CD
Chemical Formula: $C_{34}H_{47}D_5N_3O_3^+$
Exact Mass: 555.4317
OH
Chemical Formula: $C_{27}H_{46}O_3$
Exact Mass: 418.3447
Fraction-A
Cholesterol oxidase
Chemical Formula: $C_{27}H_{44}O_3$
Exact Mass: 416.3290
[$^2H_5$]GP
Chemical Formula: $C_{34}H_{47}D_5N_3O_3^+$
Exact Mass: 555.4317
Fraction-B
Chemical Formula: $C_{27}H_{44}O_3$
Exact Mass: 416.3290
Fraction-B
[$^2H_0$]GP
Chemical Formula: $C_{34}H_{52}N_3O_3^+$
Exact Mass: 550.4003

**Figure S1B**

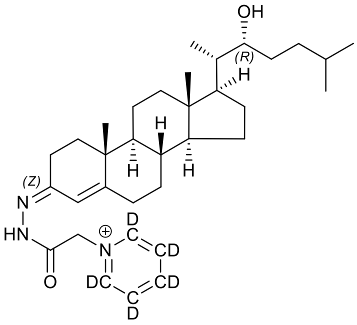

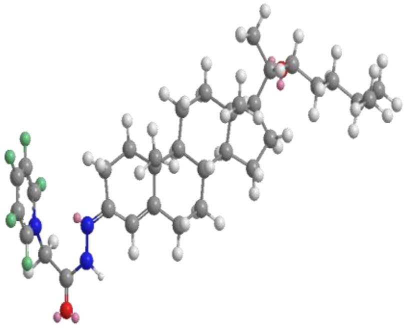

Chemical Formula: $C_{34}H_{47}D_5N_3O_2^+$
Exact Mass: 539.4368

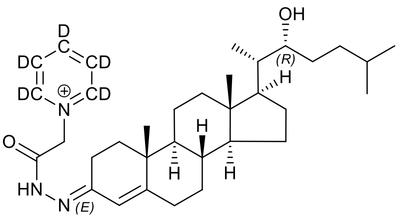

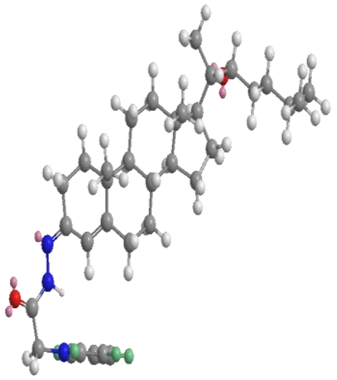

Chemical Formula: $C_{34}H_{47}D_5N_3O_2^+$
Exact Mass: 539.4368

**Figure S1C**

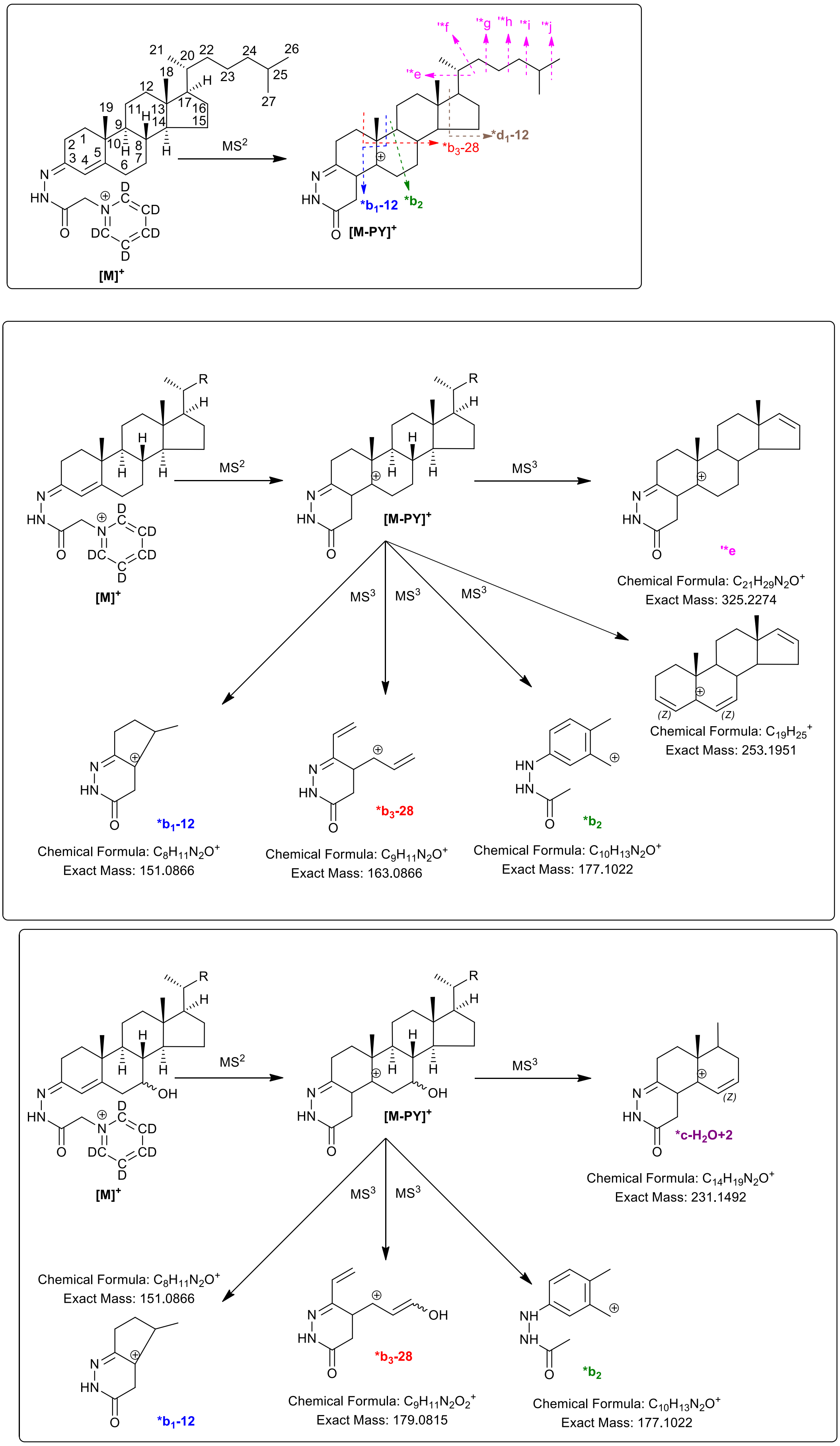
MS2
MS3
[M]+
[M-PY]+
*b1-12
*b2
*b3-28
*d1-12
**e
**f
**g
**h
**i
**j
*c-H2O+2
Chemical Formula: C21H29N2O+
Exact Mass: 325.2274
Chemical Formula: C19H25+
Exact Mass: 253.1951
Chemical Formula: C8H11N2O+
Exact Mass: 151.0866
Chemical Formula: C9H11N2O+
Exact Mass: 163.0866
Chemical Formula: C10H13N2O+
Exact Mass: 177.1022
Chemical Formula: C14H19N2O+
Exact Mass: 231.1492
Chemical Formula: C8H11N2O+
Exact Mass: 151.0866
Chemical Formula: C9H11N2O2+
Exact Mass: 179.0815
Chemical Formula: C10H13N2O+
Exact Mass: 177.1022

**Figure S1D**

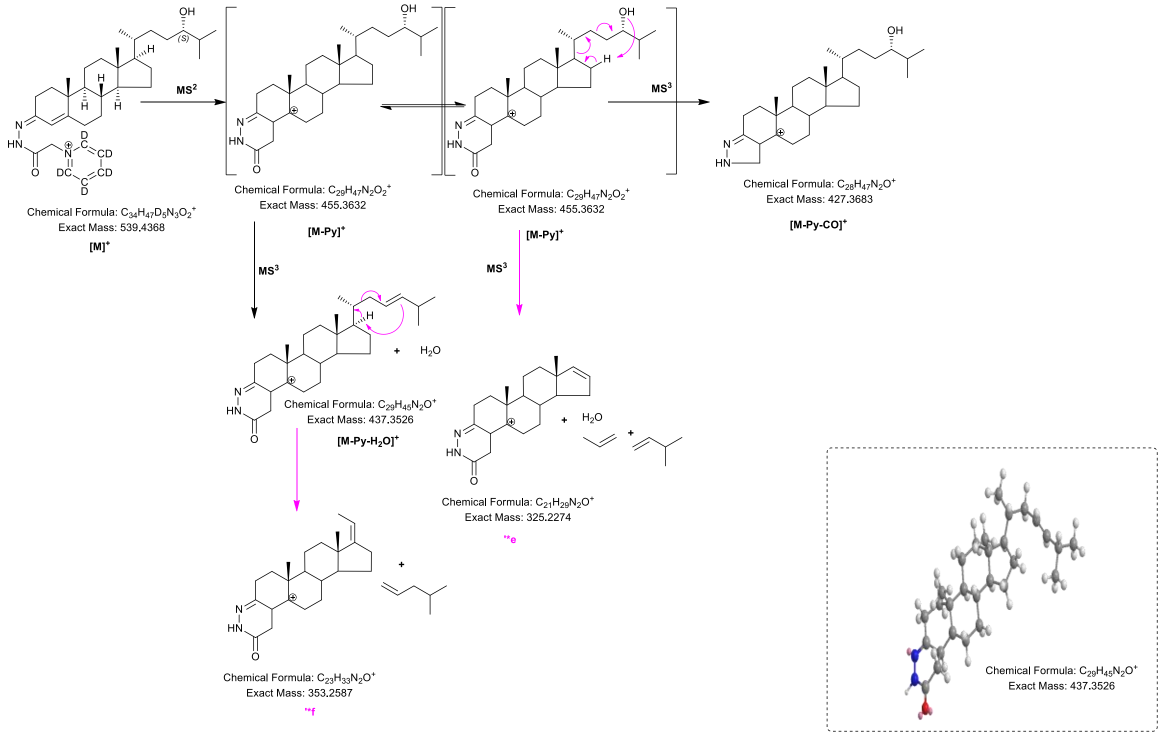

MS2
MS3
MS3
MS3
Chemical Formula: $C_{34}H_{47}D_5N_3O_2^+$
Exact Mass: 539.4368
[M]+
Chemical Formula: $C_{29}H_{47}N_2O_2^+$
Exact Mass: 455.3632
[M-Py]+
Chemical Formula: $C_{29}H_{47}N_2O_2^+$
Exact Mass: 455.3632
[M-Py]+
Chemical Formula: $C_{28}H_{47}N_2O^+$
Exact Mass: 427.3683
[M-Py-CO]+
Chemical Formula: $C_{29}H_{45}N_2O^+$
Exact Mass: 437.3526
[M-Py-H2O]+
Chemical Formula: $C_{21}H_{29}N_2O^+$
Exact Mass: 325.2274
**e
Chemical Formula: $C_{23}H_{33}N_2O^+$
Exact Mass: 353.2587
**f
Chemical Formula: $C_{29}H_{45}N_2O^+$
Exact Mass: 437.3526

**Figure S2A**

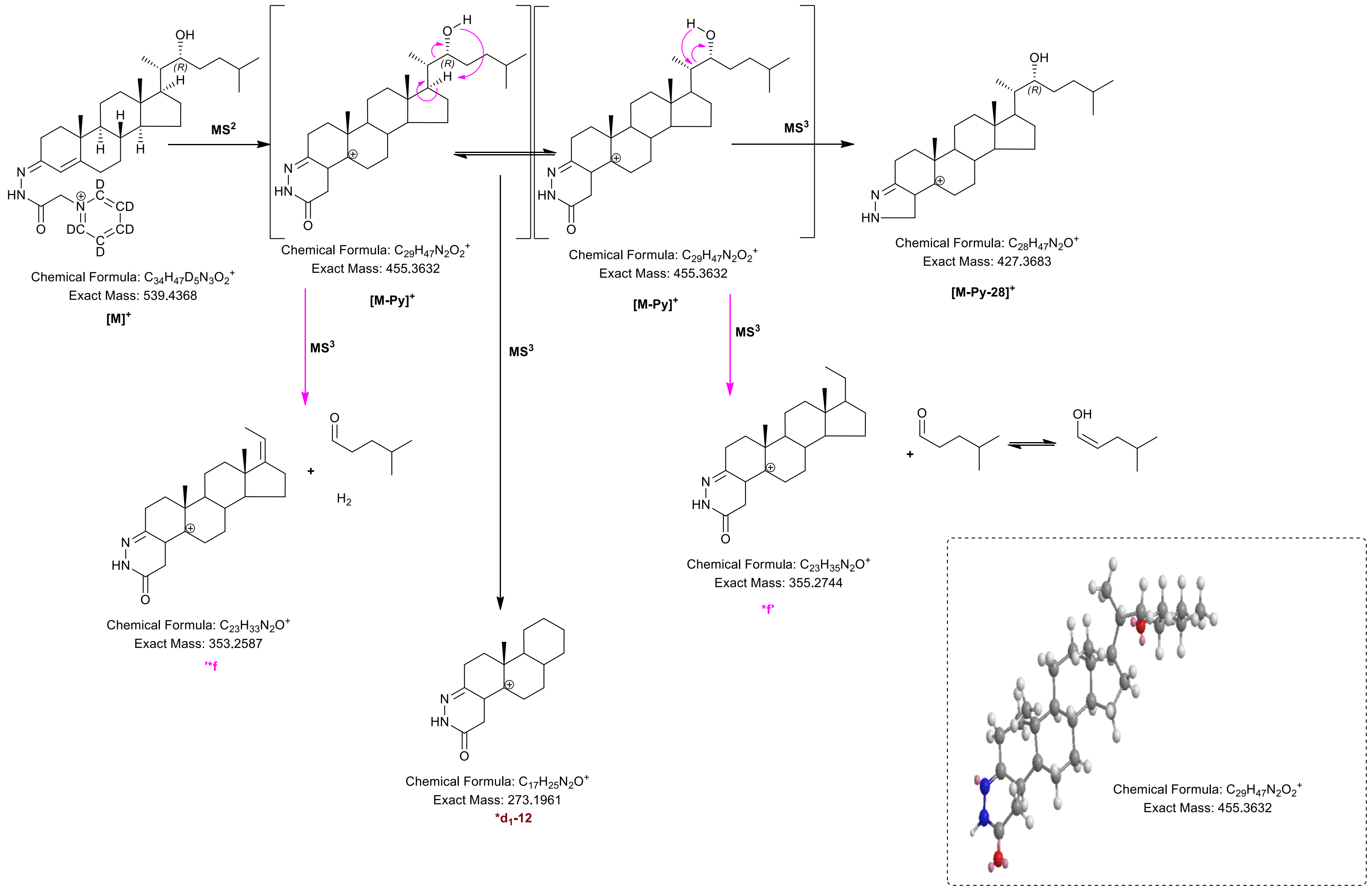

MS2
MS3
MS3
MS3
MS3
Chemical Formula: C34H47D5N3O2+
Exact Mass: 539.4368
[M]+
Chemical Formula: C29H47N2O2+
Exact Mass: 455.3632
[M-Py]+
Chemical Formula: C29H47N2O2+
Exact Mass: 455.3632
[M-Py]+
Chemical Formula: C28H47N2O+
Exact Mass: 427.3683
[M-Py-28]+
Chemical Formula: C23H33N2O+
Exact Mass: 353.2587
**f
Chemical Formula: C17H25N2O+
Exact Mass: 273.1961
*d1-12
Chemical Formula: C23H35N2O+
Exact Mass: 355.2744
*f
Chemical Formula: C29H47N2O2+
Exact Mass: 455.3632

**Figure S2B**

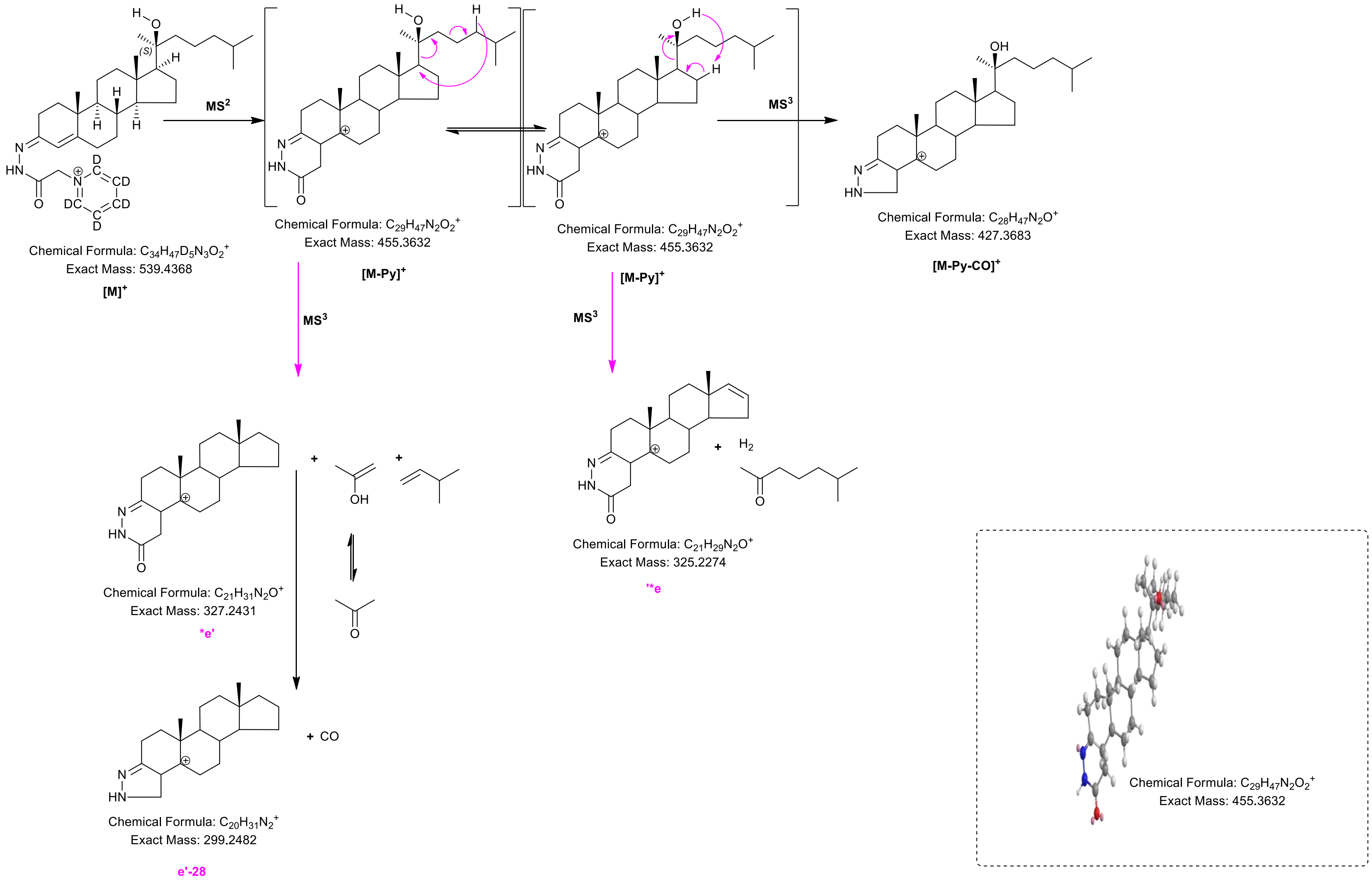

MS2
MS3
MS3
MS3
Chemical Formula: C34H47D5N3O2+
Exact Mass: 539.4368
[M]+
Chemical Formula: C29H47N2O2+
Exact Mass: 455.3632
[M-Py]+
Chemical Formula: C29H47N2O2+
Exact Mass: 455.3632
[M-Py]+
Chemical Formula: C28H47N2O+
Exact Mass: 427.3683
[M-Py-CO]+
Chemical Formula: C21H31N2O+
Exact Mass: 327.2431
*e'
Chemical Formula: C21H29N2O+
Exact Mass: 325.2274
'*e
Chemical Formula: C20H31N2+
Exact Mass: 299.2482
e'-28
+ CO
H2
Chemical Formula: C29H47N2O2+
Exact Mass: 455.3632

**Figure S2C**

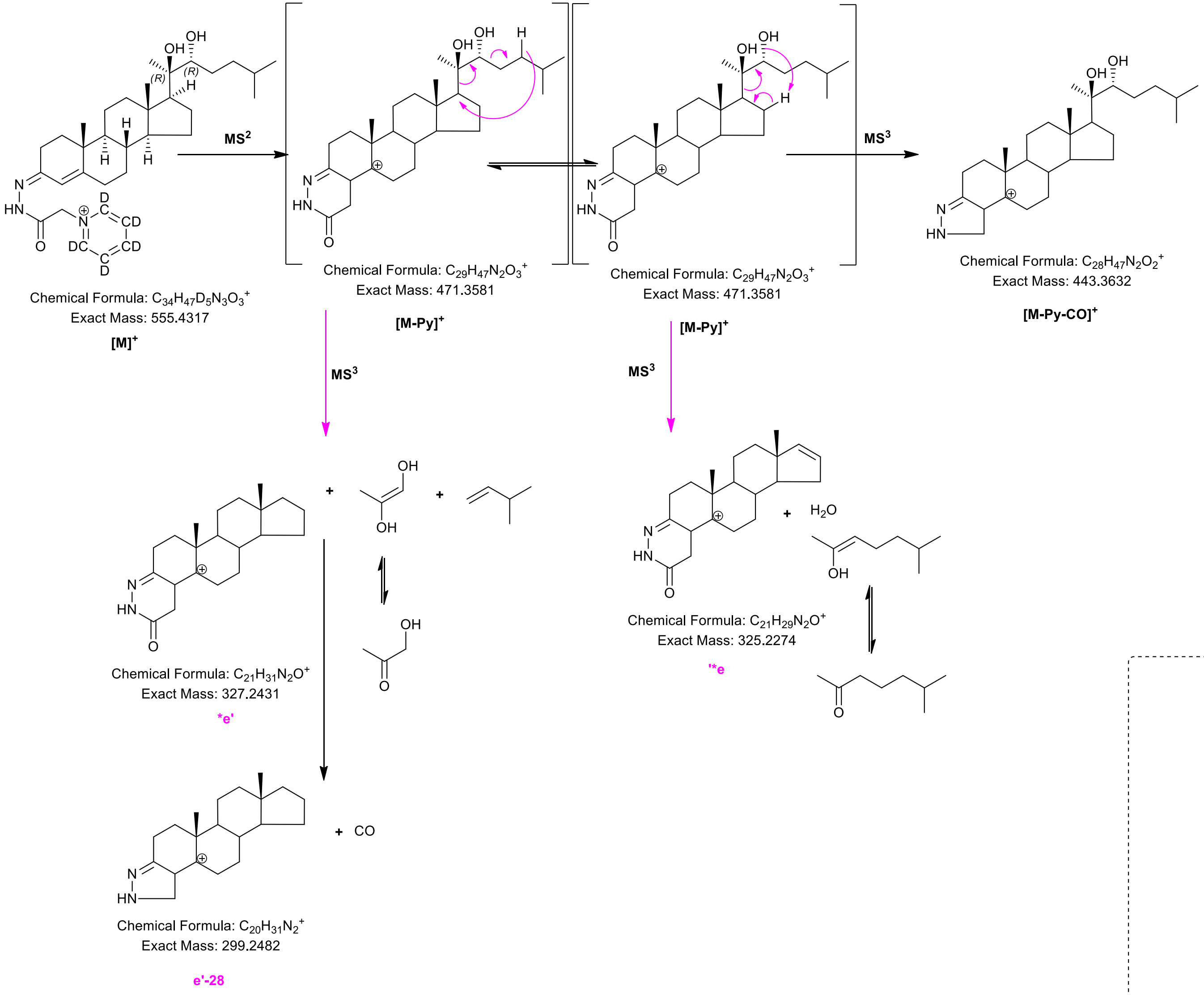

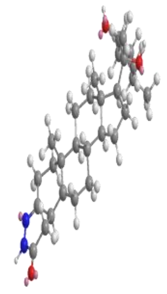

Chemical Formula: $C_{29}H_{47}N_2O_3^+$
Exact Mass: 471.3581

**Figure S2D**

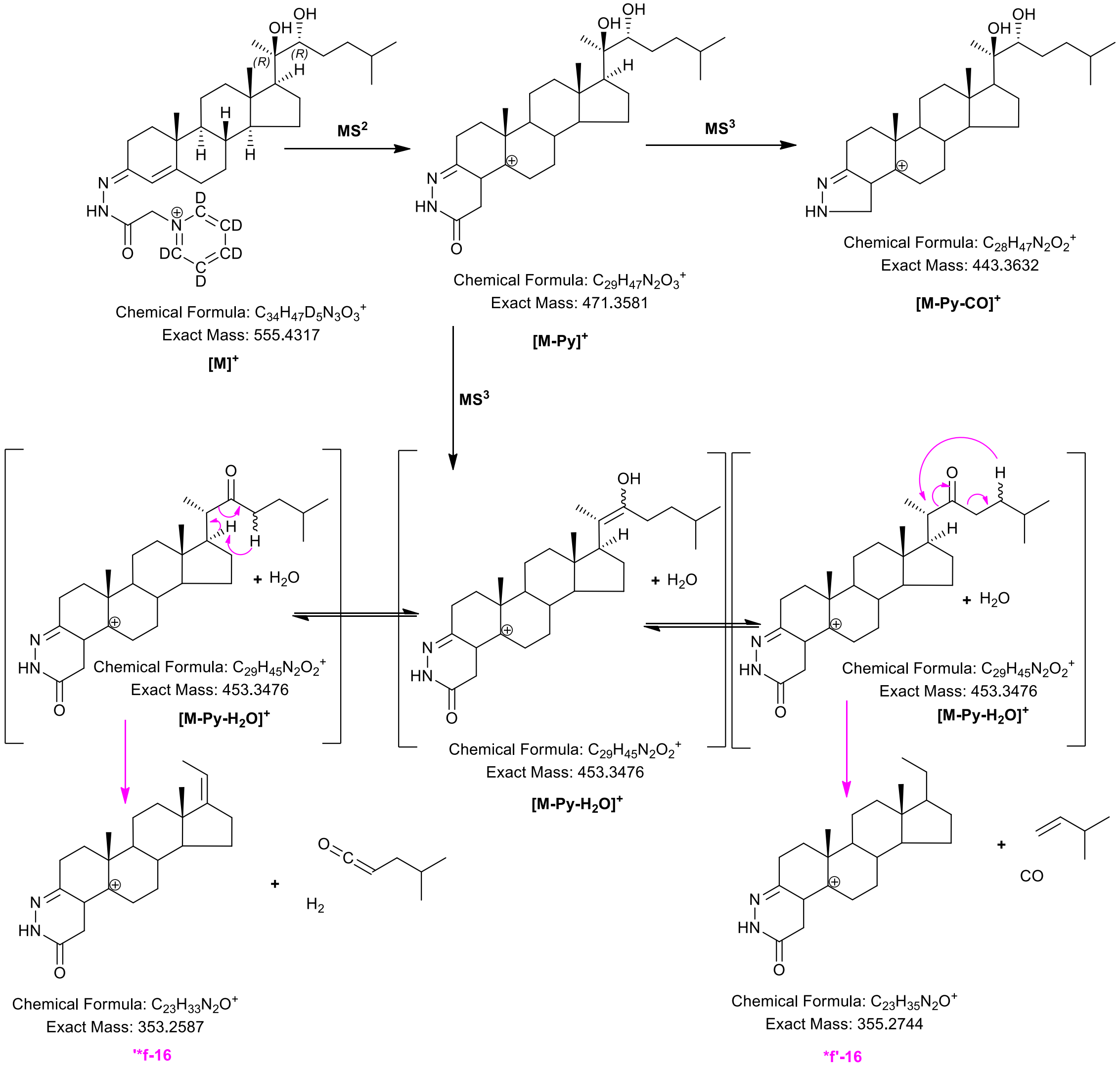
MS$^2$
MS$^3$
MS$^3$
Chemical Formula: $C_{34}H_{47}D_5N_3O_3^+$
Exact Mass: 555.4317
[M]$^+$
Chemical Formula: $C_{29}H_{47}N_2O_3^+$
Exact Mass: 471.3581
[M-Py]$^+$
Chemical Formula: $C_{28}H_{47}N_2O_2^+$
Exact Mass: 443.3632
[M-Py-CO]$^+$
+ $H_2O$
Chemical Formula: $C_{29}H_{45}N_2O_2^+$
Exact Mass: 453.3476
[M-Py-$H_2O$]$^+$
+ $H_2O$
Chemical Formula: $C_{29}H_{45}N_2O_2^+$
Exact Mass: 453.3476
[M-Py-$H_2O$]$^+$
+ $H_2O$
Chemical Formula: $C_{29}H_{45}N_2O_2^+$
Exact Mass: 453.3476
[M-Py-$H_2O$]$^+$
+ $H_2$
+ CO
Chemical Formula: $C_{23}H_{33}N_2O^+$
Exact Mass: 353.2587
'*f-16
Chemical Formula: $C_{23}H_{35}N_2O^+$
Exact Mass: 355.2744
*f'-16

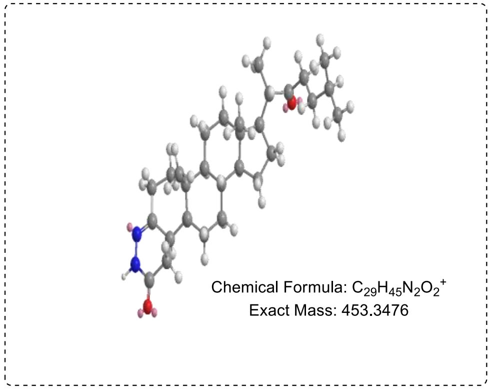
Chemical Formula: $C_{29}H_{45}N_2O_2^+$
Exact Mass: 453.3476

**Figure S2E**

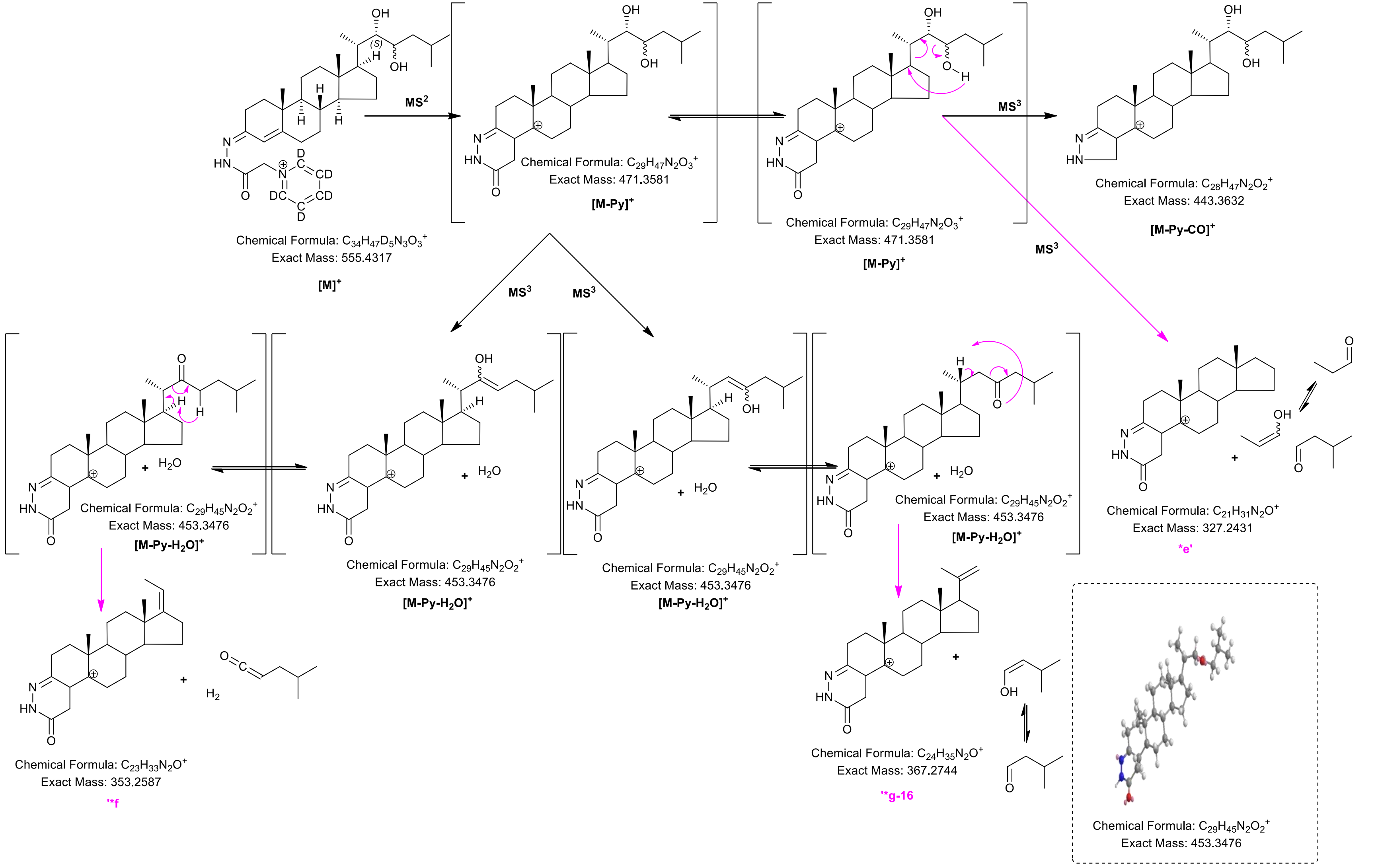

MS$^2$
MS$^3$
Chemical Formula: $C_{34}H_{47}D_5N_3O_3^+$
Exact Mass: 555.4317
[M]$^+$
Chemical Formula: $C_{29}H_{47}N_2O_3^+$
Exact Mass: 471.3581
[M-Py]$^+$
Chemical Formula: $C_{29}H_{47}N_2O_3^+$
Exact Mass: 471.3581
[M-Py]$^+$
Chemical Formula: $C_{28}H_{47}N_2O_2^+$
Exact Mass: 443.3632
[M-Py-CO]$^+$
Chemical Formula: $C_{29}H_{45}N_2O_2^+$
Exact Mass: 453.3476
[M-Py-$H_2O$]$^+$
Chemical Formula: $C_{29}H_{45}N_2O_2^+$
Exact Mass: 453.3476
[M-Py-$H_2O$]$^+$
Chemical Formula: $C_{29}H_{45}N_2O_2^+$
Exact Mass: 453.3476
[M-Py-$H_2O$]$^+$
Chemical Formula: $C_{29}H_{45}N_2O_2^+$
Exact Mass: 453.3476
[M-Py-$H_2O$]$^+$
Chemical Formula: $C_{21}H_{31}N_2O^+$
Exact Mass: 327.2431
*e'
Chemical Formula: $C_{23}H_{33}N_2O^+$
Exact Mass: 353.2587
**f
Chemical Formula: $C_{24}H_{35}N_2O^+$
Exact Mass: 367.2744
**g-16
Chemical Formula: $C_{29}H_{45}N_2O_2^+$
Exact Mass: 453.3476

**Figure S2F**

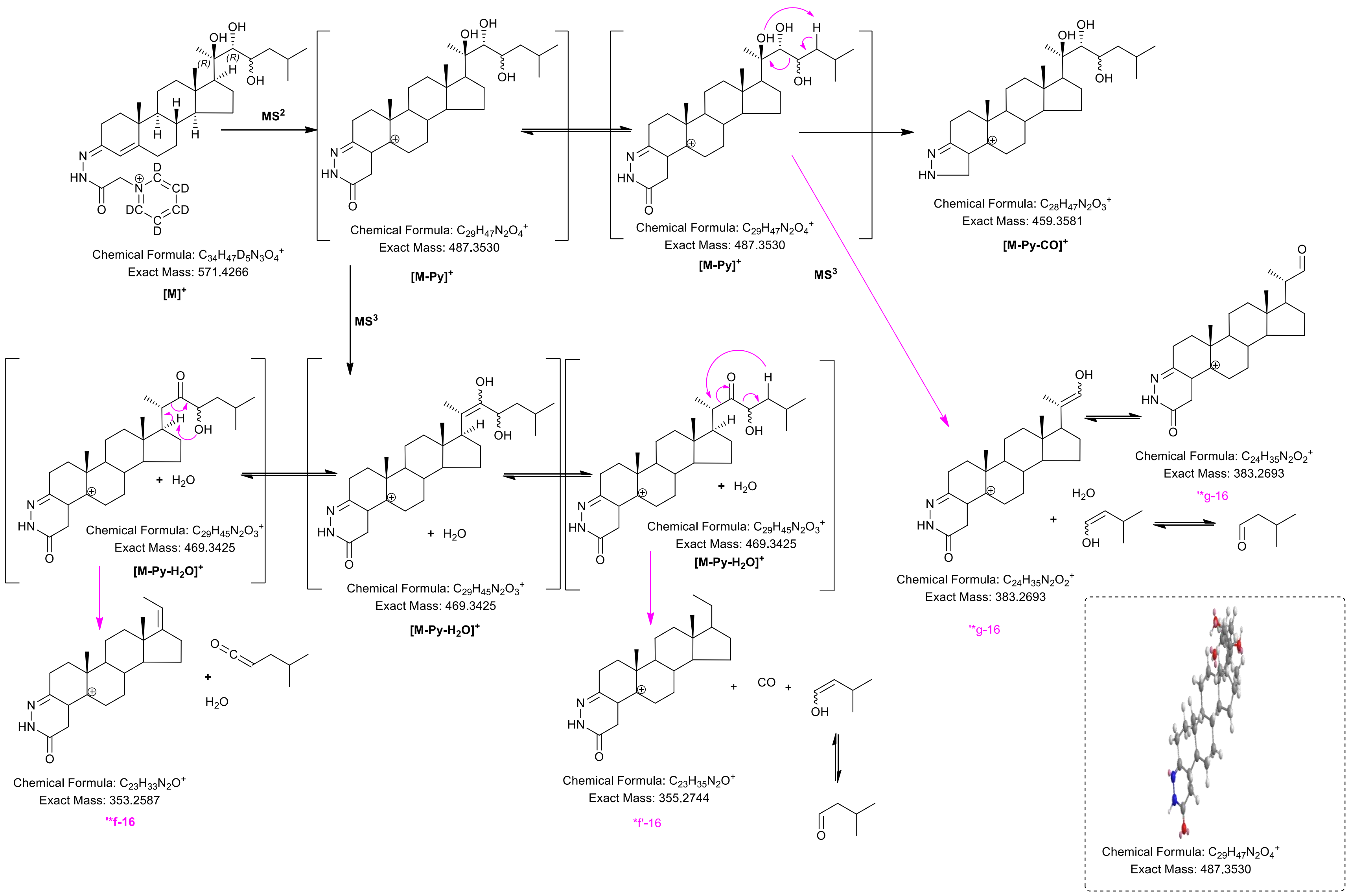
MS$^2$
MS$^3$
MS$^3$
Chemical Formula: $C_{34}H_{47}D_5N_3O_4^+$
Exact Mass: 571.4266
[M]$^+$
Chemical Formula: $C_{29}H_{47}N_2O_4^+$
Exact Mass: 487.3530
[M-Py]$^+$
Chemical Formula: $C_{29}H_{47}N_2O_4^+$
Exact Mass: 487.3530
[M-Py]$^+$
Chemical Formula: $C_{28}H_{47}N_2O_3^+$
Exact Mass: 459.3581
[M-Py-CO]$^+$
Chemical Formula: $C_{29}H_{45}N_2O_3^+$
Exact Mass: 469.3425
[M-Py-H$_2$O]$^+$
Chemical Formula: $C_{29}H_{45}N_2O_3^+$
Exact Mass: 469.3425
[M-Py-H$_2$O]$^+$
Chemical Formula: $C_{29}H_{45}N_2O_3^+$
Exact Mass: 469.3425
[M-Py-H$_2$O]$^+$
Chemical Formula: $C_{24}H_{35}N_2O_2^+$
Exact Mass: 383.2693
'*g-16
Chemical Formula: $C_{24}H_{35}N_2O_2^+$
Exact Mass: 383.2693
'*g-16
Chemical Formula: $C_{23}H_{33}N_2O^+$
Exact Mass: 353.2587
'*f-16
Chemical Formula: $C_{23}H_{35}N_2O^+$
Exact Mass: 355.2744
*f'-16
Chemical Formula: $C_{29}H_{47}N_2O_4^+$
Exact Mass: 487.3530

**Figure S2G**

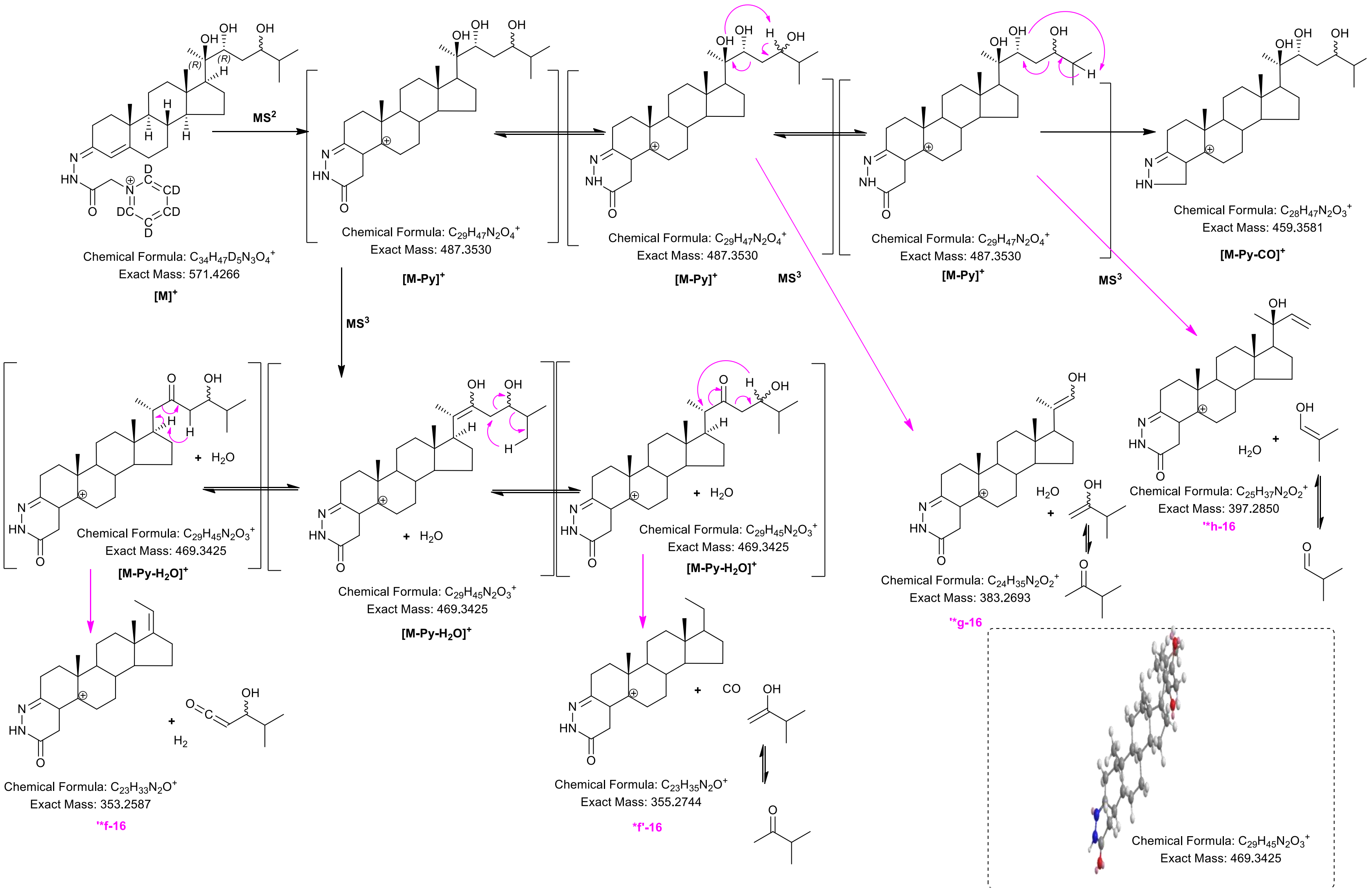
MS2
Chemical Formula: $C_{34}H_{47}D_5N_3O_4^+$
Exact Mass: 571.4266
[M]+
Chemical Formula: $C_{29}H_{47}N_2O_4^+$
Exact Mass: 487.3530
[M-Py]+
Chemical Formula: $C_{29}H_{47}N_2O_4^+$
Exact Mass: 487.3530
[M-Py]+
MS3
Chemical Formula: $C_{29}H_{47}N_2O_4^+$
Exact Mass: 487.3530
[M-Py]+
MS3
Chemical Formula: $C_{28}H_{47}N_2O_3^+$
Exact Mass: 459.3581
[M-Py-CO]+
MS3
Chemical Formula: $C_{29}H_{45}N_2O_3^+$
Exact Mass: 469.3425
$[M-Py-H_2O]^+$
Chemical Formula: $C_{29}H_{45}N_2O_3^+$
Exact Mass: 469.3425
$[M-Py-H_2O]^+$
Chemical Formula: $C_{29}H_{45}N_2O_3^+$
Exact Mass: 469.3425
$[M-Py-H_2O]^+$
Chemical Formula: $C_{24}H_{35}N_2O_2^+$
Exact Mass: 383.2693
'*g-16
Chemical Formula: $C_{25}H_{37}N_2O_2^+$
Exact Mass: 397.2850
'*h-16
Chemical Formula: $C_{23}H_{33}N_2O^+$
Exact Mass: 353.2587
'*f-16
Chemical Formula: $C_{23}H_{35}N_2O^+$
Exact Mass: 355.2744
*f'-16
Chemical Formula: $C_{29}H_{45}N_2O_3^+$
Exact Mass: 469.3425

**Figure S2H**

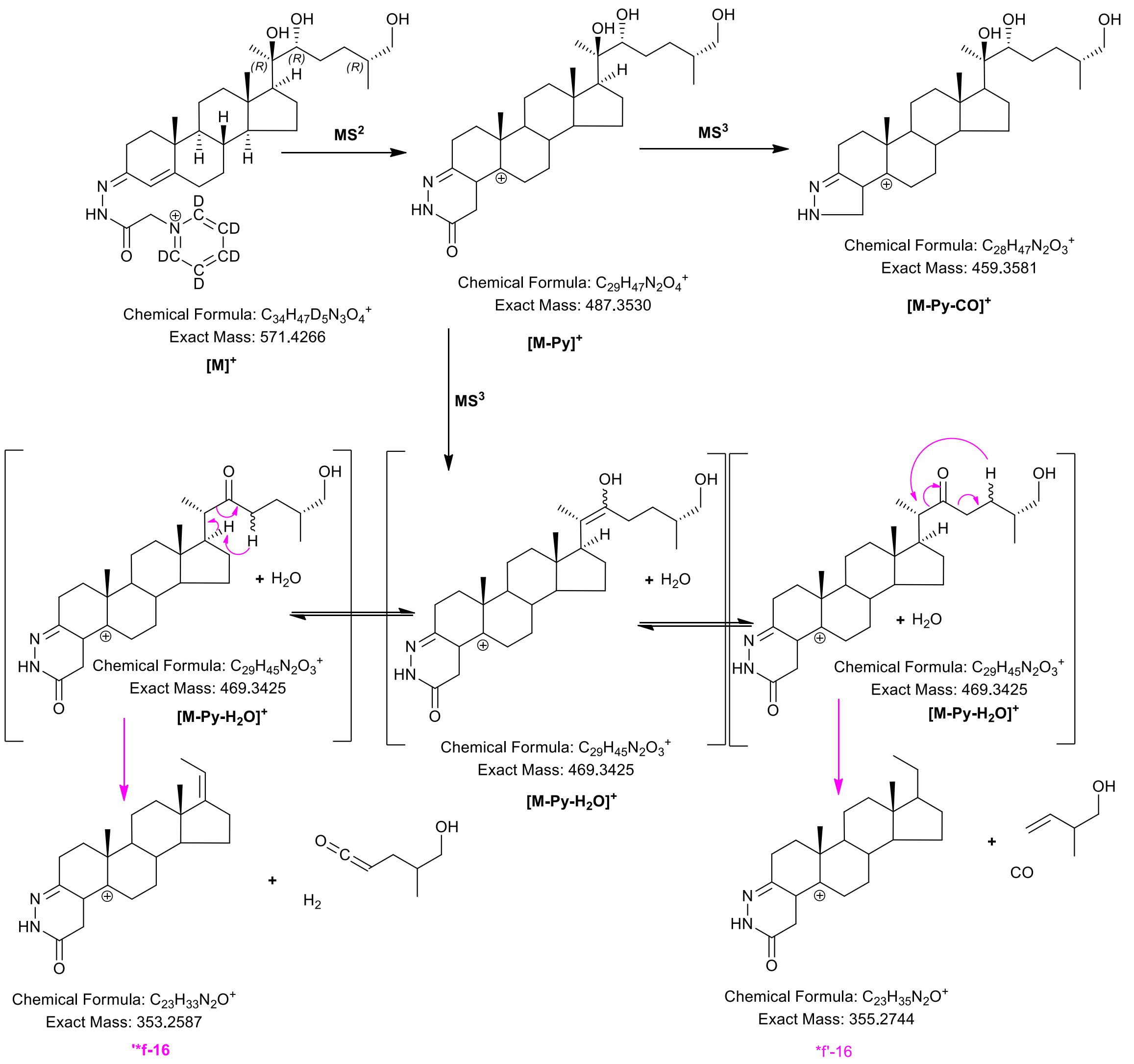
MS$^2$
MS$^3$
MS$^3$
Chemical Formula: $C_{34}H_{47}D_5N_3O_4^+$
Exact Mass: 571.4266
[M]$^+$
Chemical Formula: $C_{29}H_{47}N_2O_4^+$
Exact Mass: 487.3530
[M-Py]$^+$
Chemical Formula: $C_{28}H_{47}N_2O_3^+$
Exact Mass: 459.3581
[M-Py-CO]$^+$
+ $H_2O$
Chemical Formula: $C_{29}H_{45}N_2O_3^+$
Exact Mass: 469.3425
[M-Py-$H_2O$]$^+$
+ $H_2O$
Chemical Formula: $C_{29}H_{45}N_2O_3^+$
Exact Mass: 469.3425
[M-Py-$H_2O$]$^+$
+ $H_2O$
Chemical Formula: $C_{29}H_{45}N_2O_3^+$
Exact Mass: 469.3425
[M-Py-$H_2O$]$^+$
+ $H_2$
Chemical Formula: $C_{23}H_{33}N_2O^+$
Exact Mass: 353.2587
'*f-16
+ CO
Chemical Formula: $C_{23}H_{35}N_2O^+$
Exact Mass: 355.2744
*f'-16

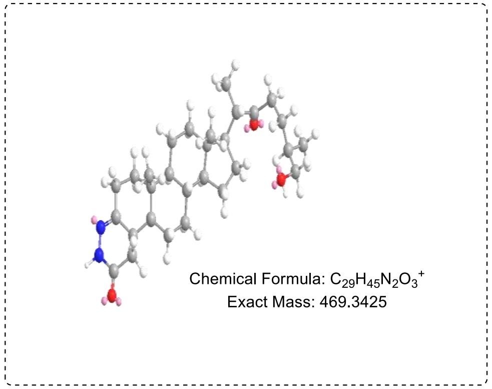
Chemical Formula: $C_{29}H_{45}N_2O_3^+$
Exact Mass: 469.3425

**Figure S2I**

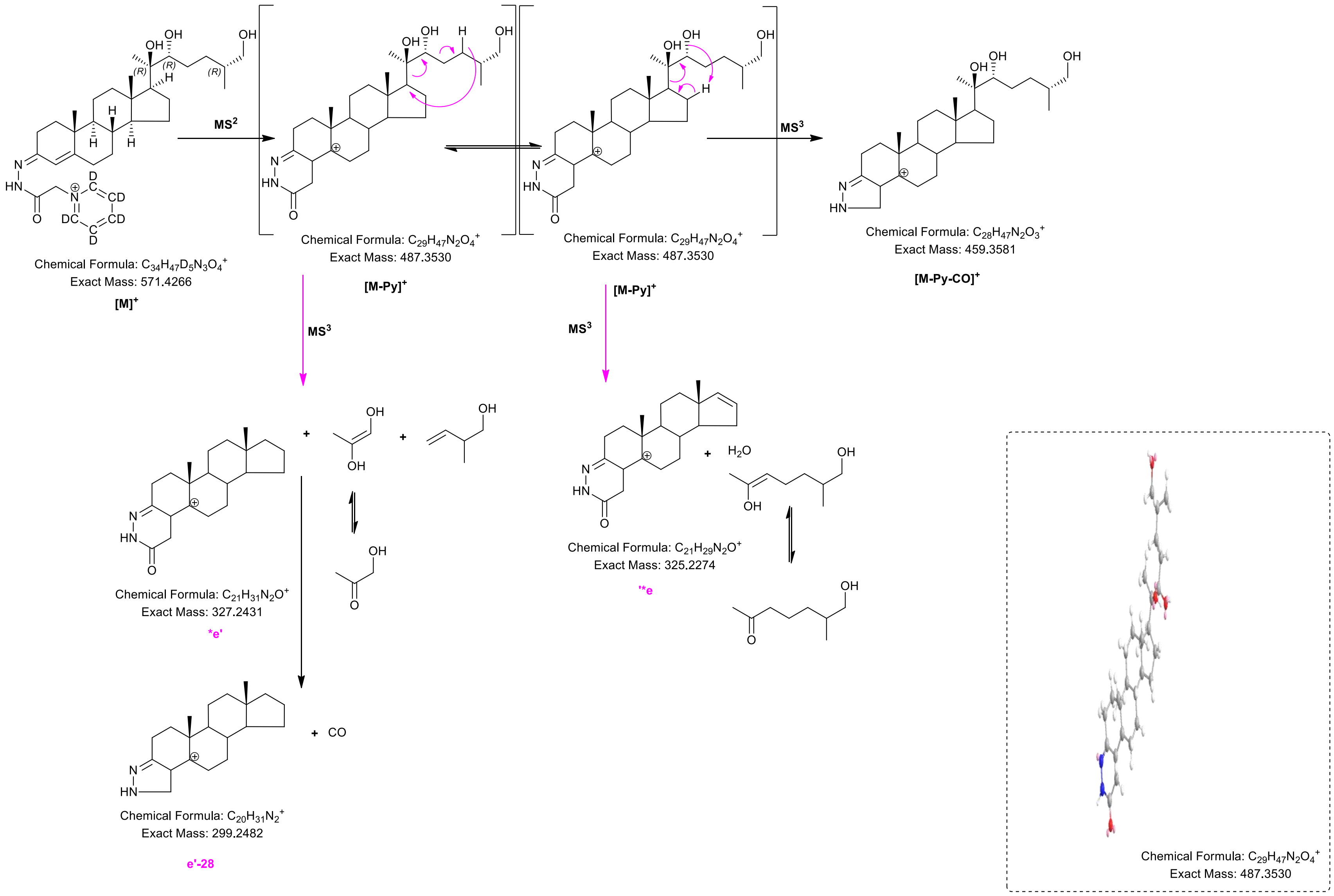

OH
OH
OH
(R)
(R)
(R)
H
H
H
H
N
HN
O
D
CD
CD
DC
D
MS2
Chemical Formula: $C_{34}H_{47}D_5N_3O_4^+$
Exact Mass: 571.4266
[M]+
Chemical Formula: $C_{29}H_{47}N_2O_4^+$
Exact Mass: 487.3530
[M-Py]+
Chemical Formula: $C_{29}H_{47}N_2O_4^+$
Exact Mass: 487.3530
[M-Py]+
MS3
Chemical Formula: $C_{28}H_{47}N_2O_3^+$
Exact Mass: 459.3581
[M-Py-CO]+
MS3
MS3
+
+
Chemical Formula: $C_{21}H_{31}N_2O^+$
Exact Mass: 327.2431
*e'
+ CO
Chemical Formula: $C_{20}H_{31}N_2^+$
Exact Mass: 299.2482
e'-28
+ $H_2O$
Chemical Formula: $C_{21}H_{29}N_2O^+$
Exact Mass: 325.2274
''*e
Chemical Formula: $C_{29}H_{47}N_2O_4^+$
Exact Mass: 487.3530

**Figure S2J**

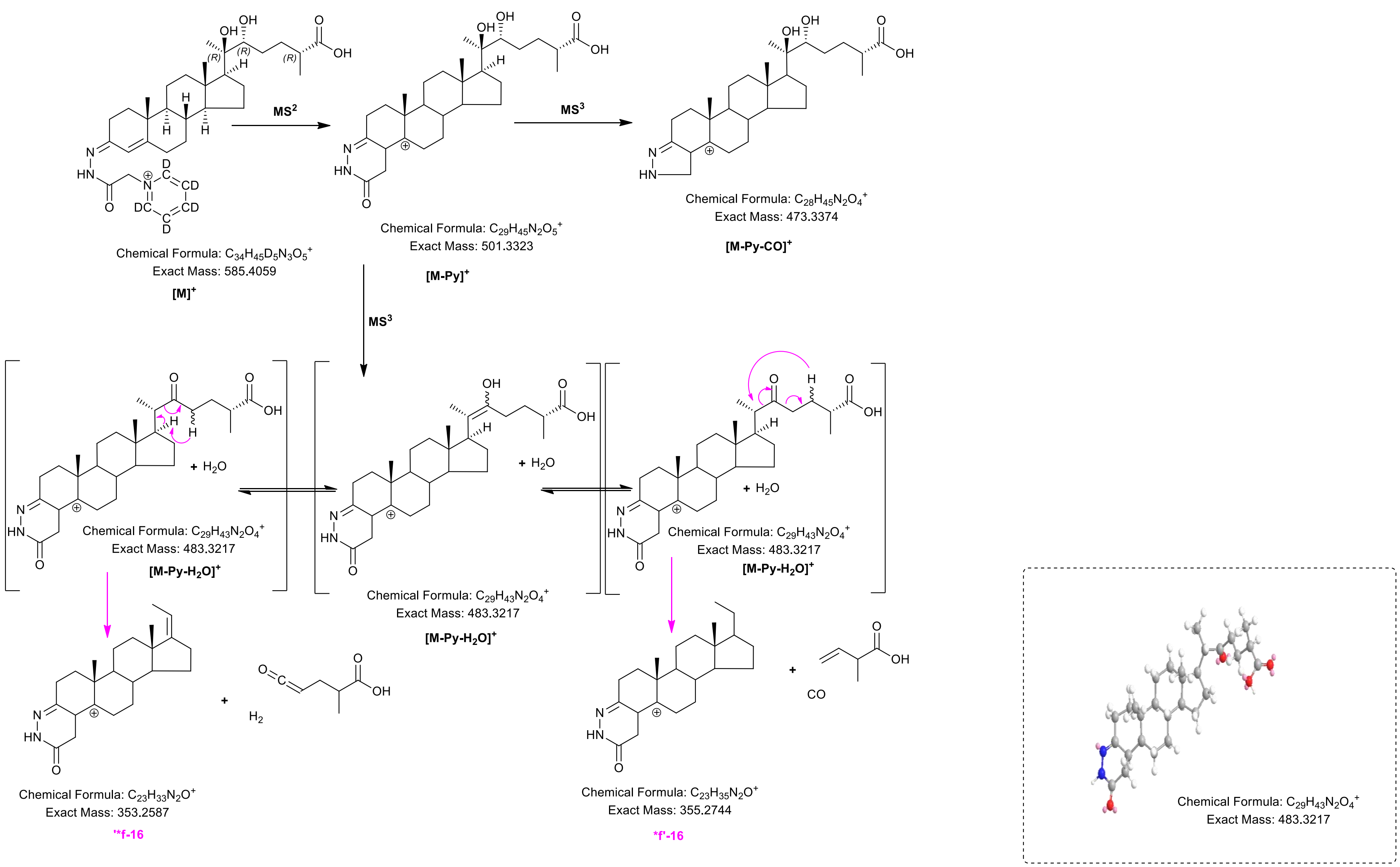
MS$^2$
MS$^3$
MS$^3$
Chemical Formula: $C_{34}H_{45}D_5N_3O_5^+$
Exact Mass: 585.4059
$[M]^+$
Chemical Formula: $C_{29}H_{45}N_2O_5^+$
Exact Mass: 501.3323
$[M\text{-}Py]^+$
Chemical Formula: $C_{28}H_{45}N_2O_4^+$
Exact Mass: 473.3374
$[M\text{-}Py\text{-}CO]^+$
+ $H_2O$
Chemical Formula: $C_{29}H_{43}N_2O_4^+$
Exact Mass: 483.3217
$[M\text{-}Py\text{-}H_2O]^+$
+ $H_2O$
Chemical Formula: $C_{29}H_{43}N_2O_4^+$
Exact Mass: 483.3217
$[M\text{-}Py\text{-}H_2O]^+$
+ $H_2O$
Chemical Formula: $C_{29}H_{43}N_2O_4^+$
Exact Mass: 483.3217
$[M\text{-}Py\text{-}H_2O]^+$
+ $H_2$
Chemical Formula: $C_{23}H_{33}N_2O^+$
Exact Mass: 353.2587
'*f-16
+ CO
Chemical Formula: $C_{23}H_{35}N_2O^+$
Exact Mass: 355.2744
*f'-16
Chemical Formula: $C_{29}H_{43}N_2O_4^+$
Exact Mass: 483.3217

**Figure S2K**

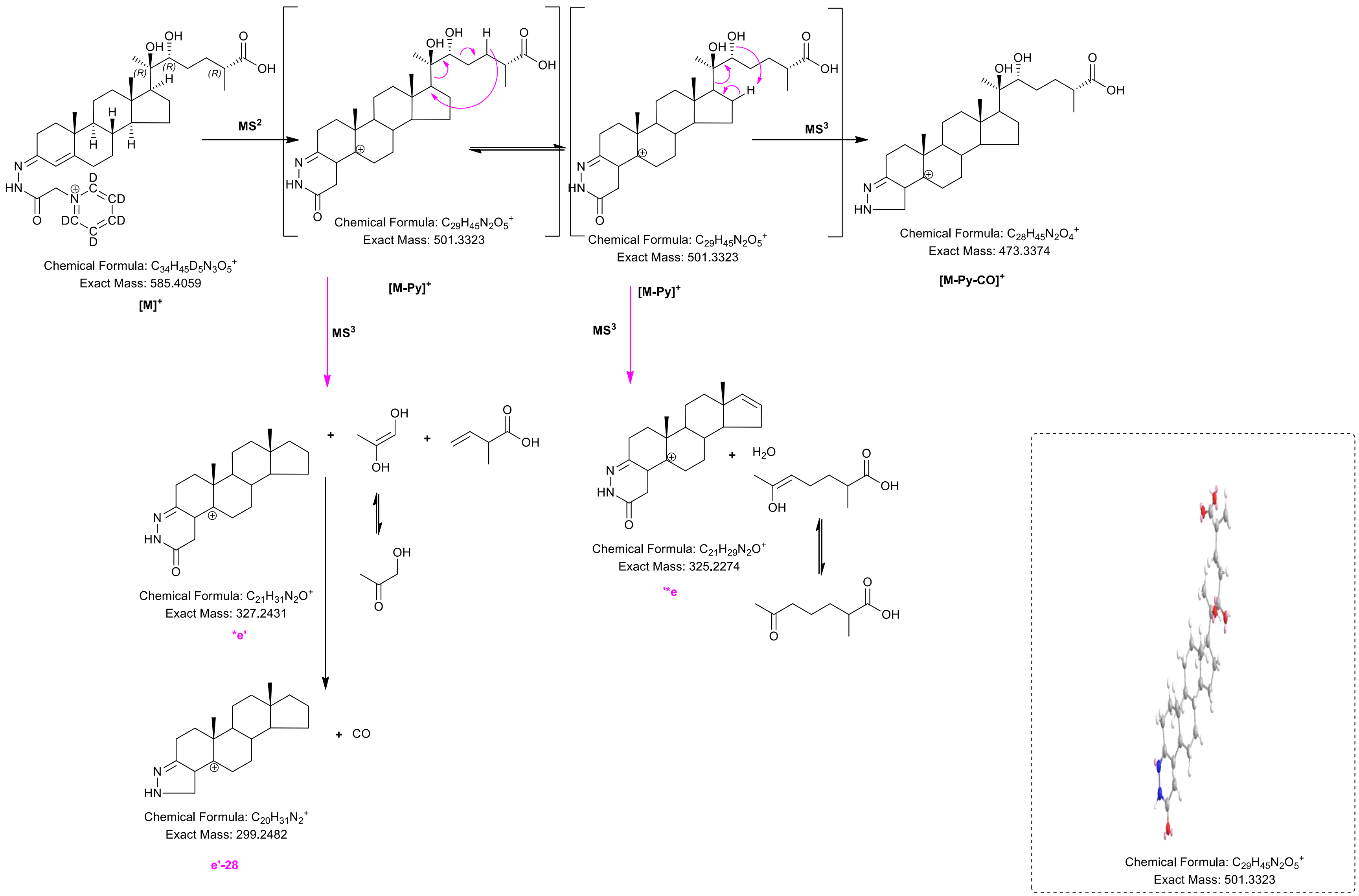

Chemical Formula: $C_{34}H_{45}D_5N_3O_5^+$
Exact Mass: 585.4059
[M]+
MS2
Chemical Formula: $C_{29}H_{45}N_2O_5^+$
Exact Mass: 501.3323
[M-Py]+
Chemical Formula: $C_{29}H_{45}N_2O_5^+$
Exact Mass: 501.3323
[M-Py]+
MS3
Chemical Formula: $C_{28}H_{45}N_2O_4^+$
Exact Mass: 473.3374
[M-Py-CO]+
MS3
MS3
Chemical Formula: $C_{21}H_{31}N_2O^+$
Exact Mass: 327.2431
*e'
+ CO
Chemical Formula: $C_{20}H_{31}N_2^+$
Exact Mass: 299.2482
e'-28
$H_2O$
Chemical Formula: $C_{21}H_{29}N_2O^+$
Exact Mass: 325.2274
'*e
Chemical Formula: $C_{29}H_{45}N_2O_5^+$
Exact Mass: 501.3323

**Figure S2L**

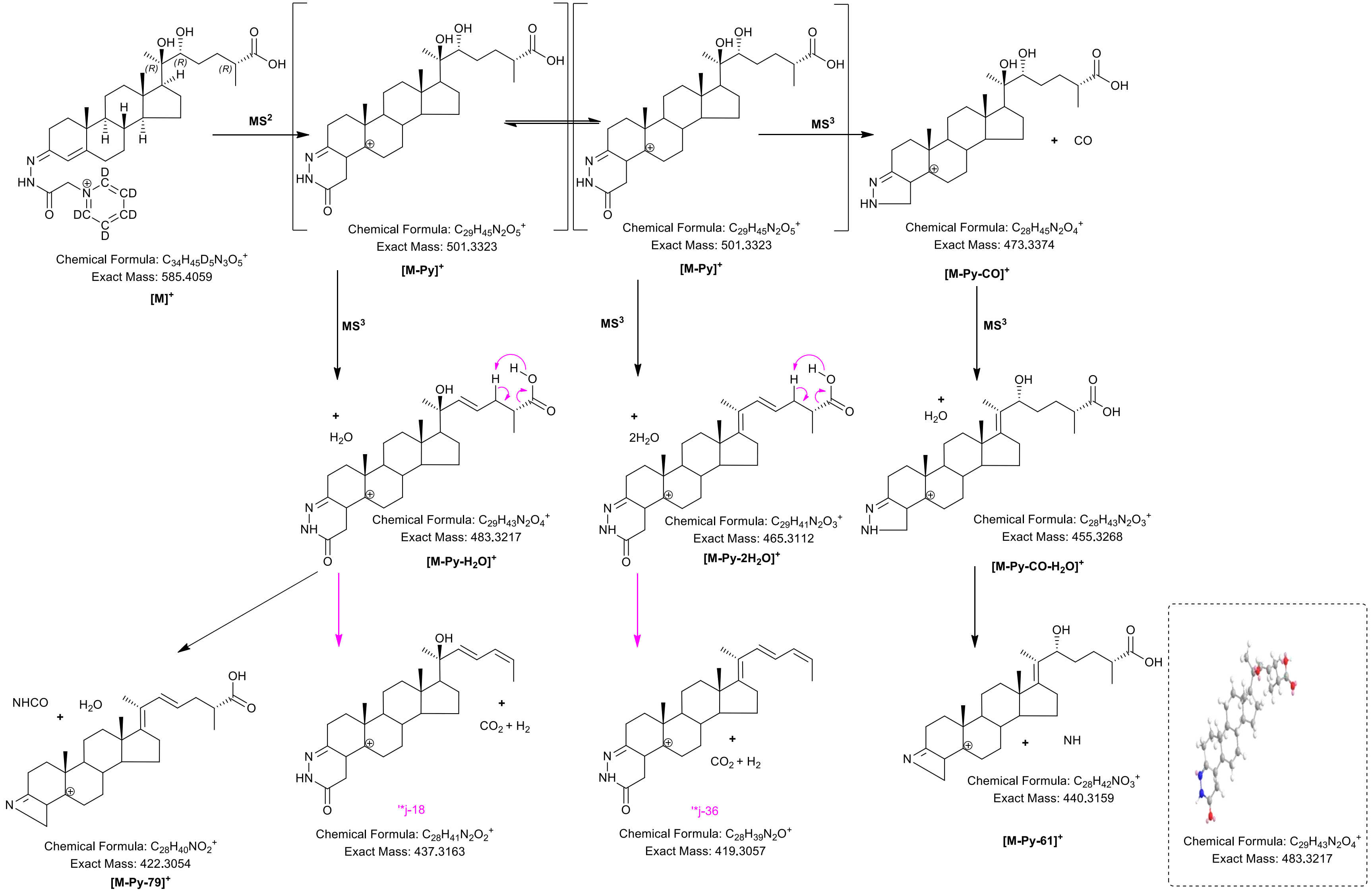

MS$^2$
MS$^3$
Chemical Formula: $C_{34}H_{45}D_5N_3O_5^+$
Exact Mass: 585.4059
[M]$^+$
Chemical Formula: $C_{29}H_{45}N_2O_5^+$
Exact Mass: 501.3323
[M-Py]$^+$
Chemical Formula: $C_{29}H_{45}N_2O_5^+$
Exact Mass: 501.3323
[M-Py]$^+$
+ CO
Chemical Formula: $C_{28}H_{45}N_2O_4^+$
Exact Mass: 473.3374
[M-Py-CO]$^+$
+ $H_2O$
Chemical Formula: $C_{29}H_{43}N_2O_4^+$
Exact Mass: 483.3217
[M-Py-$H_2O$]$^+$
+ $2H_2O$
Chemical Formula: $C_{29}H_{41}N_2O_3^+$
Exact Mass: 465.3112
[M-Py-2$H_2O$]$^+$
+ $H_2O$
Chemical Formula: $C_{28}H_{43}N_2O_3^+$
Exact Mass: 455.3268
[M-Py-CO-$H_2O$]$^+$
NHCO + $H_2O$
Chemical Formula: $C_{28}H_{40}NO_2^+$
Exact Mass: 422.3054
[M-Py-79]$^+$
+ $CO_2$ + $H_2$
'*j-18
Chemical Formula: $C_{28}H_{41}N_2O_2^+$
Exact Mass: 437.3163
+ $CO_2$ + $H_2$
'*j-36
Chemical Formula: $C_{28}H_{39}N_2O^+$
Exact Mass: 419.3057
+ NH
Chemical Formula: $C_{28}H_{42}NO_3^+$
Exact Mass: 440.3159
[M-Py-61]$^+$
Chemical Formula: $C_{29}H_{43}N_2O_4^+$
Exact Mass: 483.3217

**Figure S2M**

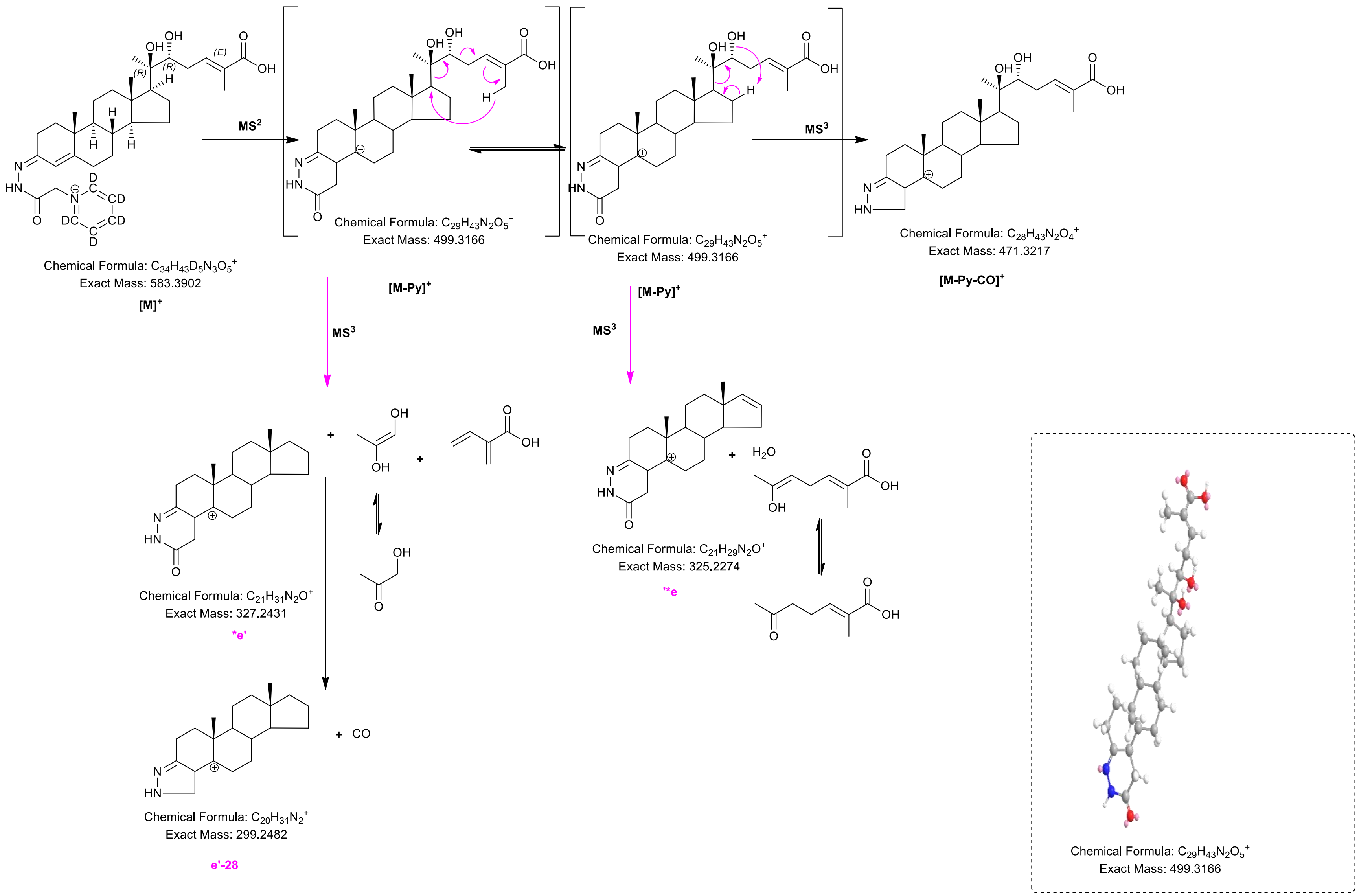

MS$^2$
MS$^3$
Chemical Formula: $C_{34}H_{43}D_5N_3O_5^+$
Exact Mass: 583.3902
[M]$^+$
Chemical Formula: $C_{29}H_{43}N_2O_5^+$
Exact Mass: 499.3166
[M-Py]$^+$
Chemical Formula: $C_{29}H_{43}N_2O_5^+$
Exact Mass: 499.3166
[M-Py]$^+$
Chemical Formula: $C_{28}H_{43}N_2O_4^+$
Exact Mass: 471.3217
[M-Py-CO]$^+$
Chemical Formula: $C_{21}H_{31}N_2O^+$
Exact Mass: 327.2431
*e'
Chemical Formula: $C_{21}H_{29}N_2O^+$
Exact Mass: 325.2274
'*e
+ $H_2O$
+ CO
Chemical Formula: $C_{20}H_{31}N_2^+$
Exact Mass: 299.2482
e'-28
Chemical Formula: $C_{29}H_{43}N_2O_5^+$
Exact Mass: 499.3166

**Figure S2N**

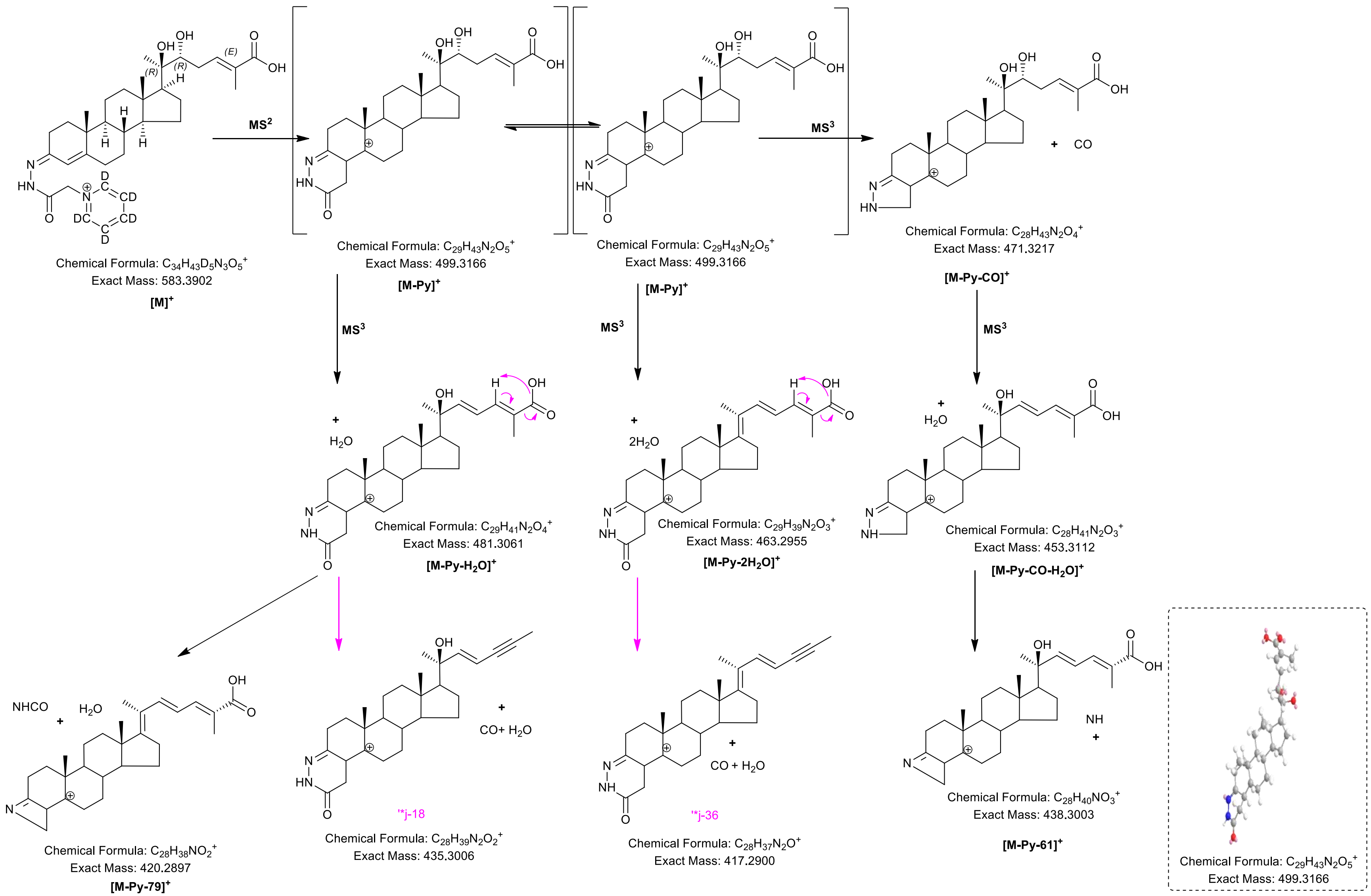

MS2
MS3
Chemical Formula: C34H43D5N3O5+
Exact Mass: 583.3902
[M]+
Chemical Formula: C29H43N2O5+
Exact Mass: 499.3166
[M-Py]+
Chemical Formula: C29H43N2O5+
Exact Mass: 499.3166
[M-Py]+
Chemical Formula: C28H43N2O4+
Exact Mass: 471.3217
[M-Py-CO]+
Chemical Formula: C29H41N2O4+
Exact Mass: 481.3061
[M-Py-H2O]+
Chemical Formula: C29H39N2O3+
Exact Mass: 463.2955
[M-Py-2H2O]+
Chemical Formula: C28H41N2O3+
Exact Mass: 453.3112
[M-Py-CO-H2O]+
Chemical Formula: C28H38NO2+
Exact Mass: 420.2897
[M-Py-79]+
'*j-18
Chemical Formula: C28H39N2O2+
Exact Mass: 435.3006
'*j-36
Chemical Formula: C28H37N2O+
Exact Mass: 417.2900
Chemical Formula: C28H40NO3+
Exact Mass: 438.3003
[M-Py-61]+
Chemical Formula: C29H43N2O5+
Exact Mass: 499.3166

**Figure S2O**

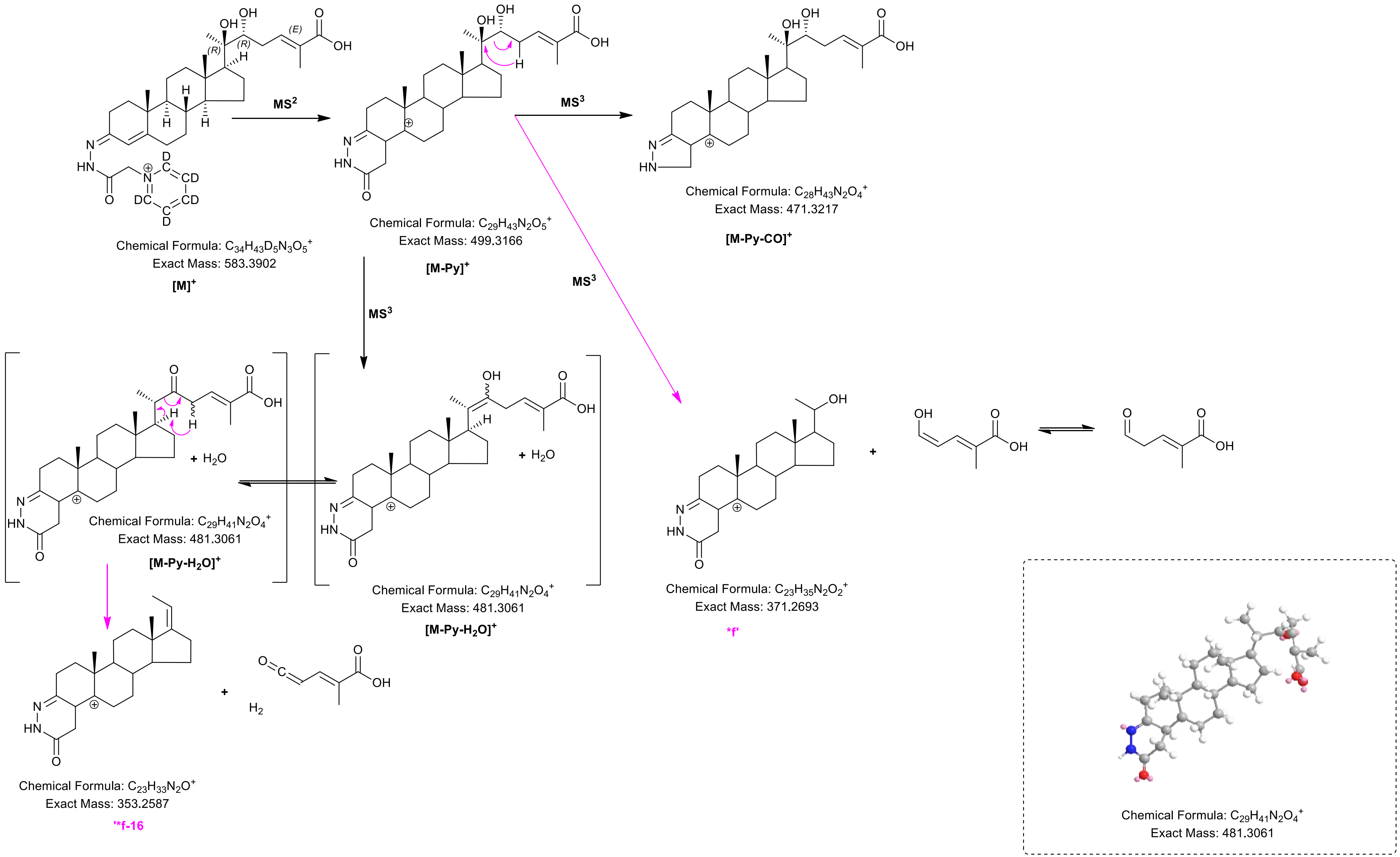
MS2
MS3
MS3
MS3
Chemical Formula: C34H43D5N3O5+
Exact Mass: 583.3902
[M]+
Chemical Formula: C29H43N2O5+
Exact Mass: 499.3166
[M-Py]+
Chemical Formula: C28H43N2O4+
Exact Mass: 471.3217
[M-Py-CO]+
+ H2O
Chemical Formula: C29H41N2O4+
Exact Mass: 481.3061
[M-Py-H2O]+
+ H2O
Chemical Formula: C29H41N2O4+
Exact Mass: 481.3061
[M-Py-H2O]+
Chemical Formula: C23H35N2O2+
Exact Mass: 371.2693
*f'
Chemical Formula: C23H33N2O+
Exact Mass: 353.2587
'*f-16
Chemical Formula: C29H41N2O4+
Exact Mass: 481.3061

S3A

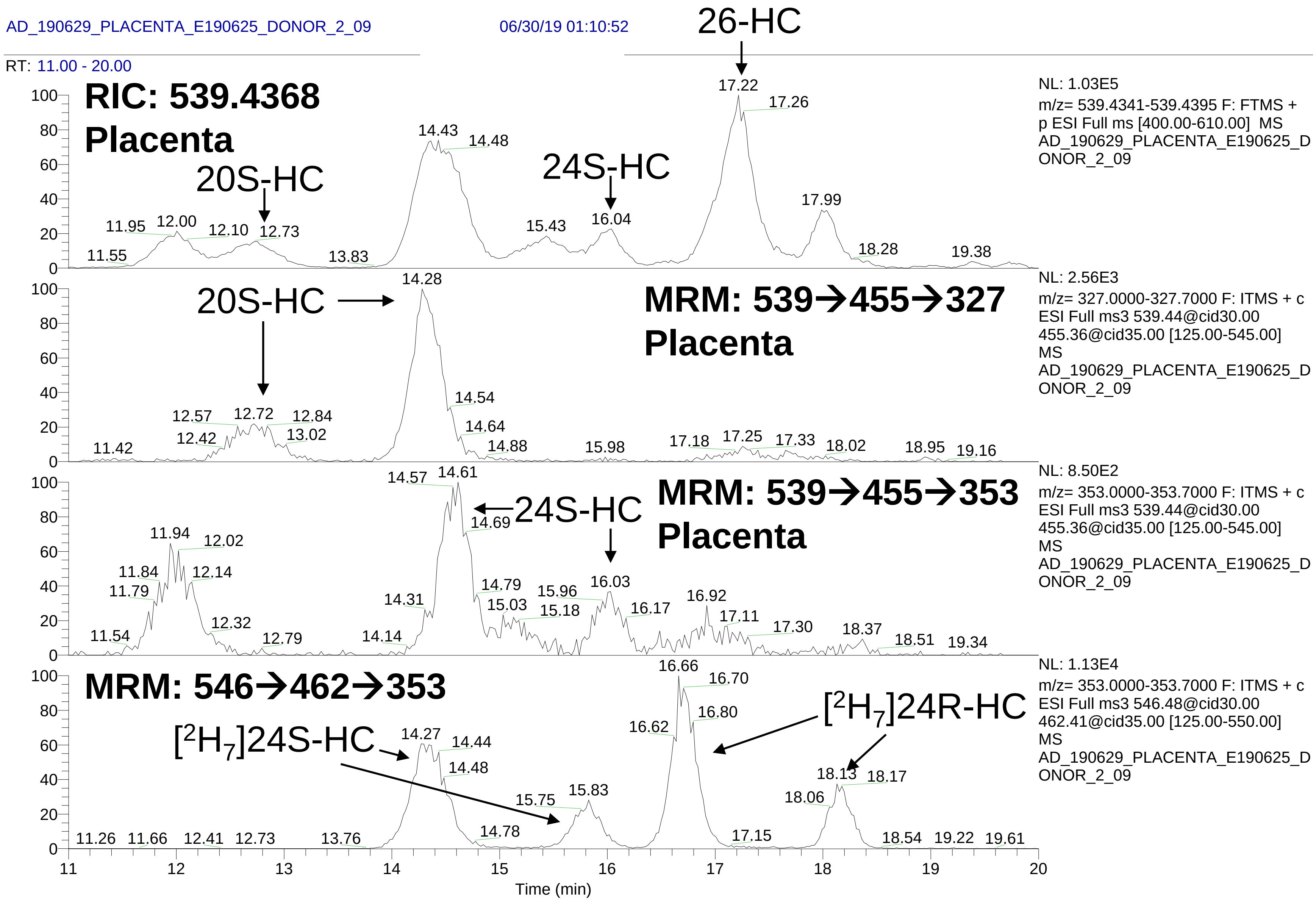
AD_190629_PLACENTA_E190625_DONOR_2_09
06/30/19 01:10:52
RT: 11.00 - 20.00
RIC: 539.4368
Placenta
20S-HC
24S-HC
26-HC
NL: 1.03E5
m/z= 539.4341-539.4395 F: FTMS + p ESI Full ms [400.00-610.00] MS AD_190629_PLACENTA_E190625_DONOR_2_09
MRM: 539→455→327
Placenta
NL: 2.56E3
m/z= 327.0000-327.7000 F: ITMS + c ESI Full ms3 539.44@cid30.00 455.36@cid35.00 [125.00-545.00] MS AD_190629_PLACENTA_E190625_DONOR_2_09
MRM: 539→455→353
Placenta
NL: 8.50E2
m/z= 353.0000-353.7000 F: ITMS + c ESI Full ms3 539.44@cid30.00 455.36@cid35.00 [125.00-545.00] MS AD_190629_PLACENTA_E190625_DONOR_2_09
MRM: 546→462→353
[2H7]24S-HC
[2H7]24R-HC
NL: 1.13E4
m/z= 353.0000-353.7000 F: ITMS + c ESI Full ms3 546.48@cid30.00 462.41@cid35.00 [125.00-550.00] MS AD_190629_PLACENTA_E190625_DONOR_2_09
Time (min)

# S3B

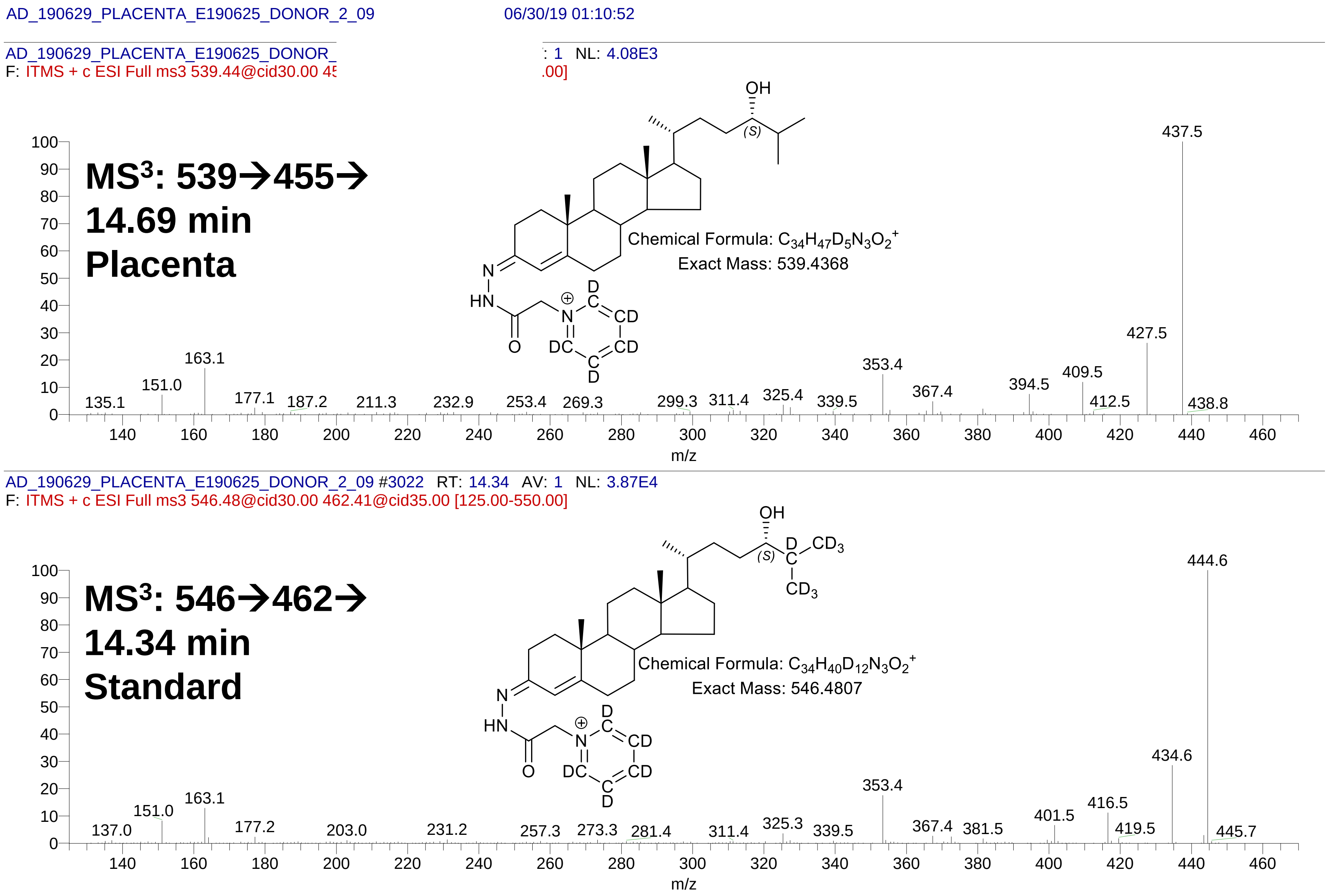

AD_190629_PLACENTA_E190625_DONOR_2_09
06/30/19 01:10:52
AD_190629_PLACENTA_E190625_DONOR_
: 1 NL: 4.08E3
F: ITMS + c ESI Full ms3 539.44@cid30.00 45
.00]
MS3: 539→455→
14.69 min
Placenta
OH
(S)
N
HN
O
N
D
C
CD
CD
DC
C
D
Chemical Formula: C34H47D5N3O2+
Exact Mass: 539.4368
437.5
427.5
409.5
412.5
438.8
394.5
367.4
353.4
339.5
325.4
311.4
299.3
269.3
253.4
232.9
211.3
187.2
177.1
163.1
151.0
135.1
m/z
AD_190629_PLACENTA_E190625_DONOR_2_09 #3022 RT: 14.34 AV: 1 NL: 3.87E4
F: ITMS + c ESI Full ms3 546.48@cid30.00 462.41@cid35.00 [125.00-550.00]
MS3: 546→462→
14.34 min
Standard
OH
(S)
D
CD3
C
CD3
N
HN
O
N
D
C
CD
CD
DC
C
D
Chemical Formula: C34H40D12N3O2+
Exact Mass: 546.4807
444.6
434.6
416.5
419.5
401.5
445.7
381.5
367.4
353.4
339.5
325.3
311.4
281.4
273.3
257.3
231.2
203.0
177.2
163.1
151.0
137.0
m/z

# S3C

AD_190629_PLACENTA_E190625_DONOR_2_09 06/30/19 01:10:52

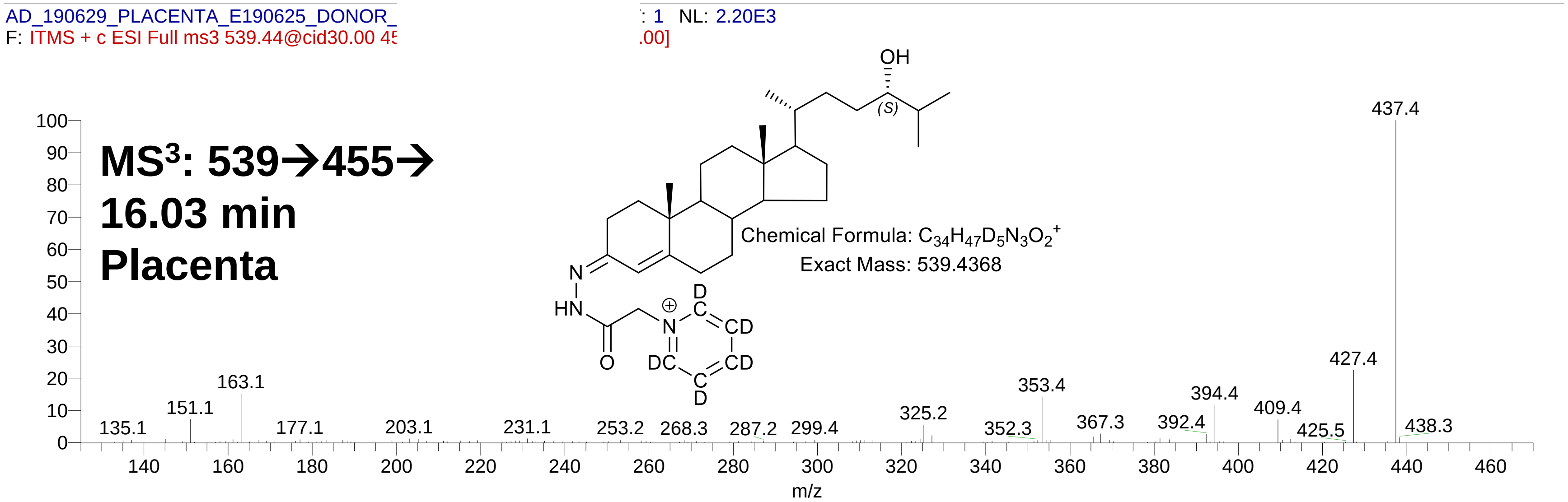

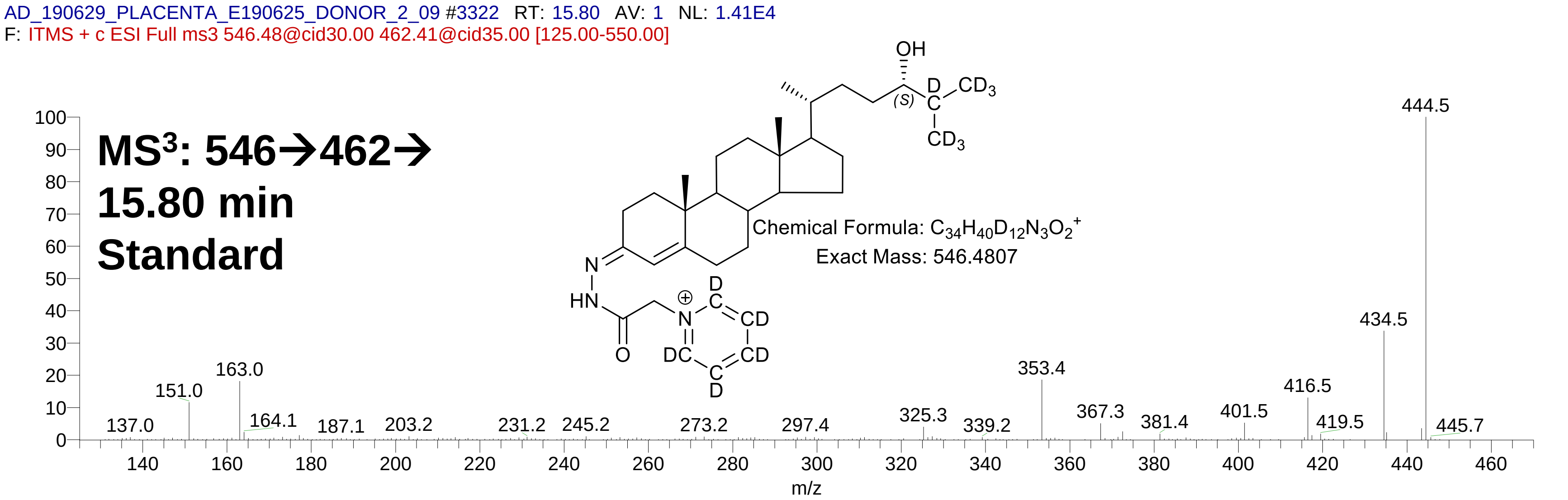

**S3D**

AD_190629_PLACENTA_E190625_DONOR_2_09 06/30/19 01:10:52

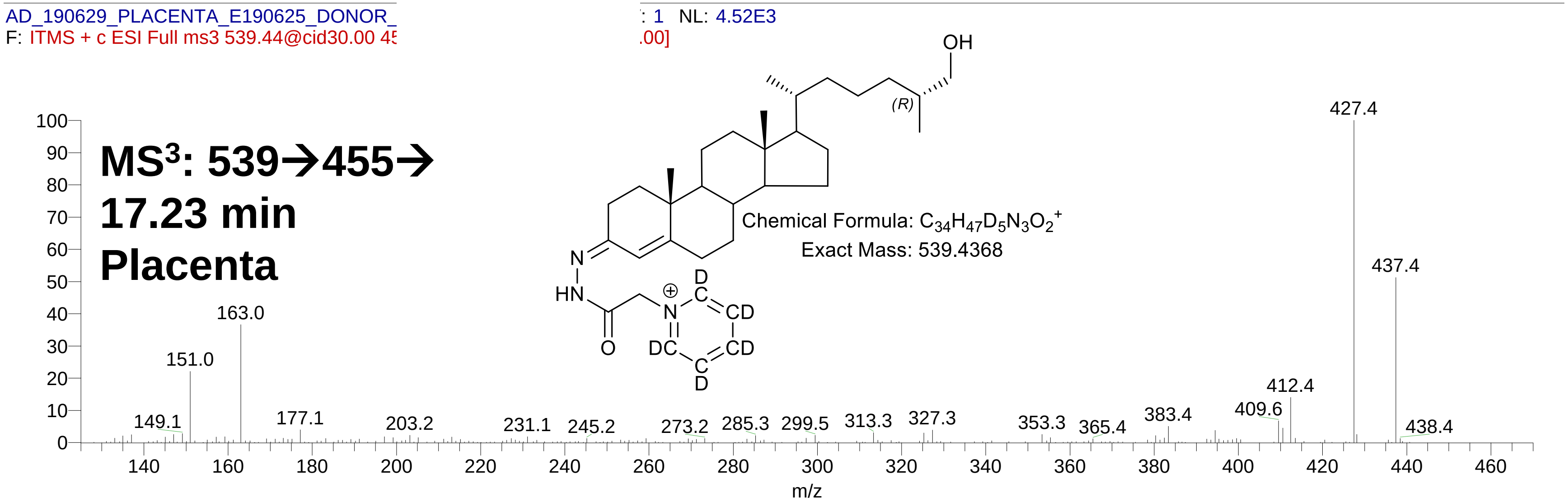

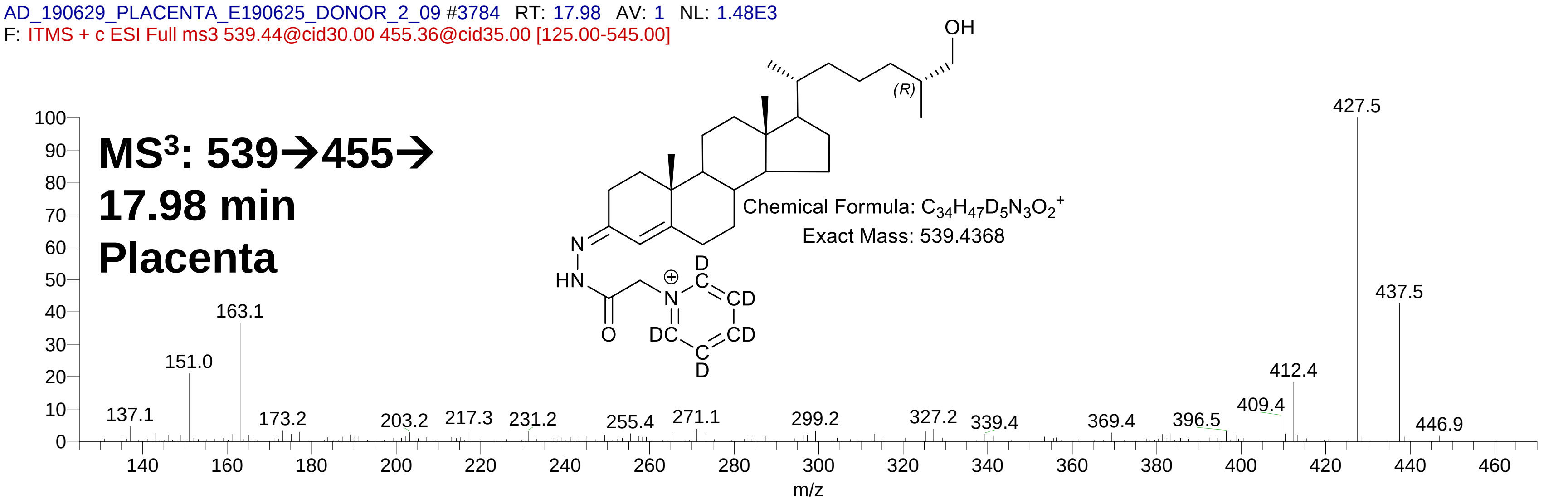

**S3E**

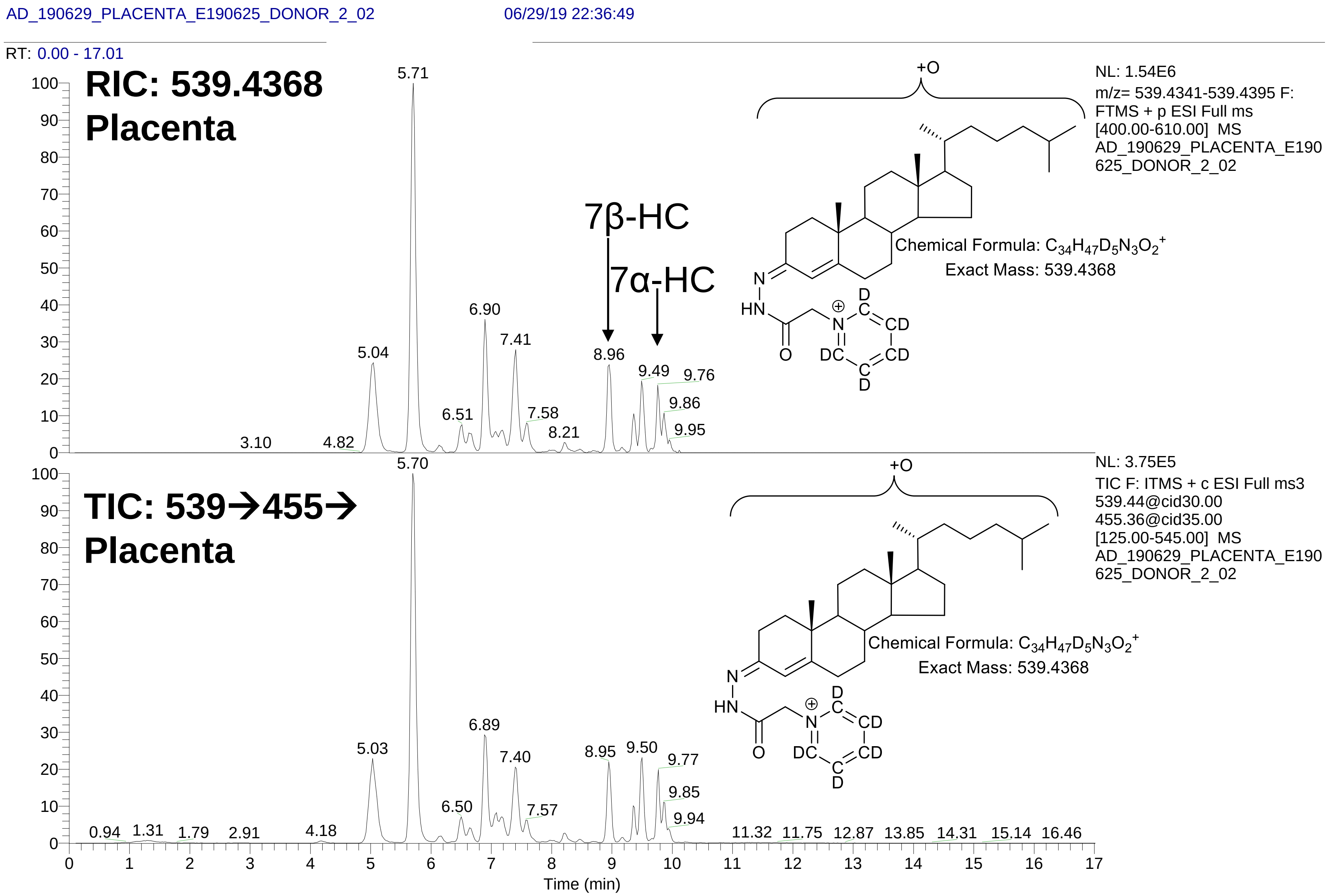
AD_190629_PLACENTA_E190625_DONOR_2_02
06/29/19 22:36:49
RT: 0.00 - 17.01
RIC: 539.4368
Placenta
NL: 1.54E6
m/z= 539.4341-539.4395 F: FTMS + p ESI Full ms [400.00-610.00] MS AD_190629_PLACENTA_E190625_DONOR_2_02
+O
7β-HC
7α-HC
Chemical Formula: $C_{34}H_{47}D_5N_3O_2^+$
Exact Mass: 539.4368
5.71
6.90
7.41
5.04
8.96
9.49
9.76
9.86
9.95
6.51
7.58
8.21
3.10
4.82
TIC: 539→455→
Placenta
NL: 3.75E5
TIC F: ITMS + c ESI Full ms3 539.44@cid30.00 455.36@cid35.00 [125.00-545.00] MS AD_190629_PLACENTA_E190625_DONOR_2_02
+O
Chemical Formula: $C_{34}H_{47}D_5N_3O_2^+$
Exact Mass: 539.4368
5.70
6.89
5.03
7.40
8.95
9.50
9.77
9.85
9.94
6.50
7.57
0.94
1.31
1.79
2.91
4.18
11.32
11.75
12.87
13.85
14.31
15.14
16.46
Time (min)

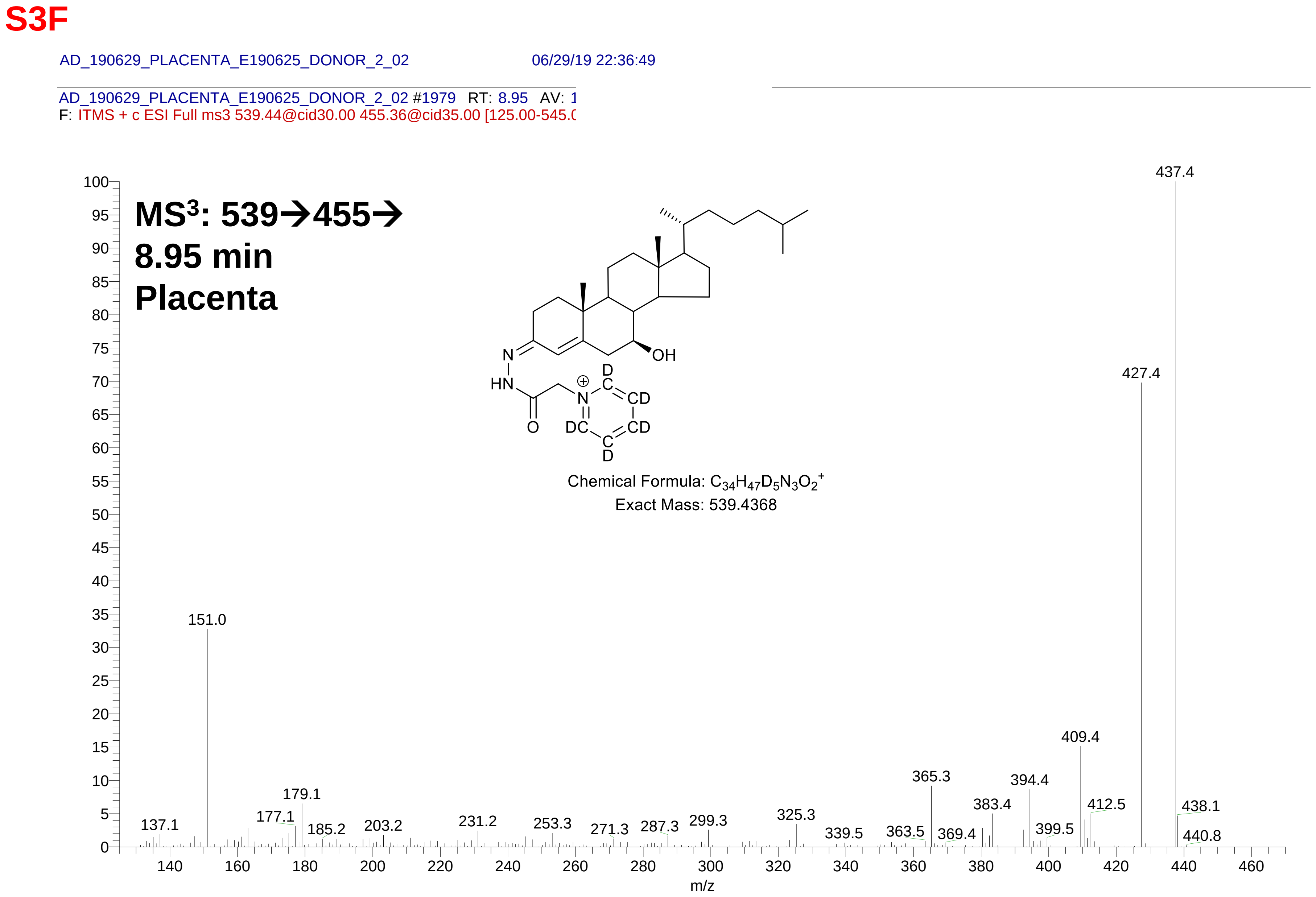
S3F
AD_190629_PLACENTA_E190625_DONOR_2_02
06/29/19 22:36:49
AD_190629_PLACENTA_E190625_DONOR_2_02 #1979 RT: 8.95 AV: 1
F: ITMS + c ESI Full ms3 539.44@cid30.00 455.36@cid35.00 [125.00-545.0
MS3: 539→455→
8.95 min
Placenta
Chemical Formula: C34H47D5N3O2+
Exact Mass: 539.4368
437.4
427.4
409.4
394.4
383.4
365.3
412.5
438.1
440.8
399.5
369.4
363.5
339.5
325.3
299.3
287.3
271.3
253.3
231.2
203.2
185.2
179.1
177.1
151.0
137.1
m/z

# S3G

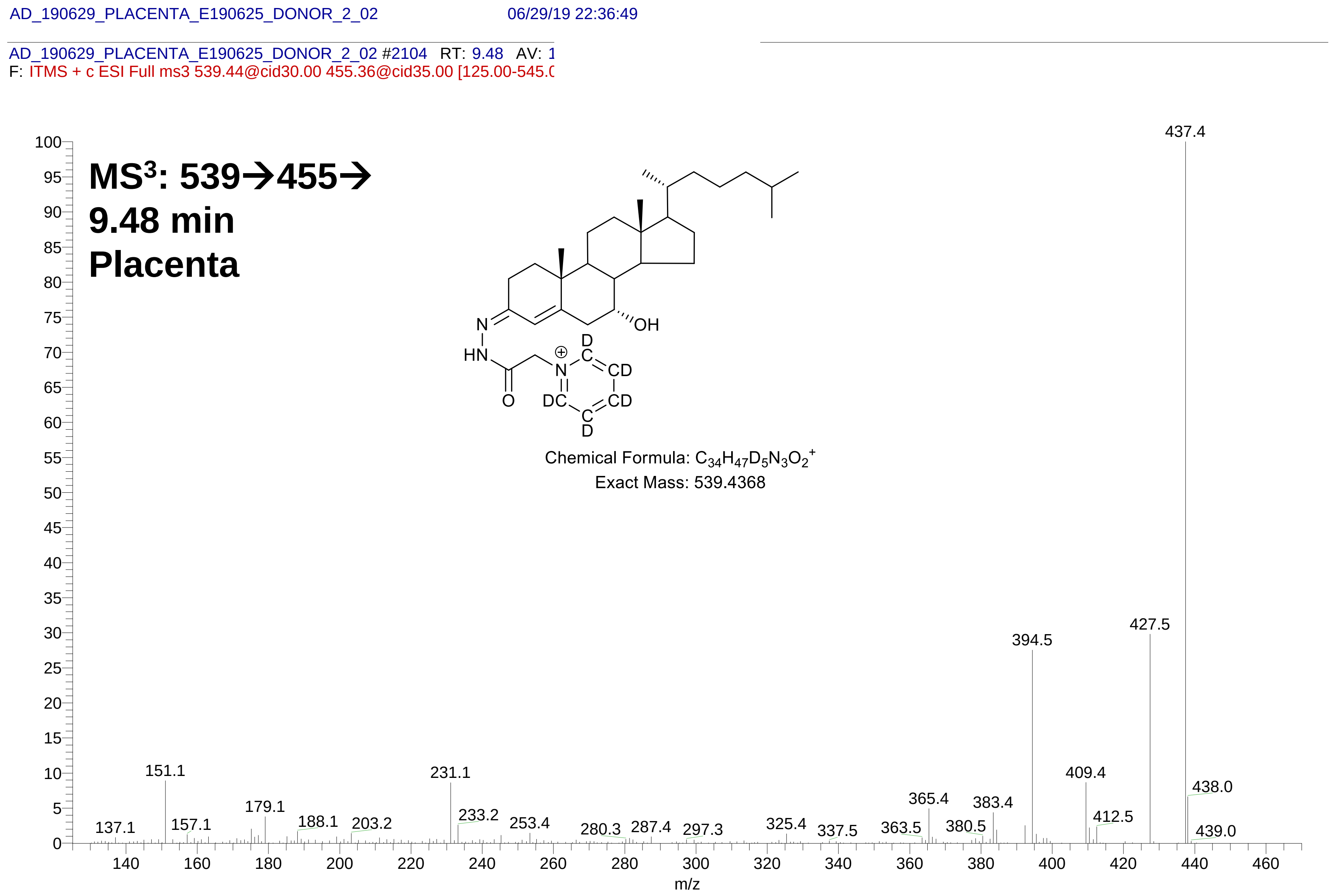
AD_190629_PLACENTA_E190625_DONOR_2_02
06/29/19 22:36:49
AD_190629_PLACENTA_E190625_DONOR_2_02 #2104 RT: 9.48 AV: 1
F: ITMS + c ESI Full ms3 539.44@cid30.00 455.36@cid35.00 [125.00-545.0
MS3: 539→455→
9.48 min
Placenta
Chemical Formula: C34H47D5N3O2+
Exact Mass: 539.4368
437.4
427.5
394.5
409.4
438.0
151.1
231.1
179.1
365.4
383.4
412.5
233.2
137.1
157.1
188.1
203.2
253.4
280.3
287.4
297.3
325.4
337.5
363.5
380.5
439.0
m/z

S4A

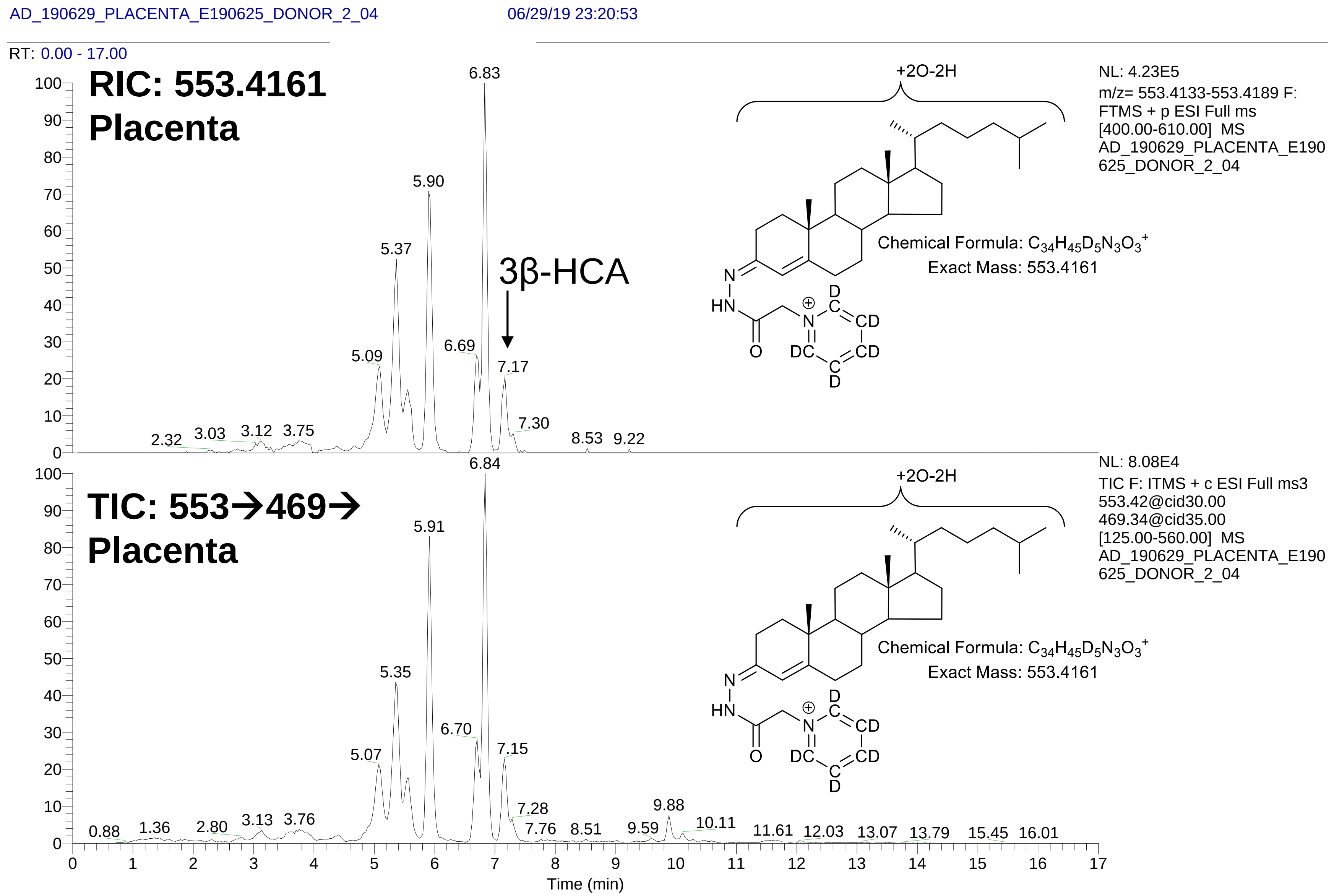

AD_190629_PLACENTA_E190625_DONOR_2_04
06/29/19 23:20:53
RT: 0.00 - 17.00
RIC: 553.4161
Placenta
3β-HCA
+2O-2H
NL: 4.23E5
m/z= 553.4133-553.4189 F: FTMS + p ESI Full ms [400.00-610.00] MS AD_190629_PLACENTA_E190625_DONOR_2_04
Chemical Formula: C34H45D5N3O3+
Exact Mass: 553.4161
TIC: 553→469→
Placenta
NL: 8.08E4
TIC F: ITMS + c ESI Full ms3 553.42@cid30.00 469.34@cid35.00 [125.00-560.00] MS AD_190629_PLACENTA_E190625_DONOR_2_04
Time (min)

**S4B**

AD_190629_PLACENTA_E190625_DONOR_2_04 06/29/19 23:20:53

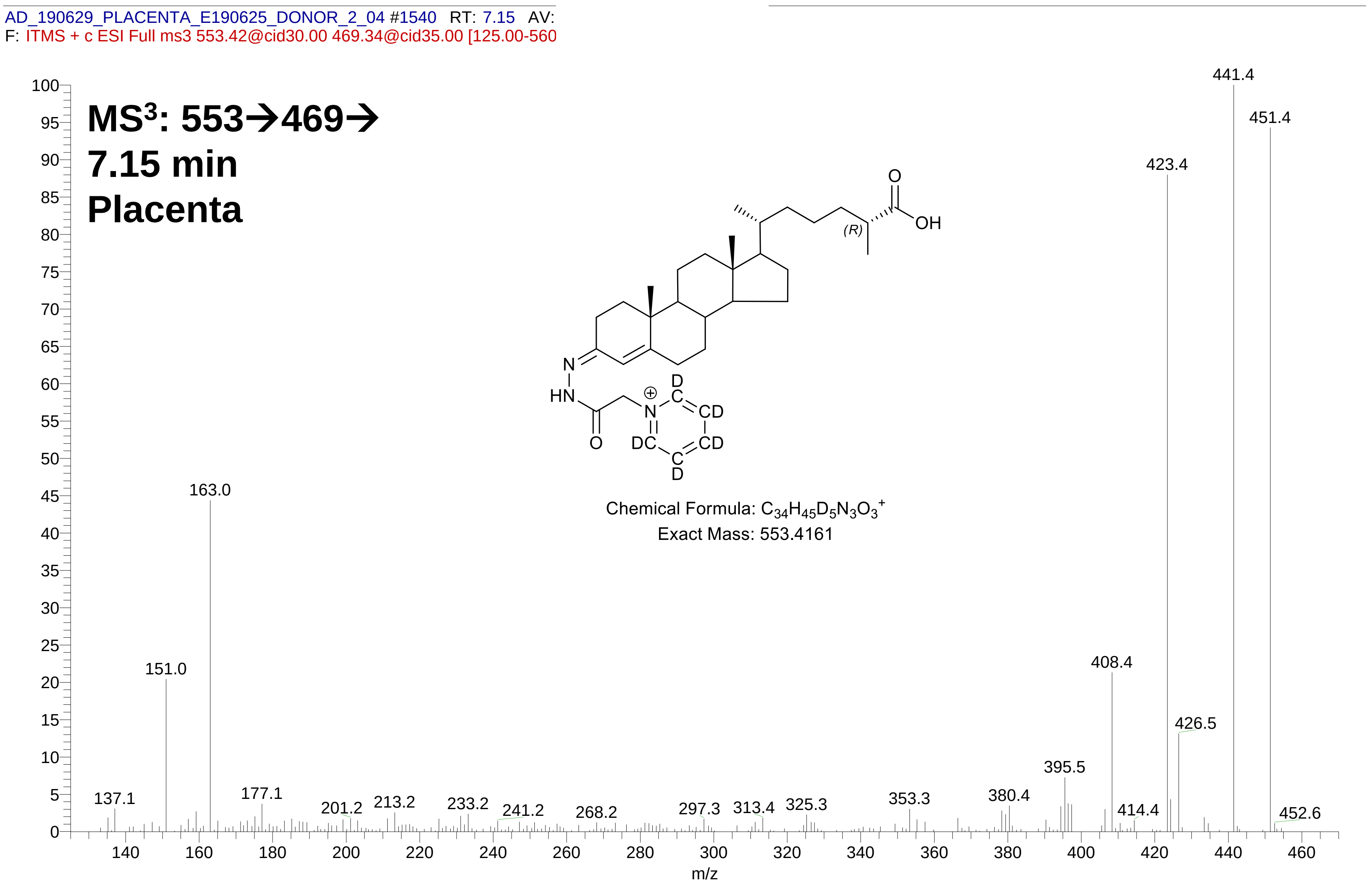

**S5A**

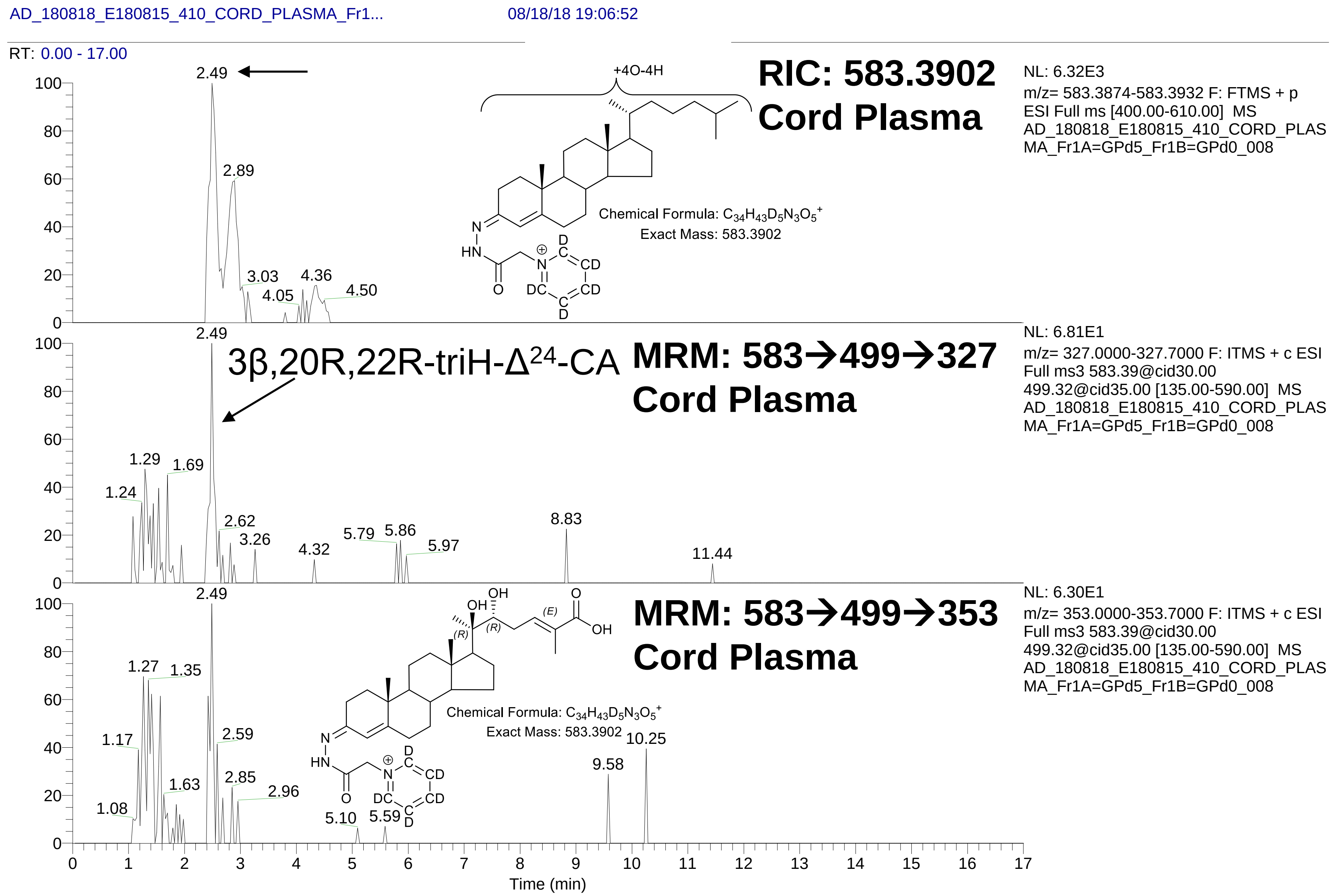

AD_180818_E180815_410_CORD_PLASMA_Fr1...
08/18/18 19:06:52
RT: 0.00 - 17.00
RIC: 583.3902
Cord Plasma
NL: 6.32E3
m/z= 583.3874-583.3932 F: FTMS + p
ESI Full ms [400.00-610.00] MS
AD_180818_E180815_410_CORD_PLASMA_Fr1A=GPd5_Fr1B=GPd0_008
+4O-4H
Chemical Formula: C34H43D5N3O5+
Exact Mass: 583.3902
2.49
2.89
3.03
4.05
4.36
4.50
MRM: 583→499→327
Cord Plasma
3β,20R,22R-triH-Δ24-CA
NL: 6.81E1
m/z= 327.0000-327.7000 F: ITMS + c ESI
Full ms3 583.39@cid30.00
499.32@cid35.00 [135.00-590.00] MS
AD_180818_E180815_410_CORD_PLASMA_Fr1A=GPd5_Fr1B=GPd0_008
1.24
1.29
1.69
2.62
3.26
4.32
5.79
5.86
5.97
8.83
11.44
MRM: 583→499→353
Cord Plasma
NL: 6.30E1
m/z= 353.0000-353.7000 F: ITMS + c ESI
Full ms3 583.39@cid30.00
499.32@cid35.00 [135.00-590.00] MS
AD_180818_E180815_410_CORD_PLASMA_Fr1A=GPd5_Fr1B=GPd0_008
Chemical Formula: C34H43D5N3O5+
Exact Mass: 583.3902
1.08
1.17
1.27
1.35
1.63
2.59
2.85
2.96
5.10
5.59
9.58
10.25
Time (min)

**S5B**

AD_180818_E180815_410_CORD_PLASMA_Fr1... 08/18/18 19:06:52

AD_180818_E180815_410_CORD_PLASMA_Fr1A=GPd5_Fr1B=GPd0_00 L: 2.87E2

F: ITMS + c ESI Full ms3 583.39@cid30.00 499.32@cid35.00 [135.00-590.(

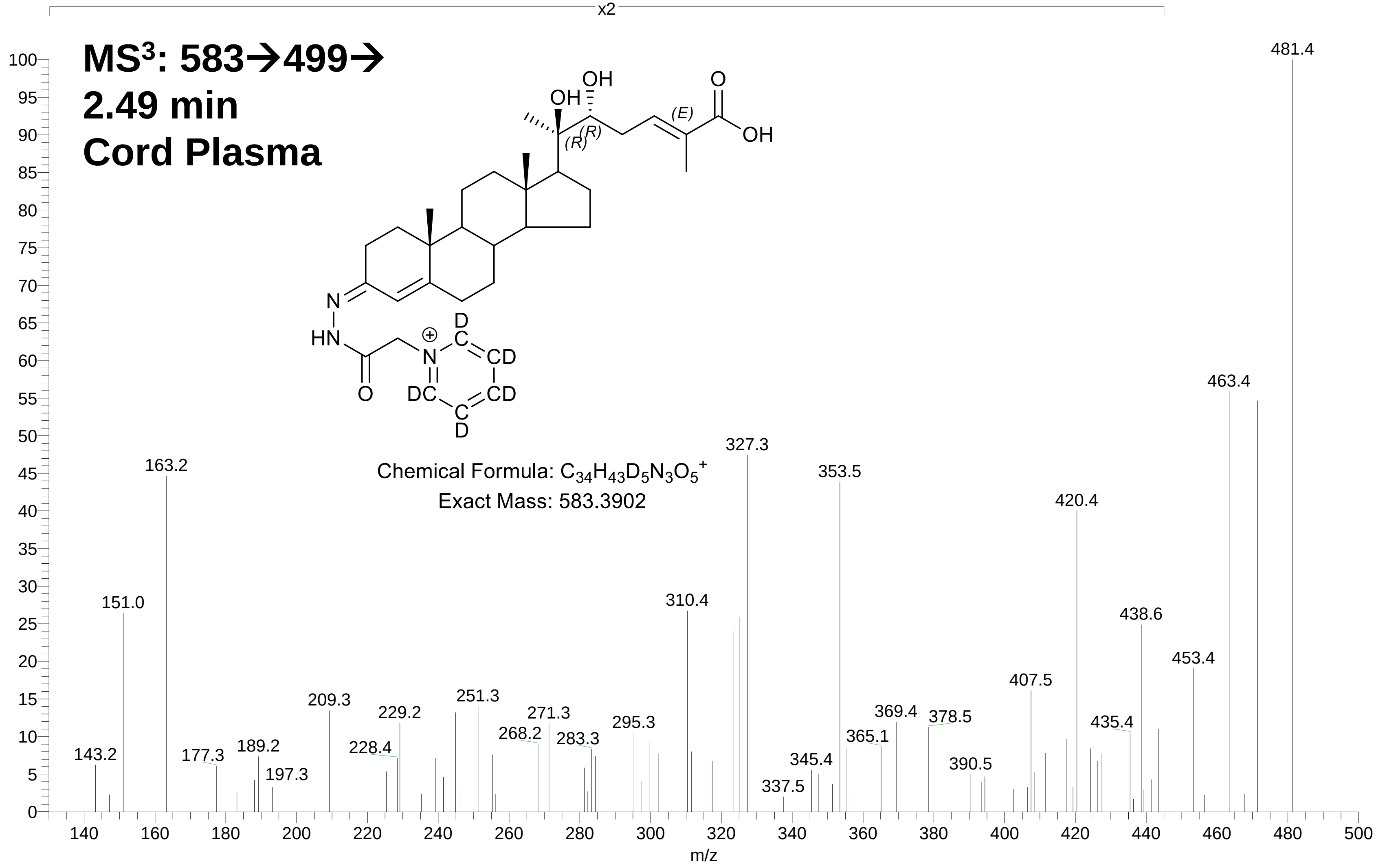

**S5C**

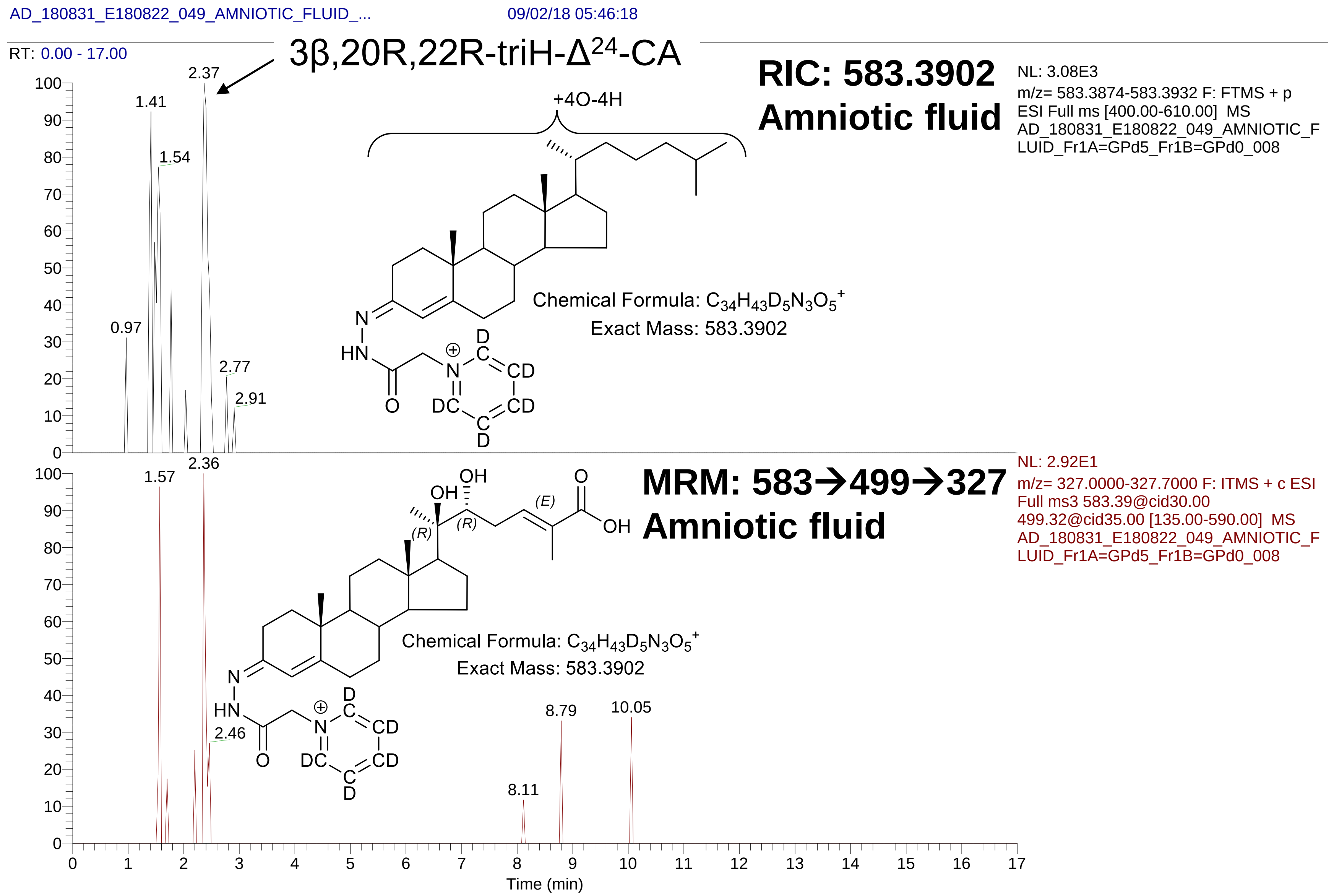

AD_180831_E180822_049_AMNIOTIC_FLUID_...
09/02/18 05:46:18
RT: 0.00 - 17.00
3β,20R,22R-triH-Δ24-CA
RIC: 583.3902
Amniotic fluid
NL: 3.08E3
m/z= 583.3874-583.3932 F: FTMS + p ESI Full ms [400.00-610.00] MS AD_180831_E180822_049_AMNIOTIC_FLUID_Fr1A=GPd5_Fr1B=GPd0_008
+4O-4H
Chemical Formula: C34H43D5N3O5+
Exact Mass: 583.3902
2.37
1.41
1.54
0.97
2.77
2.91
MRM: 583→499→327
Amniotic fluid
NL: 2.92E1
m/z= 327.0000-327.7000 F: ITMS + c ESI Full ms3 583.39@cid30.00 499.32@cid35.00 [135.00-590.00] MS AD_180831_E180822_049_AMNIOTIC_FLUID_Fr1A=GPd5_Fr1B=GPd0_008
Chemical Formula: C34H43D5N3O5+
Exact Mass: 583.3902
2.36
1.57
2.46
8.79
10.05
8.11
Time (min)

**S5D**

AD_180831_E180822_049_AMNIOTIC_FLUID_... 09/02/18 05:46:18

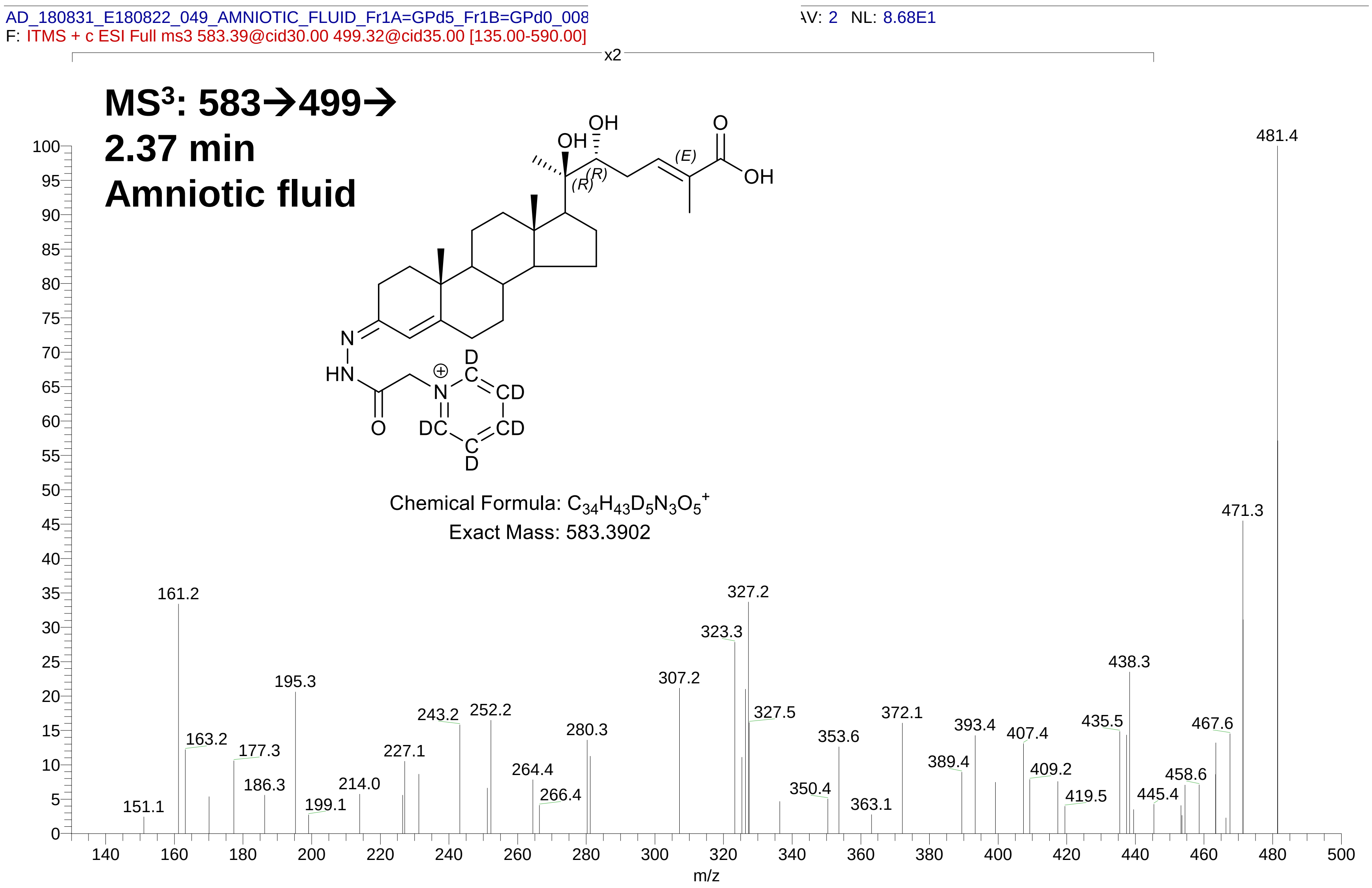
